# Nutrient colimitation is a quantitative, dynamic property of microbial populations

**DOI:** 10.1101/2023.09.27.559472

**Authors:** Noelle A. Held, Aswin Krishna, Donat Crippa, Rachana Rao Battaje, Alexander J. Devaux, Anastasia Dragan, Michael Manhart

**Affiliations:** Department of Environmental Systems Science, ETH Zurich, Zurich, Switzerland; Department of Environmental Microbiology, Swiss Federal Institute of Aquatic Science and Technology (Eawag), Dübendorf, Switzerland; Department of Biological Sciences, Marine & Environmental Biology Section, University of Southern California, Los Angeles, CA, USA; Department of Biology, ETH Zurich, Zurich Switzerland; Center for Advanced Biotechnology and Medicine, Rutgers University, Piscataway, NJ, USA; Department of Biochemistry and Molecular Biology, Robert Wood Johnson Medical School, Rutgers University, Piscataway, NJ, USA

**Keywords:** microbial growth, nutrient limitation, colimitation, *Escherichia coli*, marine microbes, biogeochemistry, biotechnology

## Abstract

Resource availability dictates how fast and how much microbial populations grow. Quantifying the relationship between microbial growth and resource concentrations makes it possible to promote, inhibit, and predict microbial activity. Microbes require many resources, including macronutrients (e.g., carbon and nitrogen), micronutrients (e.g., metals), and complex nutrients like vitamins and amino acids. When multiple resources are scarce, as occurs in nature, microbes may experience resource colimitation in which more than one resource limits growth simultaneously. Despite growing evidence for colimitation, the data is difficult to interpret and compare due to a lack of quantitative measures of colimitation and systematic tests of resource conditions. We hypothesize that microbes experience a continuum of nutrient limitation states and that nutrient colimitation is common in the laboratory and in nature. To address this, we develop a quantitative theory of resource colimitation that captures the range of possible limitation states and describes how they can change dynamically with resource conditions. We apply this approach to clonal populations of *Escherichia coli* to show that colimitation occurs in common laboratory conditions. We also show that growth rate and growth yield are colimited differently, reflecting their different underlying biology. Finally, we analyze environmental data to provide intuition for the continuum of limitation and colimitation conditions in nature. Altogether our results provide a quantitative framework for understanding and quantifying colimitation of microbes in biogeochemical, biotechnology, and human health contexts.

---

Resource availability is a fundamental control on microbial growth, physiology, and metabolic activity. Understanding which resources limit growth and by how much is therefore both a core concept of microbiology as well as useful for predicting microbial contributions to biogeochemical cycles [1], inhibiting pathogens in the human body [2], and cultivating microbes in biotechnology [3]. While limiting resources can be studied individually, there is evidence that microbes can be, and often are, simultaneously limited by multiple resources (colimitation) [4]. For one thing, the elemental composition of the environment frequently reflects that of abundant microbes, suggesting a biogeochemical balance between supply and biological demand in nature [5–9]. Furthermore, fundamental assumptions in ecology predict the possibility of colimitation; for instance, the coexistence of multiple species within a resource supply range, with each species depleting a specific resource, suggests a state of colimitation of the community as a whole [10–13]. Accordingly, in situ experiments commonly find that adding multiple resources together enhances growth of microbial communities [1, 4, 14–1 laboratory experiments have also found that microbial populations can completely deplete multiple resources simultaneously, suggesting the possibility of nutrient colimitation [3, 20].

Previous theoretical and empirical work on colimitation has largely considered limitation as a binary property of a resource (limiting or not) because of a principle known as the Law of the Minimum [3, 5, 21–24] which states that growth is solely determine by the scarcest resource relative to biological demand. As a result, empirical tests of colimitation, especially in natural samples, have primarily relied on factorial supplementation experiments, in which one measures the growth response to supplementing the population with multiple resources separately and in combination [1, 19]. However, this approach has two major shortcomings. First, the outcomes of these experiments are usually interpreted only qualitatively, mainly by classifying the outcome as single limitation or colimitation [19, 23] (see SI Appendix, Fig. S1A,B for a schematic example). This requires setting arbitrary thresholds to differentiate between these outcomes, which neglects the possibility that populations occupy a continuum of limitation states rather than discrete categories.

Furthermore, there has not been a common framework for quantifying these limitation outcomes, making them difficult to compare across studies. A similar ambiguity exists for laboratory experiments that attempt to designate a single limiting resource by keeping all other resources in excess, but without quantitatively determining what those concentrations must be to constitute single limitation. In both cases, this binary approach to limitation restricts our understanding and predictions of microbial growth in nature, where microbes experience a continuous range of resource concentrations and where many resources are simultaneously scarce [5, 8, 14, 19, 23, 25]. A second shortcoming of prior experiments, particularly supplementation experiments, is that these qualitative outcomes depend sensitively on the background resource conditions and supplemented resource concentrations (SI Appendix, Fig. S1). Besides the fact that these experiments therefore cannot tell us about the general potential for colimitation in the system beyond those specific background conditions [23], we do not know a priori how to choose the supplemented concentrations; different choices can lead to completely different outcomes, even mistaking single limitation for colimitation (SI Appendix, Fig. S1C,D). For instance, experiments that focus on extreme resource conditions, as are often used in laboratory or supplementation experiments, will miss physiological traits that emerge only under colimitation [26].

Here, we address these issues by providing a quantitative theory of resource colimitation that goes beyond the binary Law of the Minimum. We focus on independent resources, such as carbon and nitrogen sources, in clonal populations of microbes as a model case for untangling the fundamental principles of colimitation, but our results are generalizable to other types of resource combinations (substitutable or chemically-dependent [5, 23]), multispecies communities [13, 27], and higher trophic levels [19, 23, 28, 29]. We use this approach to test the hypothesis that microbes can experience a continuum of colimitation states and that these states are common enough to be observed in laboratory experiments and natural environments. We find that a wide range of laboratory conditions commonly used to study *Escherichica coli* have a significant degree of colimitation for glucose and ammonium, and that growth rate and growth yield are colimited differently. We also find that data on microbes in their natural environments is consistent with a continuum of colimitation states. This work develops a theoretical foundation for more systematically testing whether colimitation is common in nature, whether there are distinct physiological states associated with colimitation, and how resource colimitation impacts ecological, clinical, and biogeochemical outcomes.

## RESULTS

### A quantitative approach to colimitation of microbial populations

In contrast with the qualitative types of limitation (single limitation, serial limitation, additive colimitation, etc.) usually obtained in factorial supplementation experiments [5, 6, 8, 14, 19, 23, 30], we introduce a framework for precisely quantifying limitation states of populations without choosing arbitrary thresholds and categories. While in this article we focus on single-species populations of microbes, the approach can be similarly applied to study colimitation in multispecies communities [13, 27] and other organisms such as plants [19] and animals [23, 28, 29].

The per-capita growth rate *g* of a population typically depends on a large number *M* of individual limiting processes, which include uptake and utilization of different resources from the environment as well as internal cellular processes such as transcription and translation. Let *r*_*i*_ be the rate of each process *i*. We define the limitation coefficient of growth rate for process *i* as the relative change in growth rate for a small relative change in the rate of that process [3, 31]:

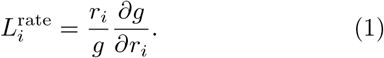

The superscript “rate” indicates that this describes limitation of the growth rate, as opposed to other growth traits as we will discuss later. For example,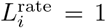 means that a 1% increase in *r*_*i*_ entails a 1% increase in growth rate, whereas 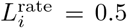 means that a 1% increase in *r*_*i*_ changes growth rate only by half as much, 0.5% (SI Appendix, section S1). For essential processes like uptake and utilization of carbon and nitrogen sources, we generally expect 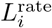 to range from zero (when *r*_*i*_ is high and the growth rate no longer depends on it) to one (when *r*_*i*_ is low and the growth rate is proportional to it). However, other values are possible if growth rate depends super-linearly or negatively on *r*_*i*_ (for example, uptake of an antibiotic).

Equation 1 is analogous with control coefficients in metabolic control analysis [32], and as in that framework, we can mathematically prove that the sum of limitation coefficients over all limiting processes must equal 1 (SI Appendix, section S2):

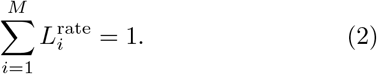

This means that greater limitation for one process/resource necessarily entails less limitation for other processes/resources. Finally, we define the effective number of growth rate-limiting processes as a metric for colimitation:

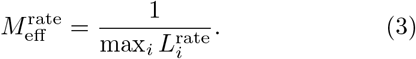

It has a minimum value of 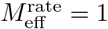 if only one resource is limiting, and a maximum value of 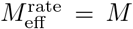(the total number of limiting processes) if all processes are equally limiting.

How does the availability of resources in the environment limit growth? The uptake rate *r* of a resource is typically assumed to be proportional to the external concentration *R* according to the law of mass action [24, 33]:

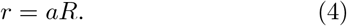

The proportionality constant *a* is sometimes known as the affinity for the resource [34]. To model the dependence of growth rate on this focal resource, we must account for the role of all other resources and internal processes that growth depends on. If we are only varying the one resource, then we can capture all these implicit limiting factors by a single rate *g*_max_, which describes the maximum growth rate when the uptake rate *r* for the focal resource is saturated; at this point, growth is limited by another process. One way to model the dependence of growth on both the focal resource and the implicit factors is to invoke the Law of the Minimum, so that the growth rate equals the rate of whichever process (uptake of the focal resource or all implicit processes combined) is slower:

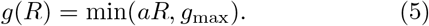

Equation 5 is sometimes known as the Blackman model [35, 36]. As shown by the solid lines in Fig. 1A, this model has binary limitation states: for resource concentrations *R < g*_max_*/a*, the limitation coefficient for the resource is 1, meaning that the implicit factors have zero limitation. For concentrations *R > g*_max_*/a*, the limitation flips and now the implicit factors are solely limiting. As a result, there can be no colimitation between the focal resource and the implicit factors. Conceptually, this situation arises if the uptake and utilization of the resource occur in parallel with the implicit factors [24, 27]. However, empirical measurements of growth rate dependence on resource concentration more commonly support the Monod model [37, 38]:

**FIG. 1.**
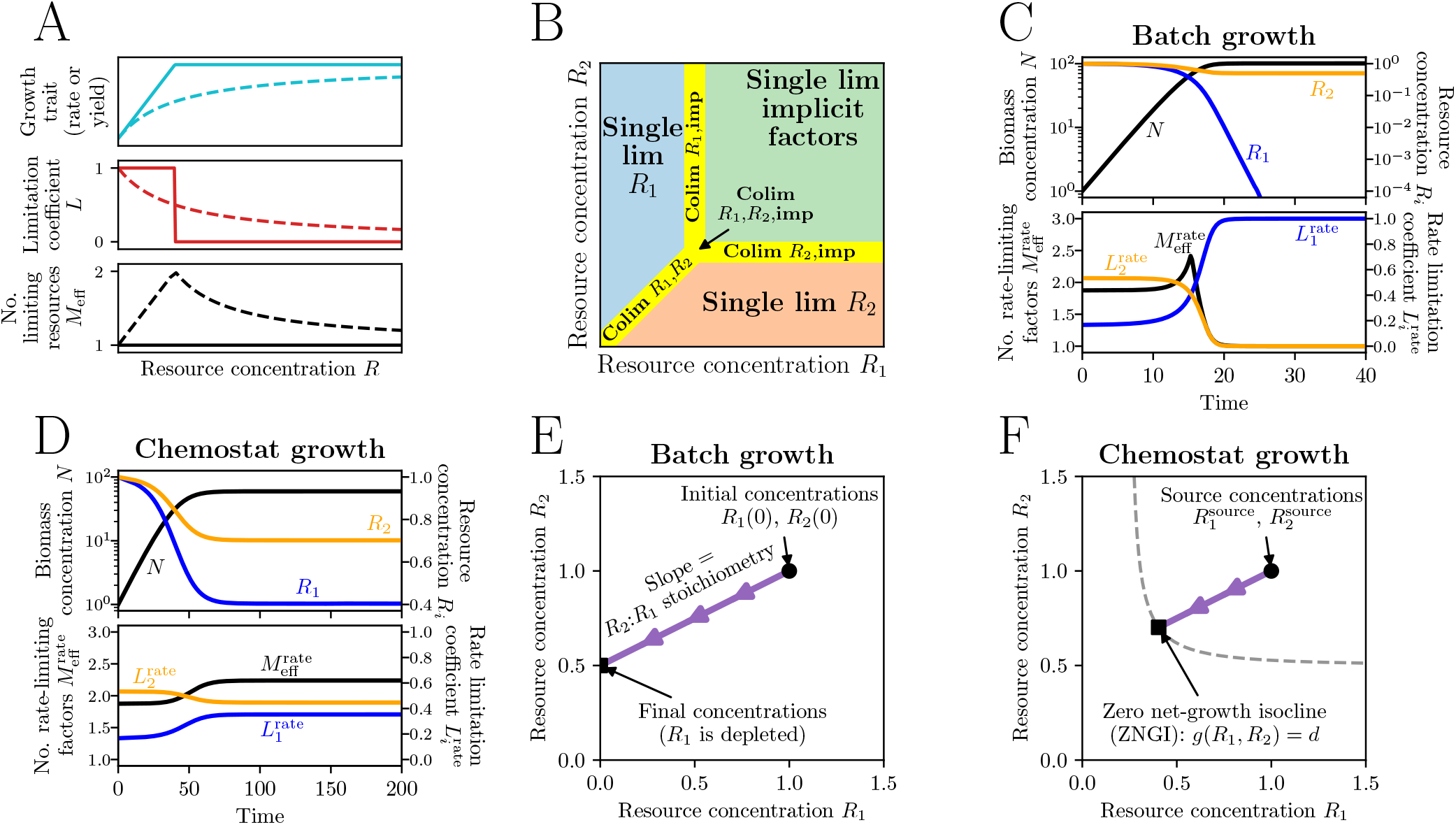
Quantifying resource limitation and colimitation of growth. (A) Schematics of how a growth trait (e.g., growth rate or growth yield) depends on a resource concentration. The solid lines show the Blackman model (Eq. 5), which does not have colimitation of the resource and the implicit factors, while the dashed lines show the Monod model (Eq. 6), which does have colimitation. (B) Schematic of limitation regimes over two resource concentrations. “Imp” denotes implicit limiting factors besides the two measured resources. (C) Simulation of batch culture dynamics and resource limitation for a population consuming two resources (blue and orange). (D) Same as (C) but for chemostat dynamics. (E) Trajectory of resource depletion for the batch dynamics in (C). (F) Same as (E) but for the chemostat dynamics in (D). Model parameters for (C)–(F) are: *g*_max_ = 1, *a*_1_ = 1, *a*_2_ = 0.5, *s*_1_ = 100, *s*_2_ = 200, *d* = 0.2 (SI Appendix, sections S3 and S4).

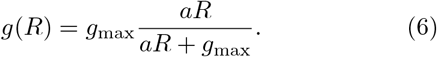

While the Monod and Blackman models have similar asymptotic behavior, the Monod model entails a smooth transition between limitation regimes, meaning that the focal resource and the implicit factors are colimiting for a finite range of concentrations around *R* = *g*_max_*/a* (Fig. 1A, dashed lines). Note that *g*_max_*/a* is also often denoted as *K*, the half-saturation concentration [38]. In particular, the Monod dependence means that limitation and colimitation take on a continuum of values, rather than being discrete states of a population as often identified in previous colimitation studies using factorial supplementation experiments. For example, Fig. 1A shows that the number of limiting factors can take intermediate values such as 1.5 if the two process have unequal but nonzero degrees of limitation (*L* = 2*/*3 for one process and *L* = 1*/*3 for another). Conceptually, colimitation of growth rate arises when resource uptake and utilization occur sequentially with the implicit processes, so that changes in either step affect the total rate [24, 27]. Note that smooth models with colimitation such as the Monod model must always have lower growth rates than do models without colimitation for the same concentration conditions, assuming the models have the same asymptotic properties.

Since microbes rely on many resources to grow, it is important to generalize this approach to considering multiple resources simultaneously. There are many models that generalize the Blackman (Eq. 5) or Monod (Eq. 6) dependence on multiple resources (SI Appendix, section S3), some of the most common being the Liebig Monod model [5, 23, 24, 27, 28, 33], the multiplicative Monod model [3, 5, 23, 27, 28, 39, 40], the Poisson arrival time/synthesizing-unit model [24, 28, 33, 41], and the additive model [28, 33]. The limitation states predicted by these models almost all follow the schematic for two explicit resources in Fig. 1B (SI Appendix, Fig. S2): there are regimes of single limitation for each of the resources as well as the implicit factors, separated by conditions of colimitation. This picture assumes both resources are independently essential and non-substitutable (e.g., a carbon source and a nitrogen source), but it can be generalized to substitutable [3, 5, 42] and chemically-dependent resources [5] as well (SI Appendix, section S3 and Fig. S3). Note that while it is common to speak of limitation for individual elements like carbon and nitrogen, in general we must specify the precise molecular forms of these elements since those forms can have distinct biological effects, especially in heterotrophs.

The quantitative nature of limitation means that it can rapidly change with the environment. For example, consider a population consuming two essential resources under batch dynamics, such that the two resources are supplied in an initial pulse (SI Appendix, section S4). Figure 1C (top panel) shows how the biomass concentration increases exponentially while the two resources are depleted. While the orange resource is more limiting initially, the blue resource becomes more limiting later before growth stops (bottom panel of Fig. 1C). The degree of colimitation, measured by the number of ratelimiting factors 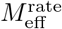, peaks at an intermediate time when the two resources have equal limitation. In Fig. 1D we show the limitation dynamics of the same system but in a chemostat, where resources are supplied to the population at constant rate (SI Appendix, section S4).

### Limitation of growth yield is distinct from growth rate limitation

Besides affecting *how fast* a population grows (growth rate), resource concentrations also determine *how much* biomass grows over a period of time, which we refer to as the growth yield. If the stoichiometry of biomass to each resource is constant (e.g., each gram of dry weight requires a fixed amount of glucose) and growth stops only when one resource concentration reaches zero, yield depends on resource concentrations according to the Law of the Minimum (similar to Eq. 5 for growth rates), since the total yield will be set by whichever resource can produce the least amount of biomass (SI Appendix, section S5). Under these conditions a single resource will therefore always be limiting, and there can be no colimitation for yield except when the resources are supplied in the exact stoichiometry that matches biological demand. For example, Fig. 1E shows the trajectory of resource depletion for the batch dynamics in Fig. 1C. The slope of the trajectory is the stoichiometry of the resources, so that constant stoichiometry means straight lines, with the length of the line being proportional to yield (SI Appendix, section S5 and Fig. S4A). Thus the yield will change if the initial concentration of resource 1 changes (which shifts the line horizontally and increases its length), while changes in the initial concentration of resource 2 will not change the yield (since that shifts the trajectory vertically and does not change its length).

However, the yield can depend smoothly on resource concentrations, and hence display colimitation, if the resource concentrations at which growth stops depend on each other (SI Appendix, section S5 and Figs. S4B and S5A,B), or if the biomass stoichiometry of one resource depends on the concentration of another resource (SI Appendix, Fig. S4C and S5C–F). The first case holds for chemostat dynamics if the zero net-growth isocline [42] (ZNGI, where growth rate equals the dilution rate; dashed lines in Fig. 1F) is curved; even with fixed stoichiometry so that depletion trajectories are straight lines, their length to the ZNGI (which is proportional to the yield) will depend on the supplied concentration of both resources (Fig. 1F; SI Appendix, section S5 and Fig. S5A,B). Variable stoichiometry, on the other hand, is possible through a variety of mechanisms, including resource consumption for biomass maintenance (SI Appendix, section S5 and Fig. S5C,D) and proteome allocation that varies with growth rate (SI Appendix, section S5 and Fig. S5E,F).

*i*We quantify growth yield limitation analogously with growth rate limitation, defining yield limitation coefficients 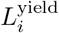 (Eq. 1 but replacing growth rate with growth yield) and the number of yield-limiting resources 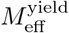 (Eq. 3 but with 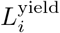 instead of 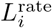. We can also use equivalent mathematical models for describing how growth yield depends on resource concentrations (SI Appendix, section S3), leading to similar possibilities for colimitation (Fig. 1A,B). However, growth rate and growth yield limitation are biologically distinct aspects of populations. In particular, resources may limit rate and yield differently — for example, in Fig. 1C the orange resource limits growth rate initially but the blue resource limits growth yield since it is depleted at the end. That being said, rate and yield limitation are coupled under certain circumstances. In batch dynamics, the yield-limiting resource (the one depleted when growth stops) is also necessarily the most rate-limiting resource at the end of growth. Under chemostat dynamics, growth rate limitation determines growth yield limitation because the former sets the shape of the ZNGI [13, 42]. For example, if two resources obey a model without growth rate colimitation, then the ZNGI forms a right angle, in which case there is also no yield colimitation [27].

### Growth rate and yield of *E. coli* are colimiting for glucose and ammonium under laboratory conditions

To demonstrate our quantitative approach to colimitation, we measure the dependence of *Escherichia coli* growth rate and yield on glucose and ammonium, two essential resources, under laboratory conditions. We also use these empirical measurements to address the second major shortcoming of traditional colimitation tests: instead of using only a single background condition and supplemented concentration for each resource, which cannot conclusively determine the extent of colimitation in a system [23] (SI Appendix, Fig. S1), we systematically scan a wide range of glucose and ammonium concentrations to determine their global potential for colimitation (Materials and Methods; Datasets S1 and S2; SI Appendix, Figs. S6–S10)

As expected, the data shows that the growth rate depends on glucose and ammonium concentrations when they are low, but once they are sufficiently high, growth rate saturates as other implicit factors becomes limiting (Fig. 2A,B). To quantify colimitation in this data, we first fit the data to a range of models both with and without colimitation (SI Appendix, Table I and Figs. S11– S13). We find that while the fits favor models with colimitation (Poisson arrival time/synthesizing-unit model; SI Appendix, section S3) for two of the three experimental replicates, this result is not conclusive as the models without colimitation also fit the data fairly well (SI Appendix, Fig. S13).

**FIG. 2.**
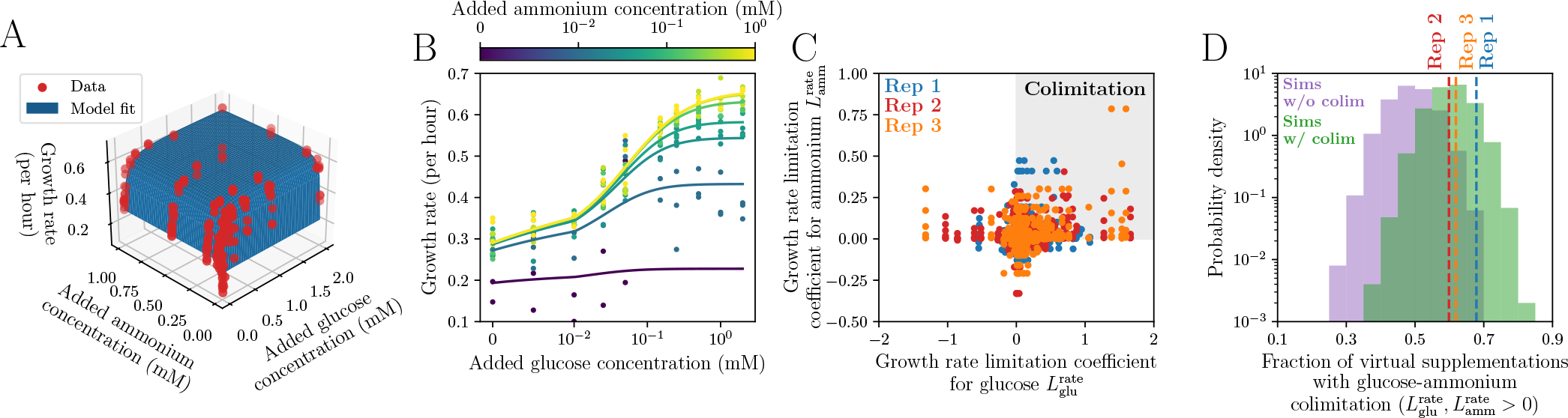
Measuring growth rate colimitation in laboratory conditions. (A) Three replicate measurements of growth rate (red points) over glucose and ammonium concentrations for *E. coli* in minimal medium (Materials and Methods, Dataset S1; see SI Appendix, Figs. S6–S8 for growth curves). The blue surface is a model fit to all replicate measurements (Poisson arrival time/synthesizing-unit model with *g*_max_*≈* 0.66 per hour, *a*_glu_*≈* 23 per hour per mM glucose, *a*_amm_*≈* 90 per hour per mM ammonium; *R*_glu,min_*≈* 0.023 mM, *R*_amm,min_*≈* 0.0039 mM; Materials and Methods; Dataset S1). (B) Same growth rate data as in (A) but plotted as a function of glucose concentration alone, with colors indicating ammonium concentrations. The lines are the same fitted model as in (A). (C) Growth rate limitation coefficients for glucose and ammonium from virtual supplementation experiments inferred from data in (A); the three replicate experiments are shown as different colors. The gray box marks the region of points with apparent colimitation of glucose and ammonium 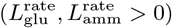. (D) Fractions of virtual supplementation experiments with growth rate colimitation between glucose and ammonium. The observed fractions from the three replicate experiments are marked with dashed lines of different colors. Compared to a null model without glucoseammonium colimitation (purple, 10^4^ simulations of the Liebig Monod model; SI Appendix, section S3), the probabilities of observing at least as much colimitation as in the data are *p* = 0.0004 for replicate 1, *p* = 0.0338 for replicate 2, and *p* = 0.0135 for replicate 3. Simulated data sets from a model with glucose-ammonium colimitation (green, Poisson arrival time/synthesizingunit model; SI Appendix, section S3) show frequencies of colimitation more consistent with the data. The simulations use parameters from the fits of these models to the experimental data and random noise estimated from the variation across experimental replicates (Materials and Methods).

However, these fits are based on an optimization that weighs all data points equally, which may not be sufficiently sensitive to differences in the subset of concentrations where there is approximate balance of limitation between resources and colimitation can occur. To better detect colimitation under the conditions that matter, we therefore leverage the combinatorial completeness of concentration conditions in our data set to perform “virtual” supplementation experiments, in which we take each concentration condition from our data and calculate the glucose and ammonium limitation coefficients for all other concentrations relative to that background (Materials and Methods). This direct-from-data approach is independent of the fits as the limitation coefficients are estimated as finite differences in growth rates and resource concentrations. Figure 2C shows these limitation coefficients for all virtual experiments across all three replicate data sets.

In principle, any points where 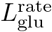 and 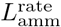 are both greater than zero indicate colimitation (Fig. 2C, gray box; SI Appendix, Fig. S14A). However, measurement noise could cause such observations to randomly appear even in the absence of true colimitation (SI Appendix, Fig. S14B). To determine whether the amount of observed glucose-ammonium colimitation is greater than would be expected from noise, we simulate a large number of data sets using models with and without colimitation but based on parameters and measurement noise fit to the data (Materials and Methods; SI Appendix, Figs. S15–S18). For each simulated data set, we calculate the fraction of virtual supplementation experiments with glucose-ammonium colimitation. Compared to 10^4^ simulations of a null model without glucose-ammonium colimitation (purple distribution in Fig. 2D; SI Appendix, Fig. S19), we find that the probabilities of generating colimitation levels at least as high as in each of our experimental replicates (*p*-values) are 0.0004, 0.0338, and 0.0135. In contrast, a null model that does have glucose-ammonium colimitation is largely consistent with the data (green distribution in Fig. 2D). This data therefore supports the existence of growth rate colimitation for *E. coli* under laboratory conditions. We perform an analogous analysis on our growth yield data and obtain similar results demonstrating the occurrence of yield colimitation for glucose and ammonium (SI Appendix, Figs. S20–S

### Comparison of growth rate and growth yield limitation in *E. coli*

Given the evidence of colimitation in our data, we can use the fitted colimitation models to quantitatively map different limitation states across resource concentrations. This is important for performing comparative measurements of cell physiology, ecology, or evolution across resource conditions; instead of simply postulating that a condition with low glucose and high ammonium is carbon-limiting, for example, we can precisely define *how* limiting that state really is. Figure 3A shows limitation coefficients of glucose and ammonium for growth rate (top panel) and growth yield (bottom panel) across concentrations. We find that *E. coli* in typical M9 medium (0.2%*≈* 11.1 mM glucose, 18.7 mM ammonium) has growth rate limitation coefficients of 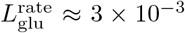 and 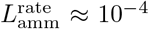this means that there is approximately single limitation for the implicit Factors 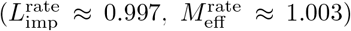. On the other hand, the limitation coefficients for growth yield in these conditions are 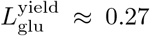 and 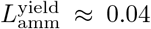, which means there is an appreciable amount of colimitation, especially between glucose and the implicit yield-limiting factors 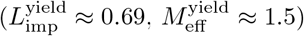.

**FIG. 3.**
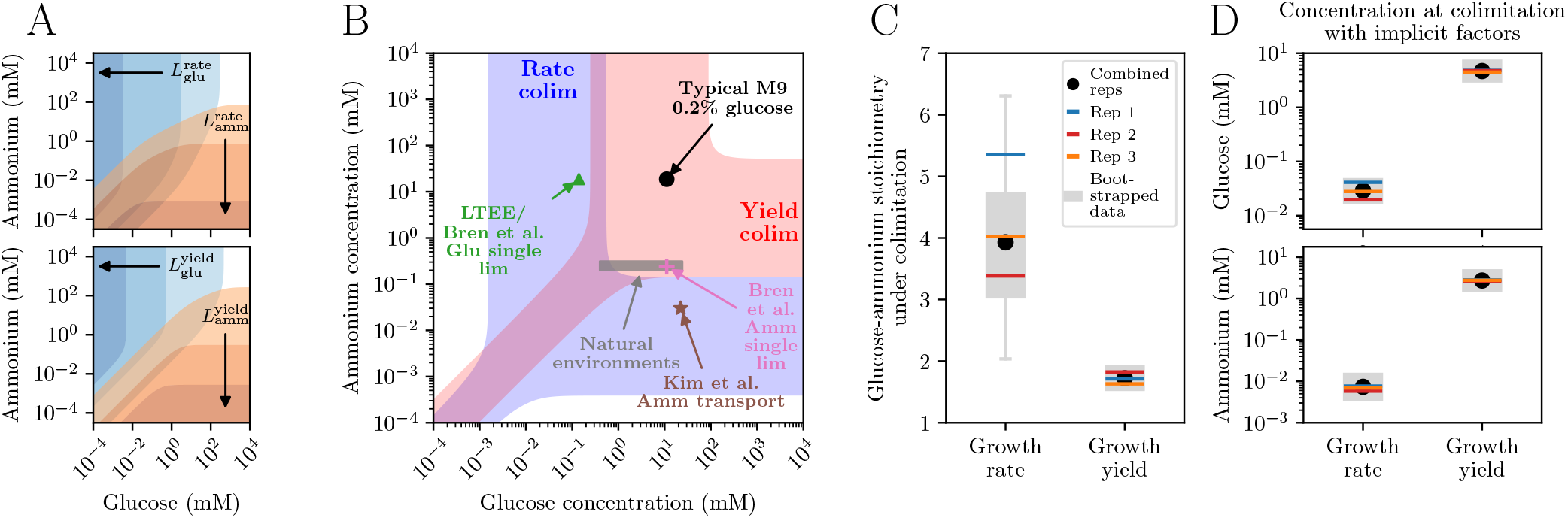
Comparison of growth rate and growth yield colimitation in *E. coli*. (A) Using models fitted to measurements of growth rate (top) and growth yield (bottom), contours of limitation coefficients for glucose (blue) and ammonium (orange). From lightest to darkest, the contours show limitation coefficients above 10^−4^, 0.01, and 0.9 for growth rate (top), and above 0.01, 0.9, and 0.999 for growth yield (bottom). (B) Phase diagram of rate and yield colimitation in *E. coli* across glucose and ammonium concentrations in M9 medium. The blue shaded region shows concentrations of glucose and ammonium with significant colimitation, where no one factor is limiting more than 95% of growth rate 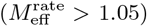. The red shaded region is the analogous concentration range for growth yield colimitation. Symbols mark reference concentrations of glucose and ammonium used in other studies: typical M9 media (0.2%*≈* 11.1 mM glucose, 18.7 mM ammonium); putative single limitation for glucose (0.14 mM glucose, 18.7 mM ammonium) used by Bren et al. [43], which is very similar to the concentrations used in the Long-Term Evolution Experiment [44] (but with*≈* 15.2 mM ammonium in Davis-Mingioli media); putative single limitation for ammonium (11.1 mM glucose, 0.24 mM ammonium) used by Bren et al. [43]; conditions at which Kim et al. found that *E. coli* activates its ammonium transporter [45]; and the estimated range of natural environments for *E. coli* (Dataset S3). (C) Stoichiometry of glucose to ammonium under growth rate and growth yield colimitation. This ratio is calculated as the ratio of ammonium to glucose affinities fitted by the models to the data (SI Appendix, section S3). The black point shows the stoichiometry from the fit to all three experimental replicates combined, while the colored bars represent the fits to each replicate individually. The gray boxes (first to third quartiles, with the whiskers extending to 1.5 times the interquartile range above and below) show the distributions of fitted stoichiometry across 100 data sets bootstrapped from the three replicates (Materials and Methods). (D) Same as (C) but for the concentrations of glucose (top panel) and ammonium (bottom panel) at which they are colimiting with the implicit limiting factors.

Figure 3B compares growth rate and growth yield colimitation across a broad range of glucose and ammonium concentrations (compare to Fig. 1B; SI Appendix, Fig. S30). The blue region marks conditions of significant colimitation (where no single resource has more than 95% of total limitation for growth rate), while the red region marks the analogous conditions for growth yield. Together they demonstrate that a wide range of conditions — from approximately 1 µM to 100 mM of either glucose or ammonium — have some degree of colimitation for either growth rate or growth yield. We also find that the conditions of glucose and ammonium colimitation for growth rate overlap significantly with those for growth yield, but growth rate and growth yield have largely non-overlapping conditions for colimitation between the implicit limiting factors and either glucose or ammonium. Besides typical M9 media, we also compare our colimitation map to several other reference media conditions, including the Long-Term Evolution Experiment [44] (green triangle, with the caveat that that experiment uses Davis-Mingioli rather than M9 media and a different strain of *E. coli*); conditions for putative single limitation for glucose and for ammonium used in a study by Bren et al. [43] (green triangle and pink cross); conditions at which *E. coli* activates its ammonium transporter [45] (brown star); and the estimated range of glucose and ammonium in *E. coli* ‘s natural environment (gray box; Materials and Methods; Dataset S3).

It is also valuable to quantitatively compare the colimitation conditions between growth rate and growth yield. We find that the glucose-ammonium stoichiometry needed for growth rate colimitation is approximately 4 (C:N = 24), while it is approximately 1.7 (C:N = 10.2) for growth yield colimitation (Fig. 3C). Both values suggest that the environment under these apparently balanced conditions must have a C:N ratio much higher than the typically observed *E. coli* biomass C:N ratio [46] (4.33–5.17 depending on the growth conditions). The conditions for colimitation between implicit factors and glucose or ammonium are also quite different between growth rate and growth yield, with the concentrations being approximately 100-fold higher for growth yield colimitation than for growth rate colimitation (Fig. 3D). This suggests that the implicit limiting factors for growth rate and growth yield are different, which is not surprising, but this may also be indicative of the different selection pressures on rate versus yield limitation. For example, stronger selection on rate limitation compared to yield limitation may have driven rate limitation toward much lower concentrations.

### Quantifying colimitation of growth rate and growth yield in natural environments

While we have shown that colimitation can occur in the laboratory, an important but more complex problem is to test whether colimitation occurs for microbes in their natural environments [4]. We now demonstrate our quantitative approach by estimating limitation and colimitation in several natural ecosystems. To quantify colimitation of growth rate, we collected measurements of half-saturation resource concentrations *K*_*i*_ (measured from the dependence of growth rate on that resource concentration according to the Monod model [38]) and environmental concentrations *R*_*i*_ for those same resources across many organisms (Fig. 4A; Materials and Methods; Dataset S3). Note that some strains with laboratory measurements are different from those in the natural environments and hence may have different trait values [38].

**FIG. 4.**
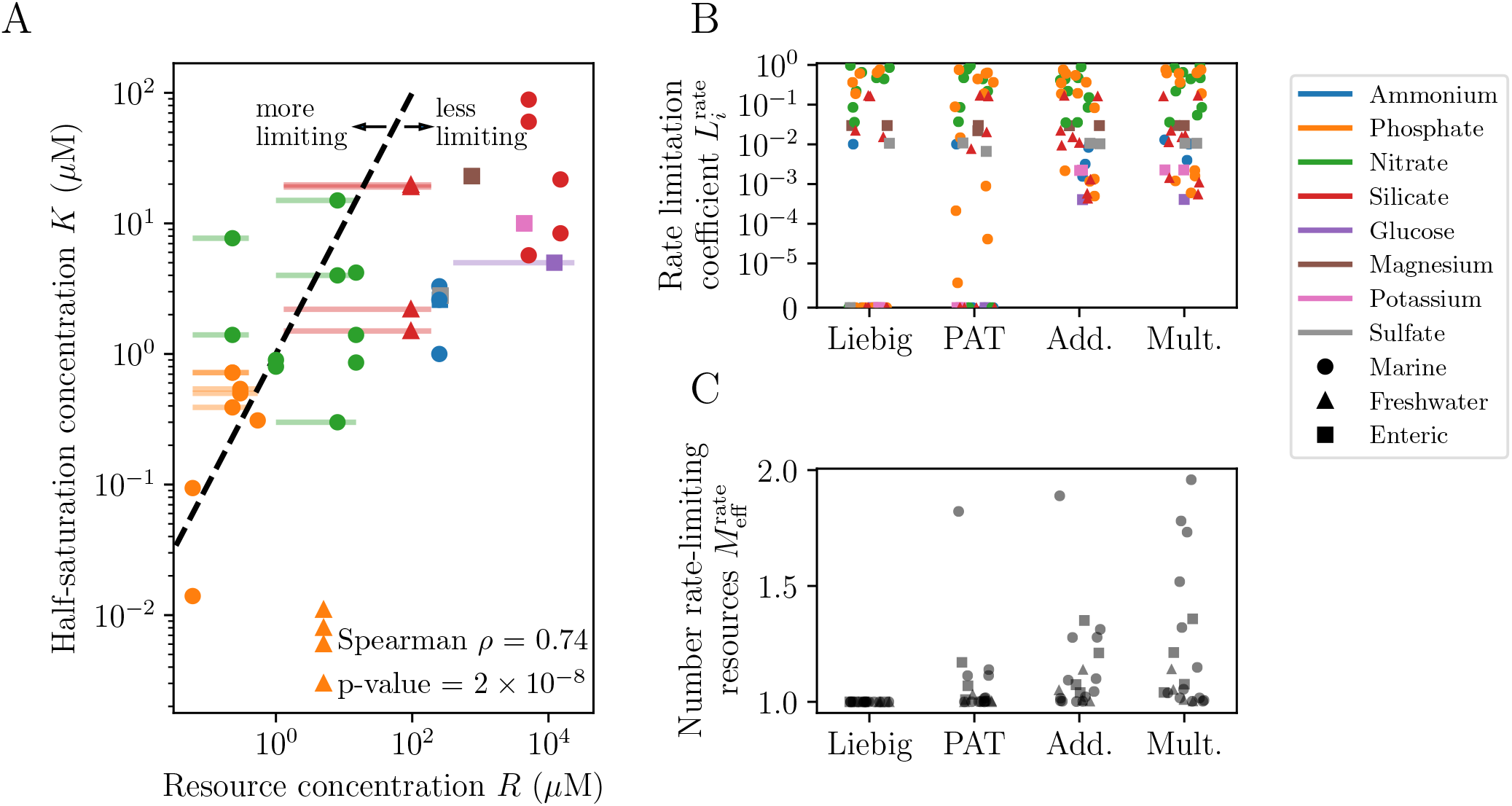
Applying limitation concepts to empirical growth rate data across species and resources. (A) Halfsaturation concentrations *K* versus environmental concentrations *R* from various organisms, resources, and environments collected from the literature (Materials and Methods; Dataset S3). When we found environmental concentrations *R* as ranges, we indicate this in the plot by horizontal bars attached to the symbol. Symbol color indicates resource, and shape indicates habitat. The Spearman correlation coefficient between *R* and *K* is *ρ≈* 0.74 with a *p*-value*≈* 2*×* 10^−8^. (B) Rate limitation coefficients 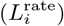 calculated from the data shown in (A); we calculated limitation coefficients for individual resources in all organism/resource combinations, considering four common growth rate models (horizontal axis: Liebig = Liebig Monod model, PAT = Poisson arrival time/synthesizing-unit model, Add. = additive model, Mult. = multiplicative Monod model; SI Appendix, Table I and section S3). (C) Number of rate-limiting resources *M* ^rate^ among the resources in the data shown in (A), also considering the same growth rate models as in (B). Each organism has data for a pair of resources, such that the maximum number of limiting resources is 2.

We use this data to estimate limitation for each resource, assuming all other resources for these organisms are present at very high concentrations. Since the true dependence of growth rate on multiple resources is unknown for these organisms and resources, we consider four common growth rate models (Liebig Monod, multiplicative Monod, Poisson arrival time/synthesizing unit, and additive; SI Appendix, Table I and section S3) to calculate limitation coefficients in Fig. 4B. The analysis reveals the necessity of our quantitative approach: the data shows a continuum of limitation coefficients, rather than a dichotomy of high and low limitation (SI Appendix, Fig. S31). In particular, we find significant intermediate limitation coefficients for many of the marine species, but low limitation for the freshwater and enteric species. This is evident from the *R* and *K* data itself; most half-saturation concentrations *K* are less than their corresponding environmental concentrations *R*, which means low limitation, with the exceptions being several marine strains on phosphate and nitrate (Fig. 4A). The lack of large growth rate limitation coefficients is not surprising given the strong selection to reduce the half-saturation concentration relative to the environment [38]. We moreover see a continuum of growth rate colimitation states between these resources according to the number of rate-limiting resources 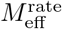 (Fig. 4C; SI Appendix, Fig. S31). While several of the marine species show high colimitation, we see intermediate levels of colimitation for some enteric species as well (Fig. 4C).

To demonstrate quantification of colimitation for growth yield, we apply our approach to elemental resources for marine phytoplankton. Using the abundance of each element in the ocean and its stoichiometry in phytoplankton biomass [6], we calculate the maximum potential growth yield of each element (Fig. 5A); this is the maximum amount of biomass that could grow from that element’s concentration in the ocean, if all other elements were unlimited. This shows roughly three tiers of resources, separated by orders of magnitude: the trace metals iron, cobalt, and manganese have the lowest potential yields, followed by another set of similar potential yields for nitrogen, phosphorus, and other metals. The remaining elements all have much higher potential yields.

**FIG. 5.**
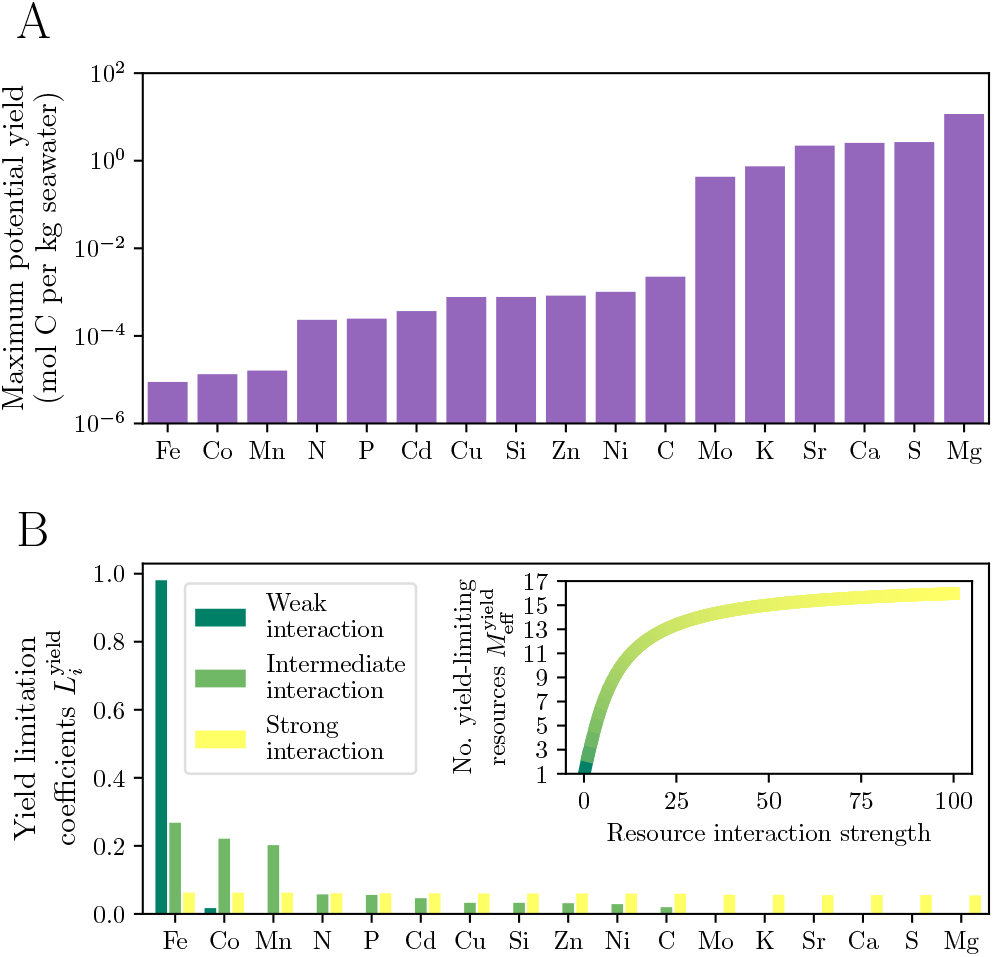
Applying limitation concepts to empirical growth yield data for marine phytoplankton. (A) Maximum potential growth yield for phytoplankton (measured in moles of carbon) per kg of seawater that can be produced by each elemental resource’s environmental concentration. Data is from Moore et al. [6] (Supplementary Table S1 therein). (B) Using potential yields from (A), we calculate growth yield limitation coefficients 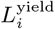 for each elemental resource. Since we do not know the underlying model for how multiple resources contribute to growth yield, we calculate limitation using a generalized-mean model with variable strength of resource interaction that tunes the degree of colimitation (SI Appendix, section S3). Inset: Number of yield-limiting resources 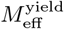 as a function of the resource interaction strength, using data from (A).

From these potential yields we can calculate yield limitation coefficients for each elemental resource. Since we do not know how phytoplankton growth yield depends on multiple resources, we use a phenomenological model based on the generalized mean, with a variable parameter that captures interactions between resources in determining the yield (SI Appendix, section S3). For weak interactions between resources, limitation is strongly concentrated in the resource most rare relative to its biomass requirements, especially iron (Fig. 5B), and hence there is little colimitation (Fig. 5B, inset). For stronger resource interactions, we see greater colimitation for yield across resources.

This analysis reflects known patterns of oceanic nutrient limitation, such as the prevalence of iron limitation [1, 6] as well as nitrogen and phosphorus limitation, though we note that it is interesting that N and P are only appreciably limiting in the strong interaction scenarios. While the data we used is coarse-grained and does not account for temporal, spatial, and biological variability (including plasticity in resource use), nor the full suite of nutrients that can potentially limit phytoplankton, it does indicate that to determine the true level of colimitation for these populations, we would need to measure the dependence of phytoplankton biomass across many resources simultaneously.

## DISCUSSION

### What are the consequences of colimitation?

Our results demonstrate that resource colimitation of microbes is possible and indeed plausible. The consequences of colimitation span individual cells, populations, communities, and ecosystems. For one thing, colimitation has been known to alter traits such as cell size [26], meaning it is possible to develop misleading ideas about physiology when the focus is only on single, strongly-limiting resources. This problem has been described in the literature since the 1970s, notably in work by Droop [47] and by Valdkamp and Jannasch [48], who pointed out that laboratory experiments that provide artificially high concentrations of resources besides a focal limiting resource do not reflect the realities of nature. Adding to this, we point out that growth is less efficient under colimitation models (e.g., Monod) versus single limitation models (e.g., Blackman), suggesting a fundamental physiological difference in single limitation versus colimitation.

Colimitation seems to be consequential for ecoevolutionary dynamics in microbial populations and communities. For one thing, resource colimitation has been linked to distinct evolutionary outcomes that are different from single nutrient limitations [49]. In microbial communities, colimitation could increase susceptibility to invasion, because there are multiple potential niches for an invader to exploit. The competitive exclusion principle says that the number of species cannot exceed the number of limiting resources. However, when “limiting” is no longer a binary state of a resource, the concept must be reexamined. Consideration of colimitation could therefore enhance the ability to predict and engineer microbial communities in environmental and human health contexts. For instance, knowing which nutrients are limiting and by how much would be useful in the design of probiotics and prebiotics.

Lastly, there is evidence that colimitation plays a role at larger ecological and biogeochemical scales. In a broad sense, ecological and biogeochemical theory indicates that the requirements of microbial life and the elemental composition of their surroundings coevolve with one another, and this could increase the possibility for resource colimitation in nature [50, 51]. Accordingly, factorial supplementation experiments often find colimitation and collectively demonstrate that colimitation is a rather common state in systems such as the surface ocean [1, 5]. Given this evidence, resource colimitation should be considered when predicting microbial contributions to ecological processes. So far, the impact of colimitation on microbial activity has been described mainly qualitatively, however describing colimitation quantitatively as we have done here will facilitate comparisons across field experiments and data and hopefully broader examination of this topic in biogeochemical and ecological models.

This raises the question of which model to use to describe nutrient colimitation. For individual resources, the biggest distinction is between models that vary smoothly with resource concentration (such as the Monod or Droop models), which allow for colimitation between the resource and a second, implicit process, versus models such as in Blackman limitation, which entail abrupt transitions between limitation regimes. While our data supports the use of smooth models, it is difficult to distinguish among them. Similarly, for multiple resources, there is a distinction between Law of the Minimum functions and colimitation models such as multiplicative model, Poisson arrival time (PAT) model, and additive models. However, it is again difficult to distinguish between these models by fitting to isolated experimental data [24, 28]. This raises the question of which formalism to use for describing microbial growth in nutrient limitation and colimitation. On one hand, the chosen formalism may not matter for broad ecosystem-level outcomes such as stability of a microbial community [52], so long as nutrient colimitation is a possibility in that model. On the other hand, there are probably situations in which the chosen formalism will make a big difference, such as the outcome of an evolution process, or rate of a biogeochemical process. The sensitivity of Earth System, clinical, and biotechnology outcomes to nutrient colimitation is an important area for future research.

### What mechanisms might cause colimitation?

Intuitively, resource colimitation implies that there is an alignment of environmental availability and biological need [4]. This in turn suggests that there are interactions among resources leading to microbial growth. The most direct examples include biochemically-dependent resource colimitation and substitutable resource colimitation, however indirect mechanisms are also possible including tradeoffs due to constraints on molecular composition, resource uptake, and energy utilization that is exacerbated when multiple resources are rare. Systems biologists have predicted the possibility or resource colimitation in laboratory systems; in individuals or populations, this results from fine-scale molecular networks and especially stoichiometric flexibility, which could be a direct result of growth optimization in resource-colimited conditions [39, 53]. While variable stoichiometry is thought to be most common in autotrophic organisms such as phytoplankton, it has also been observed and linked to resource colimitation in heterotrophic consumers [23].

Colimitation also results from heterogeneity, both biological and environmental. In microbial communities, colimitation could result from heterogeneity in resource preferences and biomass stoichiometries, such that different organisms are limited for different resources resulting in a community-level colimitation [13]. This mechanism could be an indirect result of community formation, or it could involve direct metabolic dependencies among organisms [12]. These community-level effects include changes in community composition and processing of resources through biological pools would be layered on the backdrop of colimitation mechansisms for individuals and populations. Additionally, in clonal populations, similar biological heterogeneity could occur if there is phenotypic heterogeneity [54], for instance if a subset of the population is primarily limited for one resource while another subset is limited for another resource. Lastly, environmental heterogeneity is also a factor, as it has been suggested that microscale environmental heterogeneity could reduce the chance that cells encounter both necessary resources quickly; this is implicit in the Poisson arrival time (PAT) model (SI Appendix, section S3). Because our quantitative framework is agnostic to the mechanism of the colimitation, it can be applied to all of these scenarios.

The observation of heterogeneity naturally leads to questions of stochasticity. While stochasticity is playing a role in all biological and environmental processes, our ability to replicate and witness two-dimensional trends in growth indicate that pure randomness is not a major contributor. Rather, nutrient colimitation emerges systematically. This in turn raises the question of whether the cause of nutrient colimitation is similar across organisms and resource pairs. If generalizable mechanisms can be identified in microbes, it is then possible that these mechanisms could be extended to organisms of higher orders. Supporting the idea of generalizable mechanisms, recent work has suggested that different models of colimitation are different approximations of the same underlying process [24, 27].

### Are microbes actually colimited in nature?

The fact that we find evidence of colimitation for two essential, independent resources in the model organism *E. coli*, and in particular in resource concentrations that are relevant to *E. coli* ‘s natural environments, highlights a need for closer attention to resource colimitation in nature. Prior studies have also investigated resource colimitation of *E. coli*, but used a chemical definition, i.e., whether multiple resources were simultaneously depleted to zero [20, 55, 56]. In contrast, we suggest a biological definition (how is growth affected by the resources?) because 1) this allows precise identification of resource colimitation for different growth, activity, or physiological parameters of interest; 2) this strategy can be used in either batch or chemostat cultures, meaning also that growth rate and growth yield colimitation can be quantified separately; and 3) this allows instantaneous definitions of resource limitation and colimitation.

Our example shows that it can be difficult to access growth rate limitation in laboratory experiments. This is likely due to the difficulty of reducing background contamination as well as biological adaptation to low resource concentrations. In our case, expression of the ammonium transporter AmtB likely helped *E. coli* to maintain high growth rates across a wide range of ammonium concentrations [45]. In addition to laboratory experiments, it is therefore useful to study nutrient colimitation in nature, where growth rate limitation might naturally occur and the full environmental context can be considered. So far it has been challenging to assess the results of environmental datasets because they rely on factorial supplementation experiments (as described in the Introduction; SI Appendix, Fig. S1), and are biased towards certain ecosystems (marine and freshwater) and metabolic types (photosynthetic organisms) [4, 19, 57]. Thus, it is not yet possible to statistically test the hypothesis that resource colimitation is common across organisms, environmental contexts, and outside of specific biochemical dependencies.

Another alternative or complement to factorial supplementation experiments is to develop molecular biomarkers of limitation and colimitation to measure in situ resource status. At least for single resource limitation, this is already occurring in marine biogeochemistry and human microbiome contexts [1, 58, 59]. These biomarkers could be validated and calibrated in the laboratory, then quantified in natural samples. The biomarker approach has the benefit of being able to avoid experimental effects in laboratory or in situ perturbations, and to identify how strain and even sub-strain level resource limitations vary in the community [60]. If resource colimitation represents a distinct physiological state for a population or community, it is possible that specific biomarkers for resource colimitation can also be identified. Our quantitative definitions *L*^rate^, *L*^yield^, 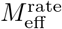, and 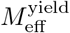 would allow direct, quantitative comparisons between experiments and in situ resource conditions, bridging the gap between the lab and nature and making a truly quantitative biomarker calibration possible.

### Comparing colimitation of growth rate versus growth yield

Our results highlight that there is a fundamental difference between growth rate limitation and biomass yield limitation. Previous work has also referred to these as “kinetic” and “stoichiometric” limitations [3]. Specifically, it is possible for two different resources or sets of resources to be limiting for rate versus yield, and for resources to become growth rate or yield limiting at different concentrations (with growth rate limitation usually occurring at lower resource concentrations; Fig. 3B). In either case, it is not necessarily possible to infer the limiting resource(s) for rate by establishing the limiting resource(s) for yield, and vice versa. While there is a body of literature describing theoretically predicted rate-yield tradeoffs that occur due to fundamental constraints on cellular resource allocation [61], there has been limited empirical evidence in many systems [62, 63]. Distinct resources limiting rate versus yield will not cause these tradeoffs in the first place (since yields evolve neutrally in the absence of spatial structure [64]), but an existing tradeoff could maintain distinct resources limiting rate and yield. The relative importance of limitation for growth rate versus growth yield depends on the context. For microbes, growth rate limitation is likely to be more important to evolution, since growth rate is under direct selection while growth yield may not be (depending on spatial structure and privatization of resources [64]). Yield limitation, on the other hand, is more important in situations where we care about environmental and ecological composition, like elemental cycling in the ocean or total production in biotechnology applications [65–67].

### Growth limitation beyond resources

Resource availability is not the only control on microbial growth, particularly in nature. Limited availability of physical resources such as light and space can limit population growth [68–70]. In the ocean, predators and viral lysis kill a substantial number of cells per day and have quantitative effects on growth rates and standing stocks of microbes [71, 72]. In host-associated microbiomes, such as the human gut, viral lysis also plays a role, as does the host immune system [73, 74]. In all of these contexts, there is also the possibility of inter-microbial warfare such as toxin release, as well as the opposite, promotion of growth through quorum-sensing behaviors due to the presence of other bacteria [75, 76]. There is still much to learn about how these factors interact with resource limitations to influence microbial growth. While the dependence of growth on these other factors is likely more complex than it is on resource availability (e.g., non-monotonic dependence), it is possible to extend the limitation metrics we present here to these cases as well, such that one can consider multiple factors at once (e.g., resource availability and virus concentrations).

### Goals for future work

Resource colimitation of microbial populations and communities may be a common state of nature, instead of a special case. It is a sensible outcome of eco-evolutionary theory and biogeochemistry and has been observed in natural habitats ranging from the surface ocean to the human microbiome. One of the most pressing questions is whether nutrient colimitation is a common or rare state; we propose performing systematic scans of field samples across gradients of multiple resource concentrations to estimate *L* and *M*_eff_, and performing these experiments in a variety of natural systems. We note that this approach is applicable not just to microbes, but across biological systems. The time is also ripe to identify underlying mechanisms of colimitation, which is possible by performing precise physiological comparisons of limitation conditions. For instance, such experiments could address at which level to coarsegrain mechanisms (e.g. whole proteome allocation versus specific biochemical reactions) and identify biomarkers of colimitation that could be leveraged in natural contexts, including potentially in higher trophic level organisms. At the same time, it will also be important to identify how these molecular mechanisms within individual cells/populations are layered to determine colimitation in microbial communities. Lastly, we suggest consideration of colimitation in experimental evolution, ecosystem and biogeochemical models in order to understand the extent to which this phenomenon impacts broader outcomes and may therefore be important for the prediction, management, and use of microbial growth.

## MATERIALS AND METHODS

### Numerical methods

We performed all numerical calculations in Python version 3.10.9, using tools from NumPy [77] version 1.24.1 and SciPy [78] version 1.10.0. We prepared all figures using Matplotlib [79] version 3.9.1. Code for reproducing all analyses and figures is available at: https://github.com/proteoceanLab/Heldetal2024_Nutrient_Colimitation_Theory.

### Measurement of growth rates from luminescence growth curves

For luminescence growth curves we used the *E. coli* strain K-12 MG1655 pCS*λ* [80], which contains a plasmid under kanamycin selection with a luciferase protein expressed under the constitutive lambda CS promoter, and a modified M9 minimal medium base without nitrogen: 12.8 g/L Na_2_HPO_4_ heptahydrate (Sigma Aldrich BioXtra grade lot SLBV4967, CAS S9390-500G), 3 g/L KH_2_PO_4_ monobasic (Sigma Aldrich ACS grade lot SZBF350AV CAS P0062-500G), 0.5 g/L NaCl (Sigma Aldrich BioXtra grade lot SYBG1530V CAS S7653-250G)), 2 mM MgSO_4_ hexahydrate (Sigma Aldrich ACS grade Lot BCBT5460), and 50 µg/mL kanamycin (Sigma Aldrich CAS 7056051-9 Lot 0000130025). We selected three colonies of the strain from an agar plate to serve as biological replicates. We grew separate overnight cultures of each replicate in the base medium along with 10 mM D(+)glucose (Sigma Aldrich BioXtra grade Lot SLBW5196 CAS G7528-1Kg) and 20 mM ammonium chloride (Sigma Aldrich ACS Reagent Lot STBH3180 CAS 31107-500G). After 16 hours of overnight growth, we collected the cells by centrifugation and washed each replicate culture three times in the base medium without glucose and ammonium, then adjusted the OD at 600 nm of each culture to 0.01 by diluting into the base medium. We prepared gradients of glucose (0–2 mM) and ammonium chloride (0–1 mM) in the base medium and aliquoted them into three white-walled, flat-bottom 96-well plates (Corning). We then diluted each replicate culture at a 1:100 ratio into the wells of its corresponding plate, resulting in a final OD of 10^*−*4^ in 200 µL at time zero for the growth curve experiments. We grew the cells at 37 °C for 22 hours in three separate Biotek Synergy H1 plate readers (Agilent Technologies, Inc.), with measurements of luminescence every 10 minutes (orbital shaking of 15 sec prior to each measurement, luminescence integration time of 3 sec). We analyzed the growth curves by 1) performing a background correction of the data using media blanks with no cells added, 2) selecting a time interval of approximate exponential growth, 3) performing linear regression of the log-luminescence in that interval, and 4) quality control based on *R*^2^ for the linear regression and manual inspection of the fit (Dataset S1; SI Appendix, Figs. S6– S8).

### Measurement of growth yields from optical density growth curves

For OD growth curves we used the *E. coli* strain K-12 MG1655 (also containing a chromosomal GFP not used in this experiment) and a modified M9 minimal medium base without nitrogen: 6 g/L Na_2_HPO_4_ (VWR Life Science, 0404, CAS: 7558-79-4), 3 g/L KH_2_PO_4_ (VWR Chemicals, BDH9268, CAS: 777877-0), 0.5 g/L NaCl (VWR Chemicals, BDH9286, CAS: 7647-14-5), 1 mM MgSO_4_ (J.T. Baker, 2506-01, CAS: 7487-88-9), and 0.1 mM CaCl_2_ (VWR Life Science, 1B1110, CAS: 10043-52-4). We selected three colonies of the strain from an agar plate to serve as biological replicates. We grew separate overnight cultures of each replicate in the base medium along with 40 mM glucose (VWR Chemicals, BDH9230, CAS: 50-99-7) and 40 mM ammonium chloride (VWR Life Science, 0621, CAS: 12125-02-9). After 16 hours of overnight growth, we collected the cells by centrifugation and washed each replicate culture five times in the base medium without glucose and ammonium, then adjusted the OD at 600 nm of each culture to 0.02 by diluting into the base medium. We prepared gradients of glucose (0–40 mM) and ammonium chloride (0–40 mM) in the base medium and aliquoted them into three transparent, flat-bottom 96well plates (Corning). We then diluted each replicate culture at a 1:2 ratio into the wells of its corresponding plate, resulting in a final OD of 0.01 in 200 µL at time zero for the growth curve experiments. We grew the cells at 37 °C under shaking conditions (500 RPM) for 24 hours in the Biotek LogPhase 600 plate reader (Agilent Technologies, Inc.), with measurements of OD at 600 nm every 10 minutes. We analyzed the growth curves by 1) performing a background correction of the data using media blanks with no cells added, 2) selecting a time interval of approximate stationary phase, 3) taking an average of the OD in that interval, and 4) quality control based on the coefficient of OD variation during the stationary interval and manual inspection of the fit (Dataset S2; SI Appendix, Figs. S9 and S10).

### Analysis of resource scan data

We fit growth rate and growth yield scans over resource concentrations to models (SI Appendix, Table I and section S3) using `scipy.optimize.curve_fit` for least-squares minimization. We supply initial guesses for each parameter: for the trait value at saturation, we guess the trait value measured at the maximum concentrations of both resources; for the affinities (growth rates) and stoichiometries (growth yields), we guess that maximum trait value divided by half the measured concentration range. We guess exponents of 1 for the Hill and generalized-mean models and zero minimum resource concentrations for models with those parameters. We bound all fitted parameters to be non-negative. We calculated the Akaike information criterion (corrected for small sample size) for each model fit using

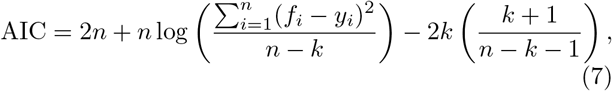

where *n* is the number of data points, *y*_*i*_ is the *i*th measured data point, *f*_*i*_ is the fitted value for that data point, and *k* is the number of parameters in the model [81]. We calculate the relative Akaike weight for a model *α* as

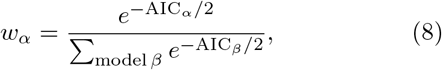

where the sum is over all fitted models. To generate the bootstrapped data sets, we randomly sample one of the three replicate measurements at each concentration condition and subject 100 data sets generated in this way to the same analysis.

### Simulated data sets

We simulate resource scan data sets by taking a model with parameters fit to the experimental data (either growth rate or growth yield) and generating trait values predicted by the model at the same resource concentrations used in the experiments. We add Gaussian noise to each measurement with mean zero and standard deviation that is a linear function of the trait value at that point; this linear function is obtained by performing a linear regression of the standard deviation of growth rates or yields across replicates to the mean growth rate or yield across replicates (SI Appendix, Figs. S15 and S25).

### Virtual supplementation experiments

Let *{R*_glu,*i*_*}* and *{R*_amm,*j*_*}* be the sets of glucose and ammonium concentrations in our resource scan data. We calculate limitation coefficients for virtual supplementation experiments by iterating over each pair of background concentrations *R*_glu,*i*_, *R*_amm,*j*_ and each supplemented concentration *R*_glu,*k*_ *> R*_glu,*i*_ and *R*_amm,*𝓁*_ *> R*_amm,*j*_. For each combination we calculate limitation coefficients as finite differences (Eq. 1):

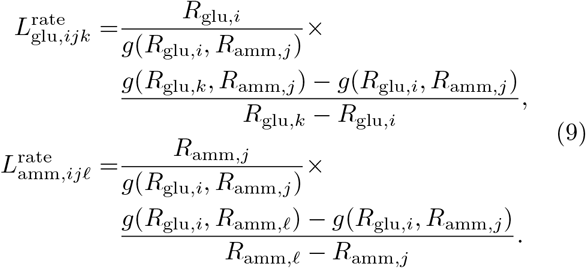

We perform an analogous analysis for the growth yield data.

### Collection and analysis of environmental data

We collected 20 studies in which the parameters of the Monod growth model (*g*_max_ and *K*) for two resoures were measured for the same species/strain. We extracted this data from a broader literature review of Monod parameters that we conducted earlier [38]. Whenever possible, we estimated background environmental nutrient concentrations (i.e., the concentrations the organism is expected to experience in the environment from which it was isolated) from data provided in the original manuscript. When this was not provided, we estimated a range of environmental concentrations from the expected habitat of the organism. For instance, for *E. coli* sp. K12, we selected representative environmental concentrations of resources in the human gut (higher concentrations) and in soils (lower concentrations). For marine diatoms, we selected representative environmental concentrations of high nutrient regions (coastal upwelling zones) and low nutrient regions (surface open ocean). In some cases, the expected resource concentration range is large and spans at least an order of magnitude, in which case we took the midpoint of that range for calculations. Using the measured Monod parameters and estimated environmental concentrations, we calculated limitation coefficients using four common models (SI Appendix, Table I and section S3).

## Supporting information

SI Appendix, including SI Text, Table, and Figures

## ACKNOWLEDGMENTS

We thank Daniel Angst for help running automated growth curve measurements and Justus Fink for valuable feedback on this paper. NAH was supported by the Principles of Microbial Ecosystems collaboration of the Simons Foundation (grant ID 542379 to Martin Ackermann) and an ETH Zurich Career Seed Grant (SEED-26 21-2 to NAH). MM was supported by an Ambizione grant from the Swiss National Science Foundation (PZ00P3 180147).

## Notes

https://github.com/proteoceanLab/Heldetal2024_Nutrient_Colimitation_Theory

### Competing Interest Statement

The authors have declared no competing interest.

### Summary of Updates

1) New statistical analysis of resource concentration scan data; 2) New experimental data for growth rate and growth yield over glucose and ammonium concentrations; 3) Integration of growth rate and growth yield in the results

