## Supplementary material for "Nutrient colimitation is a quantitative, dynamic property of microbial populations": SI Appendix, including SI Text, Table, and Figures

(Dated: August 2, 2024)

### CONTENTS

S1. Mathematical properties of limitation coefficients and number of limiting factors 3
A. Quantitative interpretation of the limitation coefficient 3
B. Range of limitation coefficient values 3
C. Geometric interpretation of $M_{\text{eff}}$ 3
D. Colimitation among a subset of limiting factors 3

S2. Proof of normalization of limitation coefficients 4

S3. Phenomenological models of growth rate and growth yield colimitation 5
A. Liebig models 5
1. Liebig Blackman 5
2. Liebig Monod 6
3. Liebig Hill 7
4. Liebig Bertalanffy 7
B. Multiplicative models 7
1. Multiplicative Blackman 7
2. Multiplicative Monod 7
3. Multiplicative Hill 8
4. Multiplicative Bertalanffy 8
C. Poisson arrival time (PAT)/synthesizing-unit model 8
D. Additive model 9
E. Mankad-Bungay model 10
F. Generalized-sum model 10
G. Mean Monod model of substitutable resources 11
H. Saito substitutable model 12
I. Chemically-dependent resource model 12

S4. Models of population dynamics 12
A. Batch dynamics 12
B. Chemostat dynamics 13

S5. Mechanistic models of growth yield colimitation 14
A. Growth stops only when at least one resource reaches zero concentration 16
B. Growth stops at some nonzero combinations of resource concentrations 16
C. Resource depletion dynamics and biomass yield with resource consumption for maintenance 17
1. Case of one resource 17
2. Two resources 19
D. Resource depletion dynamics and biomass yield with dynamic proteome allocation 19

S6. Effect of unmeasured resources on limitation and colimitation estimates 20

References 20

S7. SI Figures 22

### S1. MATHEMATICAL PROPERTIES OF LIMITATION COEFFICIENTS AND NUMBER OF LIMITING FACTORS

#### A. Quantitative interpretation of the limitation coefficient

Here we develop some quantitative intuition for the growth rate limitation coefficient $L^{\mathrm{rate}}$ (Eq. 1); these points apply similarly to the growth yield limitation coefficient $L^{\mathrm{yield}}$. Consider a small change $\Delta r$ in a limiting process rate $r$ (e.g., a resource uptake rate proportional to its external concentration as in Eq. 4). The growth rate changes according to

$$\begin{aligned} g(r+\Delta r) &\approx g(r) + \Delta r \frac{\partial g}{\partial r} \\ &= g(r)\left(1 + \frac{\Delta r}{r}\frac{r}{g(r)}\frac{\partial g}{\partial r}\right) \\ &= g(r)\left(1 + \frac{\Delta r}{r} L^{\mathrm{rate}}\right). \end{aligned} \tag{S1}$$

Hence we can think of $L^{\mathrm{rate}}$ as the factor by which a relative change in the rate $r$ changes the relative magnitude of growth rate. For example, if $L^{\mathrm{rate}} = 1$, then any relative change in $r$ entails the same relative change in $g$: a 1% change in $r$ means a 1% change in $g$. If $L^{\mathrm{rate}} = 0.5$, then any relative change in $r$ entails only half that relative change in $g$: a 1% change in $r$ means a 0.5% change in $g$. Equivalently, we can think of $L^{\mathrm{rate}}$ as being the exponent of the power-law scaling of $g$ with $r$ around a particular value of $r$:

$$g(r) \sim r^{L^{\mathrm{rate}}}. \tag{S2}$$

#### B. Range of limitation coefficient values

What is the range of values for $L^{\mathrm{rate}}$? In general, we expect that $L^{\mathrm{rate}}$ is zero at high values of $r$, when the process is saturating and no longer limiting. At low values of $r$, $L^{\mathrm{rate}}$ will often equal one, meaning that $g$ is proportional to $r$. This range holds for most common models of growth rate dependence on resource concentrations (SI Appendix, section S3), such as the Blackman (Eq. 5) and Monod (Eq. 6) models. However, negative values are possible if growth rate decreases with increasing $r$, as would occur if the process in question is uptake of an antibiotic; uptake of some resources may also be toxic at extremely high concentrations. Values greater than one are possible if the growth rate scales super-linearly with $r$, as occurs for the Hill model of growth rate [1] (also known as the Moser model [2] or Holling Type III model [3]). This may occur if the process in question involves cooperativity or sensing of a threshold before activation.

#### C. Geometric interpretation of $M_{\mathrm{eff}}$

Under a model where all limitation coefficient range between zero and one, the limitation coefficients form an $(M-1)$-simplex (for $M$ total limiting factors), since they are normalized to sum to one. (Despite the normalization condition, they will not form a simplex for some scenarios where limitation coefficients can be negative or greater than one, as occur in the Hill model or in the presence of antibiotics.) In this case, we can think of $M_{\mathrm{eff}}$ as a way of measuring the distance to the center of the simplex, which has maximum colimitation (all limitation coefficients are equal at $1/M$). Geometrically, $M_{\mathrm{eff}}$ is equivalent to taking whichever limitation coefficient is maximum (i.e., the vertex closest to the state) and measuring the distance along that axis ($\max_i L_i$). Then $M_{\mathrm{eff}}$ is the reciprocal of that, so that higher values mean smaller distance to the simplex center (greater colimitation).

#### D. Colimitation among a subset of limiting factors

Note that while the definition of the effective number of limiting factors $M_{\mathrm{eff}}$ in the main text (Eq. 3) is over all measured factors (e.g., the maximum in the denominator is over all limitation coefficients), sometimes we want to consider the effective number of limiting factors among just a subset of factors. We can do this by renormalizing each limitation coefficient by the sum of limitation coefficients in that subset. That is, among a subset of factors $\mathcal{A}$ we define the relative limitation coefficients

$$L_{i,\mathcal{A}} = \frac{L_i}{\sum_{j\in\mathcal{A}} L_j} \tag{S3}$$

and calculate the effective number of limiting factors within just that subset:

$$\begin{aligned} M_{\mathrm{eff},\mathcal{A}} &= \frac{1}{\max_{i\in\mathcal{A}}\left(L_{i,\mathcal{A}}\right)} \\ &= \frac{1}{\max_{i\in\mathcal{A}}\left(\frac{L_i}{\sum_{j\in\mathcal{A}} L_j}\right)} \\ &= \frac{\sum_{i\in\mathcal{A}} L_i}{\max_{i\in\mathcal{A}} L_i}. \end{aligned} \tag{S4}$$

For example, in our experiments with varying glucose and ammonium, besides calculating the total amount of colimitation among glucose, ammonium, and implicit factors, we may wish to calculate colimitation between just glucose and ammonium. The quantitative range of the number of limiting factors for a subset is the same as it is for the total; it ranges from a minimum of 1 (if only one of the factors is limiting at all) to $|\mathcal{A}|$ (the total number of factors in the subset under consideration, if all are equally limiting).

### S2. PROOF OF NORMALIZATION OF LIMITATION COEFFICIENTS

The total growth rate of biomass $g$ is a function of many individual processes with rates $r_i$:

$$g = g(r_1, r_2, \ldots, r_M). \tag{S5}$$

Since both $g$ and the individual rates $r_i$ must have units of per time, then $g$ must depend on the rates in the following functional form

$$g(r_1, r_2, \ldots, r_M) = r_1 f_1\left(\frac{r_2}{r_1}, \frac{r_3}{r_1}, \ldots, \frac{r_M}{r_1}\right). \tag{S6}$$

We can argue this by dimensional analysis: we must be able to write $g$ as the product of the one the rates (to give $g$ the same units) and a dimensionless function $f$, which being dimensionless must depend only on $M-1$ dimensionless ratios of the rates. For example, the Monod model can be written

$$\begin{aligned} g(r_1, r_2) &= \frac{r_1 r_2}{r_1 + r_2} \\ &= r_1 \frac{1}{r_1/r_2 + 1}, \end{aligned} \tag{S7}$$

where $f(r_2/r_1) = 1/(r_1/r_2 + 1)$. Note that there need not be anything special about $r_1$ — we can write $g$ in that form for any rate $r_i$ for the same dimension argument, although the form of the function $f$ will differ (hence writing it with subscript 1).

We can now calculate the sum of limitation coefficients. First, for $i \neq 1$,

$$\begin{aligned} L_i &= \frac{r_i}{g}\frac{\partial g}{\partial r_i} \\ &= \frac{r_i}{r_1}\frac{f^{(i-1)}\left(\frac{r_2}{r_1}, \frac{r_3}{r_1}, \ldots, \frac{r_M}{r_1}\right)}{f\left(\frac{r_2}{r_1}, \frac{r_3}{r_1}, \ldots, \frac{r_M}{r_1}\right)}. \end{aligned} \tag{S8}$$

And for $L_1$:

$$
\begin{aligned}
L_1 &= \frac{r_1}{g}\frac{\partial g}{\partial r_1} \\
&= \frac{r_1}{g}\left[ f\left(\frac{r_2}{r_1}, \frac{r_3}{r_1}, \ldots, \frac{r_M}{r_1}\right) - \sum_{i=2}^{M} \frac{r_i}{r_1} f^{(i-1)}\left(\frac{r_2}{r_1}, \frac{r_3}{r_1}, \ldots, \frac{r_M}{r_1}\right)\right] \\
&= 1 - \sum_{i=2}^{M} \frac{r_i}{r_1} \frac{f^{(i-1)}\left(\frac{r_2}{r_1}, \frac{r_3}{r_1}, \ldots, \frac{r_M}{r_1}\right)}{f\left(\frac{r_2}{r_1}, \frac{r_3}{r_1}, \ldots, \frac{r_M}{r_1}\right)}.
\end{aligned}
\tag{S9}
$$

We thus see that $L_1 = 1 - \sum_{i=2}^{M} L_i$, and hence $\sum_{i=1}^{M} L_i = 1$.

### S3. PHENOMENOLOGICAL MODELS OF GROWTH RATE AND GROWTH YIELD COLIMITATION

Many different mathematical models of how growth rate depends on resource concentrations have been studied in the literature. Here we summarize a variety of the most common ones, discussing their mechanistic interpretations and limitation properties. This section discusses all models in terms of growth rate $g$, but for fitting the data we apply identical versions of these models to the growth yield as well. Table I lists all models for a generic growth trait $z$ (which could be growth rate $g$ or growth yield $N$) in the special case of two explicit resources, which we use for fitting our scans of *E. coli* growth across glucose and ammonium concentrations (Figs. 2, S13, S20, and S23).

#### A. Liebig models

This is arguably the most widely-used model of multiple-resource dependence [4–8]. It is inspired by the Law of the Minimum attributed to Justus von Liebig [9]: that only a single resource determines growth rate under any given set of conditions. Mathematically, this is expressed as a minimum over a set of functions for each resource. However, there are different types of Liebig models depending on those underlying functions.

##### 1. *Liebig Blackman*

The Blackman model for growth rate (Eq. 5) is a minimum function of the rate for one resource and the rate of all implicit processes [10], so the Liebig Blackman model is a minimum of all resources as well as implicit factors:

$$
\begin{aligned}
g(\mathbf{R}) &= \min_{\text{resource}\, i}\left[\min\left(a_i R_i, g_{\max}\right)\right] \\
&= g_{\max} \min\left(1, \frac{a_1 R_1}{g_{\max}}, \frac{a_2 R_2}{g_{\max}}, \ldots, \frac{a_M R_M}{g_{\max}}\right).
\end{aligned}
\tag{S10}
$$

The limitation coefficients are all zero or one:

$$
\begin{aligned}
L_i^{\text{rate}}(\mathbf{R}) &= \frac{R_i}{g}\frac{\partial g}{\partial R_i} \\
&= \begin{cases} 1 \ \text{ if } \dfrac{a_i R_i}{z_{\max}} = \min\left(1, \dfrac{a_1 R_1}{g_{\max}}, \dfrac{a_2 R_2}{g_{\max}}, \ldots, \dfrac{a_M R_M}{g_{\max}}\right), \\ \\ 0 \ \text{ otherwise.} \end{cases}
\end{aligned}
\tag{S11}
$$

Thus the number of limiting resources $M_{\text{eff}}^{\text{rate}} = 1$ always.

The Liebig Blackman model is arguably the most basic model for yield dependence on resource concentrations, as it holds if we assume fixed stoichiometry for all resources and that growth stops only once at least one resource is completely exhausted.

| **Model name** | **Parameters** | **Formula for trait $z$ (growth rate or growth yield)** | **Colimitation between resources?** | **Colimitation with implicit factors?** |
|---|---|---|---|---|
| Liebig Blackman | $z_{\max}$,$a_1$,$a_2$ | $z_{\max} \min\left(\frac{a_1R_2}{z_{\max}}, \frac{a_2R_2}{z_{\max}}, 1\right)$ | No | No |
| Liebig Monod | $z_{\max}$,$a_1$,$a_2$ | $z_{\max} \min\left(\frac{a_1R_1}{a_1R_1 + z_{\max}}, \frac{a_2R_2}{a_2R_2 + z_{\max}}\right)$ | No | Yes |
| Liebig Hill | $z_{\max}$,$a_2$,$a_2$, $n_1$,$n_2$ | $z_{\max} \min\left(\frac{(a_1R_1)^{n_1}}{(a_1R_1)^{n_1} + z_{\max}^{n_1}}, \frac{(a_2R_2)^{n_2}}{(a_2R_2)^{n_2} + z_{\max}^{n_2}}\right)$ | No | Yes |
| Liebig Bertalanffy | $z_{\max}$,$a_1$,$a_2$ | $z_{\max} \min\left(1 - 2^{-a_1R_1/z_{\max}}, 1 - 2^{-a_2R_2/z_{\max}}\right)$ | No | Yes |
| Multiplicative Blackman | $z_{\max}$,$a_1$,$a_2$ | $z_{\max} \min\left(\frac{a_1R_2}{z_{\max}}, 1\right) \cdot \min\left(\frac{a_2R_2}{z_{\max}}, 1\right)$ | Yes | No |
| Multiplicative Monod | $z_{\max}$,$a_1$,$a_2$ | $z_{\max}\left(\frac{a_1R_1}{a_1R_1 + z_{\max}}\right)\left(\frac{a_2R_2}{a_2R_2 + z_{\max}}\right)$ | Yes | Yes |
| Multiplicative Hill | $z_{\max}$,$a_2$,$a_2$, $n_1$,$n_2$ | $z_{\max}\left(\frac{(a_1R_1)^{n_1}}{(a_1R_1)^{n_1} + z_{\max}^{n_1}}\right)\left(\frac{(a_2R_2)^{n_2}}{(a_2R_2)^{n_2} + z_{\max}^{n_2}}\right)$ | Yes | Yes |
| Multiplicative Bertalanffy | $z_{\max}$,$a_1$,$a_2$ | $z_{\max}\left(1 - 2^{-a_1R_1/z_{\max}}\right)\left(1 - 2^{-a_2R_2/z_{\max}}\right)$ | Yes | Yes |
| Poisson arrival time/ synthesizing unit | $z_{\max}$,$a_1$,$a_2$ | $z_{\max}\frac{a_1R_1a_2R_2(a_1R_1 + a_2R_2)}{a_1R_1a_2R_2(a_1R_1 + a_2R_2) + z_{\max}[(a_1R_1)^2 + a_1R_1a_2R_2 + (a_2R_2)^2]}$ | Yes | Yes |
| Additive | $z_{\max}$,$a_1$,$a_2$ | $\frac{1}{(a_1R_1)^{-1} + (a_2R_2)^{-1} + z_{\max}^{-1}}$ | Yes | Yes |
| Mankad-Bungay | $z_{\max}$,$a_1$,$a_2$ | $z_{\max}\left(\frac{a_1R_1a_2R_2}{a_1R_1 + a_2R_2}\right)\left(\frac{1}{a_1R_1 + z_{\max}} + \frac{1}{a_2R_2 + z_{\max}}\right)$ | Yes | Yes |
| Generalized mean | $z_{\max}$,$a_1$,$a_2$, $q$ | $\frac{1}{\left((a_1R_1)^{-q} + (a_2R_2)^{-q} + z_{\max}^{-q}\right)^{1/q}}$ | Yes | Yes |
| Saito substitutable | $z_{\max}$,$a_1$,$a_2$ | $g_{\max}\frac{a_1R_1 + a_2R_2}{a_1R_1 + a_2R_2 + z_{\max}}$ | Yes | Yes |
| Mean Monod | $z_{\max}$,$a_1$,$a_2$ | $g_{\max}\left(\frac{1}{2}\frac{a_1R_1}{a_1R_1 + z_{\max}} + \frac{1}{2}\frac{a_2R_2}{a_2R_2 + z_{\max}}\right)$ | Yes | Yes |
| Chemically-dependent | $z_{\max}$,$a_1$,$a_2$ | $z_{\max}\frac{a_1R_1a_2R_2}{a_1R_1a_2R_2 + z_{\max}(a_2R_2 + z_{\max})}$ | Yes | Yes |

TABLE I. Summary of all trait models for fitting resource scans of growth rate and growth yield. Each model has a corresponding "Rmin" version in which the resource concentrations $R_1$ and $R_2$ are shifted by additional free parameters $R_{1,\min}$ and $R_{2,\min}$. See Figs. S13 and S23 for fits of all these models to the growth rate and growth yield data.

#### *2. Liebig Monod*

This is the most common form of Liebig model, in which the minimum is over individual Monod models for each resource (Fig. S2, first row and first column):

$$g(\mathbf{R}) = g_{\max} \min_{\text{resource } i}\left(\frac{a_iR_i}{a_iR_i + g_{\max}}\right). \tag{S12}$$

The limitation coefficients in the Liebig model are (Fig. S2, second row and first column)

$$L_i^{\text{rate}}(\mathbf{R}) = \frac{R_i}{g}\frac{\partial g}{\partial R_i} = \begin{cases} \dfrac{g_{\max}}{a_i R_i + g_{\max}} & \text{if } i = \underset{\text{resource } j}{\arg\min}\left(\dfrac{a_j R_j}{a_j R_j + g_{\max}}\right), \\ 0 & \text{otherwise.} \end{cases} \tag{S13}$$

#### *3. Liebig Hill*

The Liebig Hill model uses Hill functions for each resource:

$$g(\mathbf{R}) = g_{\max} \min_{\text{resource } i}\left(\frac{(a_i R_i)^{n_i}}{(a_i R_i)^{n_i} + g_{\max}^{n_i}}\right), \tag{S14}$$

where $n_i$ is a cooperativity coefficient for each resource. When $n_i = 1$, the dependence is equivalent to Monod, but $n_i > 1$ creates cooperativity such that growth rate increases super-linearly with resource concentration at low concentrations. In particularly, this allows for switch-like dependence such that growth rate depends only weakly on a resource when it is low, but then rapidly jumps to a maximum concentration above a threshold concentration of that resource [1]. The super-linear dependence in the Hill model entails limitation coefficients greater than one for some resources (SI Appendix, section S1).

#### *4. Liebig Bertalanffy*

This model uses Bertalanffy functions [11] for each resource, which saturates more rapidly at high resource concentrations (exponentially) compared to the Monod model (hyperbolically):

$$g(\mathbf{R}) = g_{\max} \min_{\text{resource } i}\left(1 - 2^{-a_i R_i / g_{\max}}\right). \tag{S15}$$

### B. Multiplicative models

Multiplicative models are also a common form of growth rate dependence on multiple resources [5, 6, 11]. These are based on the law of mass action, in which the reaction rate is a product of the concentrations of all substrates. The multiplicative form means that the limitation coefficient for each resource is independent of all other resource concentrations.

#### *1. Multiplicative Blackman*

This model is analogous to the Liebig Blackman model, but multiplying individual Blackman models for each resource:

$$g(\mathbf{R}) = g_{\max} \prod_{\text{resource } i} \min\left(\frac{a_i R_i}{g_{\max}}, 1\right). \tag{S16}$$

Note that parameterize this slightly differently (rescaling $a_i R_i$ by $g_{\max}$) so that the product across resources is dimensionless and can thus be rescaled by $g_{\max}$ overall to get the total rate.

#### *2. Multiplicative Monod*

This model is a product of Monod models for each resource [5, 11] (Fig. S2, first row and fourth column):

$$g(\mathbf{R}) = g_{\max} \prod_{\text{resource } i} \frac{a_i R_i}{a_i R_i + g_{\max}}. \tag{S17}$$

Since the growth rate exactly factorizes into separate contributions from each resource, the limitation coefficients for the multiple-resource case are the same as for the single-resource Monod model (Fig. S2, second row and fourth column):

$$\begin{aligned} L_i^{\text{rate}}(\mathbf{R}) &= \frac{R_i}{g}\frac{\partial g}{\partial R_i} \\ &= \frac{g_{\text{max}}}{a_i R_i + g_{\text{max}}}. \end{aligned} \tag{S18}$$

#### 3. *Multiplicative Hill*

This model is a product of Hill models for each resource [1]:

$$\text{Multiplicative Hill model [1]: } g(\mathbf{R}) = g_{\text{max}} \prod_{\text{resource } i} \frac{R_i^{n_i}}{R_i^{n_i} + K_i^{n_i}}, \tag{S19}$$

Like the Liebig Hill model, it allows for limitation coefficients greater than one in some regimes. In particular, it is possible that multiple resources could have limitation coefficients greater than one simultaneously, which means that the implicit limiting factors must actually have negative limitation coefficient (because the sum of limitation coefficients remains normalized to one). Counterintuitively, this indicates that increasing the rate of the implicit processes actually slows down growth.

#### 4. *Multiplicative Bertalanffy*

This model is a product of Bertalanffy functions for each resource [11]:

$$g(\mathbf{R}) = g_{\text{max}} \prod_{\text{resource } i} \left(1 + 2^{-a_i R_i / g_{\text{max}}}\right). \tag{S20}$$

### C. Poisson arrival time (PAT)/synthesizing-unit model

The PAT model (also often known as the synthesizing-unit model) derives from a semi-mechanistic view of biomass growth [4, 8, 11, 12]. The idea is that the time $t_{\text{biomass}}$ to make a new unit of biomass is the sum of the time $t_{\text{uptake,total}}$ to uptake units of all resources and the time $t_{\text{metabolism}}$ to metabolize those resources into new biomass:

$$t_{\text{biomass}} = t_{\text{uptake,total}} + t_{\text{metabolism}}. \tag{S21}$$

In general, these times for uptake and metabolism are stochastic, described by some probability distributions. We approximate the overall growth rate of biomass as the reciprocal of the mean time:

$$\begin{aligned} g &= \frac{1}{\langle t_{\text{biomass}} \rangle} \\ &= \frac{1}{\langle t_{\text{uptake,total}} \rangle + \langle t_{\text{metabolism}} \rangle}. \end{aligned} \tag{S22}$$

We assume the mean time of metabolism $\langle t_{\text{metabolism}} \rangle$ does not depend on the environmental resource concentrations $\mathbf{R}$, since metabolism relies on resources already recruited into cells in sufficient quantities. Without loss of generality, we therefore parameterize the mean time of metabolism as $\langle t_{\text{metabolism}} \rangle = 1/g_{\text{max}}$, since this will be the maximum growth rate if uptake time is zero.

The PAT model assumes that uptake of each resource unit occurs as an independent Poisson process, where the uptake time $t_{\text{uptake},i}$ of the $i$th resource has an exponential probability distribution

$$p_{\text{uptake},i}(t_{\text{uptake},i}) = a_i R_i e^{-a_i R_i t_{\text{uptake},i}}. \tag{S23}$$

We therefore assume the rate of each resource unit's uptake is proportional to the concentration $R_i$ of the resource in the environment, with a constant of proportionality $a_i$. Since uptake of each resource unit occurs independently, the total uptake time for all resource units is the maximum of their individual uptake times:

$$t_{\text{uptake,total}} = \max_{\text{resource } i} t_{\text{uptake},i}. \tag{S24}$$

We want to calculate the average total uptake time so we can calculate the growth rate using Eq. S22. The cumulative probability that the total uptake time $t_{\text{uptake,total}}$ is less than $t$ equals the probability that all individual resource uptake times are less than $t$:

$$\begin{aligned} p\left(t_{\text{uptake,total}} < t\right) &= \prod_{\text{resource } i} p_i(t_{\text{uptake},i} < t) \\ &= \prod_{\text{resource } i} \left(1 - e^{-a_i R_i t}\right), \end{aligned} \tag{S25}$$

where we have used the exponential distribution for the individual resource uptake times (Eq. S23) and the fact these times are independent of each other. The probability distribution of the total uptake time $t_{\text{uptake,total}}$ is the derivative of this cumulative distribution:

$$\begin{aligned} p(t_{\text{uptake,total}}) &= \frac{d}{dt}\, p\left(t_{\text{uptake,total}} < t\right) \\ &= \sum_{\text{resource } i} a_i R_i e^{-a_i R_i t_{\text{uptake,total}}} \prod_{\text{resource } j \neq i} \left(1 - e^{-a_j R_j t_{\text{uptake,total}}}\right). \end{aligned} \tag{S26}$$

We can now calculate the average total uptake time:

$$\begin{aligned} \langle t_{\text{uptake,total}} \rangle &= \int_0^\infty dt_{\text{uptake,total}}\; p(t_{\text{uptake,total}})\; t_{\text{uptake,total}} \\ &= \sum_{\text{resource } i} a_i R_i \int_0^\infty dt_{\text{uptake,total}}\; t_{\text{uptake,total}} e^{-a_i R_i t_{\text{uptake,total}}} \prod_{\text{resource } j \neq i} \left(1 - e^{-a_j R_j t_{\text{uptake,total}}}\right) \\ &= \sum_{\text{resource } i} \frac{1}{a_i R_i} - \sum_{\text{resource pairs } i,j} \frac{1}{a_i R_i + a_j R_j} + \sum_{\text{resource triplets } i,j,k} \frac{1}{a_i R_i + a_j R_j + a_k R_k} - \cdots, \end{aligned} \tag{S27}$$

where the sums are over individual, pair, triplet, etc. combinations of distinct resources (e.g., pairs of resource indices $i$ and $j$ such that $i \neq j$). Note the alternating signs across these sums.

Thus the overall growth rate from Eq. S22 for the PAT model is therefore (Fig. S2, first row and second column)

$$g(\mathbf{R}) = \frac{g_{\max}}{1 + g_{\max} \langle t_{\text{uptake,total}} \rangle}, \tag{S28}$$

where we calculate $g_{\max}\langle t_{\text{uptake,total}}\rangle$ using Eq. S27, which contains the dependence on the resource concentrations $\mathbf{R}$.

Using Eqs. S27 and S28, the limitation coefficients in the PAT model are (Fig. S2, second row and second column)

$$\begin{aligned} L_i^{\text{rate}}(\mathbf{R}) =& \frac{R_i}{g} \frac{\partial g}{\partial R_i} \\ =& -\frac{g}{g_{\max}} R_i \frac{\partial}{\partial R_i} \left(g_{\max} \langle t_{\text{uptake,total}} \rangle\right) \\ =& g a_i R_i \left( \frac{1}{(a_i R_i)^2} - \sum_{\text{resource } j \neq i} \frac{1}{(a_i R_i + a_j R_j)^2} + \sum_{\text{resources } j \neq i, k \neq i, k \neq j} \frac{1}{(a_i R_i + a_j R_j + a_k R_k)^2} - \cdots \right). \end{aligned} \tag{S29}$$

### D. Additive model

The motivation for this model is similar to that of the PAT model, in terms of decomposing biomass growth into uptake of individual units of resources followed by metabolism. However, instead of assuming uptake processes of all resource units occur in parallel, such that the total uptake time is the maximum of these individual uptake times (Eq. S24), we assume that the uptake processes occur sequentially, such that the individual uptake times add up to the total uptake time [4, 11]:

$$t_{\text{uptake,total}} = \sum_{\text{resource } i} t_{\text{uptake},i}. \tag{S30}$$

We otherwise make the same assumptions, i.e., that uptake of each resource unit is a Poisson process with rate $a_i R_i$. Therefore the growth rate is (Fig. S2, first row and third column)

$$\begin{aligned} g(\mathbf{R}) &= \frac{1}{\langle t_{\text{metabolism}} \rangle + \langle t_{\text{uptake,total}} \rangle} \\ &= \frac{1}{\langle t_{\text{metabolism}} \rangle + \sum_{\text{resource } i} \langle t_{\text{uptake},i} \rangle} \\ &= \frac{1}{g_{\max}^{-1} + \sum_{\text{resource } i} (a_i R_i)^{-1}} \end{aligned} \tag{S31}$$

The limitation coefficients for the additive model are (Fig. S2, second row and third column)

$$\begin{aligned} L_i^{\text{rate}}(\mathbf{R}) &= \frac{R_i}{g} \frac{\partial g}{\partial R_i} \\ &= \frac{(a_i R_i)^{-1}}{g_{\max}^{-1} + \sum_{\text{resource } j} (a_j R_j)^{-1}}. \end{aligned} \tag{S32}$$

### E. Mankad-Bungay model

The Mankad-Bungay model is a phenomenological model proposed by Mankad and Bungay [13, 14] in the form of a weighted sum of Monod functions for each resource (originally for two resources only, but generalized here to an arbitrary number):

$$\begin{aligned} g(\mathbf{R}) &= g_{\max} \sum_{\text{resource } i} \left( \frac{(a_i R_i)^{-1}}{\sum_j (a_j R_j)^{-1}} \right) \left( \frac{a_i R_i}{a_i R_i + g_{\max}} \right) \\ &= g_{\max} \frac{\sum_{\text{resource } i} (a_i R_i + g_{\max})^{-1}}{\sum_{\text{resource } i} (a_i R_i)^{-1}}. \end{aligned} \tag{S33}$$

Note that the Mankad-Bungay model has non-monotonic dependence on resource concentrations in some regimes, unlike all other models considered here. Specifically, the growth rate decreases as a function of $R_1$ when

$$R_1 > \frac{1}{a_1} \left( a_2 R_2 + \sqrt{2 a_2 R_2 (g_{\max} + a_2 R_2)} \right). \tag{S34}$$

### F. Generalized-sum model

The aforementioned additive model is based on the idea that the total time to uptake all resources is the sum of uptake times for each one. This means the growth rate is the harmonic sum (reciprocal sum of reciprocals) of the rates for each step in the process. A phenomenological way to generalize this is to use a generalized sum, based on the idea of the generalized (or power) mean [15]; this is a family of ways to average over a set of numbers, encompassing several more common means such as the arithmetic, harmonic, and geometric means. The generalized sum model of growth rate is

$$g(\mathbf{R}) = \left( g_{\max}^{-q} + \sum_{\text{resource } i} (a_i R_i)^{-q} \right)^{-1/q}. \tag{S35}$$

This definition has several convenient properties, mostly mediated by the key parameter $q$. First we consider the regime where $q > 0$. Since the growth rate $g(\mathbf{R})$ will then always be zero if any one resource $i$ has zero concentration

$R_i$, this regime describes essential, non-substitutable resources, all of which are required for nonzero growth. It also has linear proportionality to $R_i$ in the limit where $R_i \ll g_{\text{max}}/a_i$ and $R_i \ll a_j R_j/a_i$ for all other resources $j \neq i$.

One of the most important properties of the generalized-sum model is that in the limit of $q \to \infty$, the model recovers Liebig Blackman dependence on resources:

$$\lim_{q\to\infty} g(\mathbf{R}) = \min\left(g_{\text{max}}, a_1 R_1, a_2 R_2, \ldots, a_M R_M\right). \tag{S36}$$

Finite values of $q$ thus represent deviations from this Liebig limit. Another interesting limit is $q = 1$, where the growth rate is equivalent to the additive model:

$$\lim_{q\to 1} g(\mathbf{R}) = \left(g_{\text{max}}^{-1} + \sum_{\text{resource } i} (a_i R_i)^{-1}\right)^{-1}. \tag{S37}$$

Note that with positive values of $q$, the smallest values of $a_i R_i$ always dominate the sum, and the magnitude of $q$ determines by how much — infinite $q$ represents complete dominance of the minimum (Liebig case), while finite $q$ allows for a soft dependence on the other non-minimum resources. We also point out that the growth rate $g(\mathbf{R})$ is a monotonically increasing function of the parameter $q$, so that the Liebig case ($q \to \infty$) always represents the greatest growth rate that can be produced for a given set of resources. Finite values of $q$ represent less efficient growth rate. Altogether this suggests that we can think of the magnitude of the parameter $q$ as measuring the interaction between resources in determining the total growth rate.

The limitation coefficients in the generalized-sum model are

$$\begin{aligned} L_i^{\text{rate}}(\mathbf{R}) &= \frac{R_i}{g}\frac{\partial g}{\partial R_i} \\ &= \frac{(a_i R_i)^{-q}}{g_{\text{max}}^{-q} + \sum_{\text{resource } j} (a_j R_j)^{-q}}. \end{aligned} \tag{S38}$$

Note these limitation coefficients range from 0 (if resource $i$ is unlimited, so $R_i \to \infty$) to 1 (if resource $i$ is the only finite resource). We can use these limitation coefficients to quantify the concentrations at which limitation switches from one resource to another.

#### G. Mean Monod model of substitutable resources

So far we have considered only resources that are essential and non-substitutable, like a carbon source and a nitrogen source. However, some resources are substitutable, such as a two different carbon sources, so it is valuable to consider growth rate limitation on these resources as well. If substitutable resources are consumed sequentially (diauxie), then their growth rate limitation is the same as for single resources, since by definition only a single resource affects growth rate at any instant in time. Therefore the important case to consider here is when the resources are consumed (and therefore affect growth rate) simultaneously.

There are two common models for how growth rate would depend on such resource concentrations. The first is that growth rate is a weighted average of Monod models over all resources (Fig. S2, second column and first row) [16]:

$$g(\mathbf{R}) = g_{\text{max}} \sum_{\text{resource } i} \alpha_i \frac{a_i R_i}{a_i R_i + g_{\text{max}}}, \tag{S39}$$

where the weights $\alpha_i$ are normalized: $\sum_{\text{resource } i} \alpha_i = 1$. In this model, growth rate is zero only if all resource concentrations are zero; compare this to the models of non-substitutable resources, where growth rate is zero if *any* resource concentration is zero. The growth rate saturates at $g_{\text{max}}$ if all resources are unlimited, but note that if one resource is unlimited, the growth rate still responds to changes in concentration of the other resources. In this model, the limitation coefficients are (Fig. S2, second column and second row)

$$\begin{aligned} L_i^{\text{rate}}(\mathbf{R}) &= \frac{R_i}{g}\frac{\partial g}{\partial R_i} \\ &= \frac{\alpha_i a_i R_i g_{\text{max}}/(a_i R_i + g_{\text{max}})^2}{\sum_{\text{resource } j} \alpha_j a_j R_j/(a_j R_j + g_{\text{max}})}. \end{aligned} \tag{S40}$$

#### H. Saito substitutable model

A second model for growth rate dependence on substitutable resources is (Fig. S2, first column and first row) [6]

$$g(\mathbf{R}) = g_{\max} \frac{\sum_{\text{resource } i} a_i R_i}{g_{\max} + \sum_{\text{resource } i} a_i R_i}. \tag{S41}$$

Note that this model differs from Eq. S39 in that if any one resource is unlimited, then the growth rate is constant, regardless of the concentrations of the other resources. The limitation coefficients are (Fig. S2, first column and second row)

$$\begin{aligned} L_i^{\text{rate}}(\mathbf{R}) &= \frac{R_i}{g} \frac{\partial g}{\partial R_i} \\ &= \frac{a_i R_i / g_{\max}}{\left(\sum_{\text{resource } j} a_j R_j\right)\left(g_{\max} + \sum_{\text{resource } j} a_j R_j\right)}. \end{aligned} \tag{S42}$$

#### I. Chemically-dependent resource model

Another possible relationship between non-substitutable resources is where one resource may be chemically-dependent on the other for uptake or usage [6]. This is true especially for pairs of resources involving a metal serving as an enzyme cofactor; for example, the growth rate dependence on bicarbonate in marine diatoms depends on the availability of zinc [6]. In this case, growth rate will depend asymmetrically on the concentrations of both resources, since the dependent resource will have a weaker effect. In the case where resource 1 depends on resource 2, one model for growth rate is this case is (Fig. S2, third column and first row) [6]

$$g(R_1, R_2) = g_{\max} \frac{a_1 R_1 a_2 R_2}{a_1 R_1 a_2 R_2 + g_{\max}(a_2 R_2 + g_{\max})}. \tag{S43}$$

We can interpret this such that the effective half-saturation concentration for resource 1 depends on the concentration of resource 2: at low concentrations of resource 2, the effective half-saturation concentration for resource 1 becomes very large, while at high concentrations of resource 2, the half-saturation concentration for resource 1 decreases down to approximately an intrinsic lower limit of $K_1 = g_{\max}/a_1$.

The limitation coefficient for the dependent resource is (Fig. S2, third column and second row)

$$\begin{aligned} L_1^{\text{rate}}(R_1, R_2) &= \frac{R_1}{g} \frac{\partial g}{\partial R_1} \\ &= \frac{g_{\max}(a_2 R_2 + g_{\max})}{a_1 R_1 a_2 R_2 + g_{\max}(a_2 R_2 + g_{\max})}. \end{aligned} \tag{S44}$$

Again, this is equivalent to the limitation coefficient for resource 1 by itself but with an effective half-saturation concentration that depends on $R_2$. The limitation coefficient for the independent resource is (Fig. S2, third column second row)

$$\begin{aligned} L_2^{\text{rate}}(R_1, R_2) &= \frac{R_2}{g} \frac{\partial g}{\partial R_2} \\ &= \frac{g_{\max}^2}{a_1 R_1 a_2 R_2 + g_{\max}(a_2 R_2 + g_{\max})}. \end{aligned} \tag{S45}$$

### S4. MODELS OF POPULATION DYNAMICS

#### A. Batch dynamics

Batch dynamics entails an initial biomass being given a fixed initial concentration of resources, which it consumes until growth stops. Let $N(t)$ be the concentration of biomass and $\mathbf{R}(t) = (R_1(t), R_2(t), \ldots, R_M(t))$ be the vector of all $M$ resource concentrations in the environment at time $t$, with some fixed initial conditions $N(0)$ and $\mathbf{R}(0)$. Their dynamics over time are determined by

$$\begin{aligned} \frac{d}{dt}N(t) &= g(\mathbf{R}(t))N(t), \\ \frac{d}{dt}R_i(t) &= -\frac{1}{Y_i}\frac{d}{dt}N(t), \end{aligned} \tag{S46}$$

where $g(\mathbf{R})$ is the per-capita biomass growth rate that depends on the environmental resource concentrations and $Y_i$ is the intrinsic yield (amount of new biomass produced per unit resource, with all other resource unlimited) for resource $i$. We solve these equations numerically using `scipy.integrate.solve_ivp` (e.g., Fig. 1C,E).

#### B. Chemostat dynamics

Here we define a model analogous to the batch case (Eq. S46) but for a chemostat:

$$\begin{aligned} \frac{d}{dt}N(t) &= \left(g(\mathbf{R}) - d\right)N(t), \\ \frac{d}{dt}R_i(t) &= -\frac{1}{Y_i}g(\mathbf{R})N(t) + d\left(R_i^{\text{source}} - R_i(t)\right), \end{aligned} \tag{S47}$$

where $R_i^{\text{source}}$ is the concentration of resource $i$ in the source media flowing into the chemostat and $d$ is the dilution factor ($d = \omega/V$, where $\omega$ is volume flowing in and out per unit time and $V$ is the total volume). Here we assume constant yields $Y_i$, but we can generalize our results to the case of variable yields as we do in the batch case (see SI Appendex, section S5). Note that we express the resource consumption by biomass (first term on the right-hand side of $dR_i/dt$) as being proportional to $g(\mathbf{R})N(t)$ rather than $dN/dt$ as in the batch model (Eq. S46). This is because $dN/dt$ includes a death process as well as the growth of new biomass, while we assume resource consumption is only associated with growth of new biomass.

For chemostats we primarily care about their steady states, so let $N^*$ and $\mathbf{R}^*$ be the steady-state concentrations of biomass and resources. To calculate these concentrations, we first set $dN/dt = 0$, which gives us the condition that the growth rate must equal the dilution rate:

$$g(\mathbf{R}^*) = d. \tag{S48}$$

Assuming there are $M$ total resources, this defines an $(M-1)$-dimensional manifold in the space of resource concentrations $\mathbf{R}^*$. Next we must also set $dR_i/dt = 0$, which gives us

$$N^* = (R_i^{\text{source}} - R_i^*)Y_i. \tag{S49}$$

Note this requires that $R_i^{\text{source}} > R_i^*$ for a steady state to exist. Since this must hold for each resource $i$, we must have

$$(R_1^{\text{source}} - R_1^*)Y_1 = (R_2^{\text{source}} - R_2^*)Y_2 = \cdots = (R_M^{\text{source}} - R_M^*)Y_M. \tag{S50}$$

These equations are also apparent from the assumption of constant stoichiometry of resources and biomass: the difference in resource concentrations between the source and the vessel corresponds to what the population is consuming, and that consumed quantity must exactly match the stoichiometry of the biomass ($Y_i/Y_j$) for it to be in steady state. To solve the steady state, we must solve for solutions of Eqs. S48 and S50 ($M$ equations) to get the resource concentrations $\mathbf{R}^*$ ($M$ unknowns), and then plug those into Eq. S49 to get the biomass concentration $N^*$. In Fig. 1D,F we plot the dynamics to steady state for an example chemostat model.

We define growth rate limitation under chemostat conditions as the response of the instantaneous growth rate to a small perturbation in resource concentration at steady state. Note that the response to this perturbation represents a transient departure from steady state itself. First, let us consider the case of a single resource. In this case the steady-state resource concentration is determined by $g(R^*) = d$ (Eq. S48). Assuming a Monod model of growth rate $g(R)$, the resource concentration is (assuming $R^* < R^{\text{source}}$ such that a steady state exists)

$$R^* = K\frac{d}{g_{\max} - d}. \tag{S51}$$

Since the growth rate limitation coefficient for a single resource in the Monod model is $L^{\text{rate}}(R) = K/(R+K)$ (Eq. 1), the limitation coefficient for a steady-state chemostat is

$$\begin{aligned} L^{\text{rate}}(R^*) &= \frac{K}{R^* + K} \\ &= 1 - \frac{d}{g_{\max}}. \end{aligned} \tag{S52}$$

Therefore if the dilution rate is far below the maximum growth rate of the population ($d \ll g_{\max}$), then limitation is very close to its maximum value of $L = 1$. This makes sense as limitation is highest when growth rates are slow, which corresponds to low resources. In contrast, if the dilution rate $d$ is very close to the maximum growth rate $g_{\max}$ (fast growth), then limitation becomes close to its minimum value of $L = 0$, which corresponds to a high steady-state resource concentration $R^*$.

For multiple resources, we repeat this procedure, except we must typically calculate the steady-state concentrations $\mathbf{R}^*$ numerically from the nonlinear Eqs. S48 and S50. We can then plug these concentrations into equations for the limitation coefficients under any model of growth rate (SI Appendix, section S3 and Table I). We show examples of these limitation coefficients over chemostat dynamics in Fig. 1E.

We define yield limitation of a population growing under chemostat conditions as the response of its steady-state biomass to changes in the source concentrations of resources being supplied. Let us first consider the case of a single resource. The biomass yield at steady state is (Eq. S49)

$$\begin{aligned} N^* &= (R^{\text{source}} - R^*)Y \\ &= \left( R^{\text{source}} - K \frac{d}{g_{\max} - d} \right) Y, \end{aligned} \tag{S53}$$

where we have used the solution for $R^*$ in Eq. S51. Therefore the yield limitation coefficient is (Eq. 1)

$$\begin{aligned} L^{\text{yield}} &= \frac{R^{\text{source}}}{N^*} \frac{\partial N^*}{\partial R^{\text{source}}} \\ &= \frac{R^{\text{source}}}{R^{\text{source}} - Kd/(g_{\max} - d)}. \end{aligned} \tag{S54}$$

When $R^{\text{source}}$ is high, then the limitation coefficient takes approximately its minimum value of $L^{\text{yield}} = 1$: biomass yield responds approximately linearly to changes in $R^{\text{source}}$. When $R^{\text{source}}$ is low (i.e., close to its lower bound set by the dilution rate $d$ and intrinsic growth traits $g_{\max}$ and $K$ such that a steady state is feasible), then limitation is much larger than 1. Note that while the biomass is always a linear function of $R^{\text{source}}$ (Eq. S49), there is an apparent super-linearity here due to the shift in $R^{\text{source}}$ by $R^*$. So on a log-log scale, $N^*$ will increase faster than linear with $R^{\text{source}}$ near that minimum value.

We follow a similar procedure for multiple resources but with one complication. Unlike in the single-resource case, the steady-state concentrations $R_i^*$ depend on the source concentrations $R_i^{\text{source}}$ under multiple resources. Mathematically, this is because the steady states $R_i^*$ are defined as solutions of Eqs. S48 and S50, where the latter depends on the source concentrations $R_i^{\text{source}}$. Intuitively, this is because the source concentrations determine where along the manifold $g(\mathbf{R}^*) = d$ the resource depletion trajectory intersects (e.g., Fig. 1F). Therefore the yield limitation coefficients can generally be written only as

$$L_i^{\text{yield}} = \frac{R_i^{\text{source}}}{(R_i^{\text{source}} - R_i^*)} \left( 1 - \frac{\partial R_i^*}{\partial R_i^{\text{source}}} \right). \tag{S55}$$

Therefore one must solve Eqs. S48 and S50 and calculate the derivative of $R_i^*$ with respect to $R_i^{\text{source}}$ (or do so numerically) to determine the limitation coefficient.

### S5. MECHANISTIC MODELS OF GROWTH YIELD COLIMITATION

Here we describe the growth and resource depletion of a batch culture of microbes consuming two essential, non-substitutable resources in the environment; the model and its properties are straightforward to generalize to an arbitrary number of resources. Let $N(t)$ be the concentration of biomass at time $t$ and $R_i(t)$ be the environmental concentration of resource $i$ at time $t$, with initial conditions $N(0)$ and $R_i(0)$. The dynamics of these concentrations are determined by (Fig. 1C):

$$
\begin{aligned}
\frac{d}{dt}N(t) &= g(R_1(t), R_2(t))N(t), \\
\frac{d}{dt}R_1(t) &= -\frac{1}{Y_1(R_1(t), R_2(t))}\frac{d}{dt}N(t), \\
\frac{d}{dt}R_2(t) &= -\frac{1}{Y_2(R_1(t), R_2(t))}\frac{d}{dt}N(t),
\end{aligned}
\tag{S56}
$$

where $g$ is the per-capita growth rate (assumed to depend only on the instantaneous environmental resource concentrations through a model as in SI Appendix, section S3 and Table I) and $Y_i(R_1, R_2)$ is the biomass yield for resource $i$ (amount of new biomass produced per unit of resource $i$). Note that here we allow for the possibility that the yields can change depending on the environmental resource concentrations. Equation S56 assumes that resources are consumed only by growth of new biomass (hence the proportionality of $dR_i/dt$ with $dN/dt$), but in a following section we address the case where resources are also consumed by existing biomass for maintenance.

If we are concerned only with the relative dynamics of the two resources, then we can simplify Eq. S56 to describe dynamics in the two-dimensional space of resource concentrations:

$$
\frac{dR_2}{dR_1} = \frac{Y_1(R_1, R_2)}{Y_2(R_1, R_2)}. \tag{S57}
$$

We obtain this mathematically by taking the ratio of the last two lines in Eq. S56 and then changing variables from $t$ to $R_1$, which is possible because both resource concentrations $R_1$ and $R_2$ must depend monotonically on time $t$ since the resources are only consumed and not produced in this model. The right-hand side of Eq. S57 is the ratio of biomass yields for the two resources, which is the stoichiometry of those two resources in the biomass. For example, if every unit of biomass requires 3 units of resource 1 and 4 units of resource 2, then the stoichiometric ratio in Eq. S57 is 4/3.

This ratio thus defines the slope of depletion trajectories in the space of resource concentrations, as in Fig. 1D. Therefore depletion trajectories will be straight lines if the stoichiometry is constant over resource concentrations. However, if the stoichiometry is variable over concentrations, then the trajectories can curve. Let

$$
R_2 = f(R_1) \tag{S58}
$$

be any solution to Eq. S57. The solution for a specific set of initial concentrations $R_1(0)$ and $R_2(0)$ is therefore

$$
R_2 = R_2(0) + f(R_1) - f(R_1(0)). \tag{S59}
$$

For example, in the case where stoichiometry is constant, then $f(R_1)$ is a linear function of $R_1$ and so

$$
R_2 = R_2(0) - \frac{Y_1}{Y_2}\left(R_1(0) - R_1\right). \tag{S60}
$$

The total biomass yield of the population is the amount of new biomass produced in the infinite time limit:

$$
\begin{aligned}
\lim_{t\to\infty} N(t) - N(0) &= \Delta N \\
&= \int_0^{\infty} dt\ \frac{d}{dt}N(t) \\
&= -\int_{R_1(0)}^{\lim_{t\to\infty} R_1(t)} dR_1\ Y_1(R_1, R_2(R_1)) \\
&= -\int_{R_2(0)}^{\lim_{t\to\infty} R_2(t)} dR_2\ Y_2(R_1(R_2), R_2),
\end{aligned}
\tag{S61}
$$

where the latter two equations are obtained from the first by changing variables from time $t$ to either resource concentration $R_1$ or $R_2$ using Eq. S56 (which again is valid since these quantities all depend monotonically with each other). To calculate this biomass yield, we therefore need to know the limit of either $R_1$ or $R_2$ in the infinite time limit, which depends on how growth rate depends on these resources.

#### A. Growth stops only when at least one resource reaches zero concentration

The simplest case is where growth only stops when one or both resources reach zero concentration (Fig. 1D); this is the case for all the growth rate models discussed in SI Appendix, section S3, where $g(R_1, R_2) = 0$ if and only if $R_1 = 0$ or $R_2 = 0$. In this case, $\lim_{t\to\infty} R_1(t) = 0$ or $\lim_{t\to\infty} R_2(t) = 0$. Which limit is realized depends on the initial conditions $R_1(0)$ and $R_2(0)$. If we imagine the phase portrait of Eq. S57 (Fig. S4), this problem is geometrically realized as determining which trajectories terminate at the vertical $R_1 = 0$ axis (the population runs out of resource 1 first) or at the horizontal $R_2 = 0$ axis (the population runs out of resource 2 first).

There is only one trajectory that hits the origin $(R_1, R_2) = (0, 0)$. This initial conditions that follow this trajectory are defined by $R_2(0) = f(R_1(0)) - f(0)$ (using Eq. S59). Because trajectories cannot cross each other, all trajectories above this one will end at the $R_1 = 0$ axis, while all trajectories below will end at the $R_2 = 0$ axis. Define the former region as $\Omega_1$ and the latter region as $\Omega_2$, or more precisely,

$$\begin{aligned}\Omega_1 &= \{(R_1(0), R_2(0)) \text{ such that } R_2(0) > f(R_1(0)) - f(0)\},\\ \Omega_2 &= \{(R_1(0), R_2(0)) \text{ such that } R_2(0) < f(R_1(0)) - f(0)\}.\end{aligned} \tag{S62}$$

For example, in the case of constant stoichiometry, the set of initial conditions that follow the trajectory ending at the origin is given by the straight line $R_2(0) = (Y_1/Y_2)R_1(0)$ (Fig. S4A). So all initial conditions above this line ($R_2(0) > (Y_1/Y_2)R_1(0)$, region $\Omega_1$) will follow trajectories terminating at $R_1 = 0$, while all trajectories below ($R_2(0) < (Y_1/Y_2)R_1(0)$, region $\Omega_2$) will terminate at $R_2 = 0$.

We can now determine the resource concentration limits $\lim_{t\to\infty} R_1(t)$ and $\lim_{t\to\infty} R_2(t)$, and in turn the yield of the biomass, in terms of the initial conditions $R_1(0)$, $R_2(0)$:

$$\Delta N = \begin{cases} \displaystyle\int_0^{R_1(0)} dR_1 \; Y_1\left(R_1, R_2(0) + f(R_1) - f(R_1(0))\right) & \text{if } (R_1(0), R_2(0)) \in \Omega_1, \\ \displaystyle\int_0^{R_2(0)} dR_2 \; Y_2\left(R_1(0) + f^{-1}(R_2) - f^{-1}(R_2(0)), R_2\right) & \text{if } (R_1(0), R_2(0)) \in \Omega_2. \end{cases} \tag{S63}$$

That is, we choose the resource integral from Eq. S61 based on which resource goes to zero (since that simplifies the limits of the integral) and use the solution to the resource dynamics ODE to express the integral fully in terms of that one resource (using Eq. S59). In the special case of constant stoichiometry, Eq. S63 simplifies to

$$\begin{aligned}\Delta N &= \begin{cases} R_1(0)Y_1 & \text{if } R_2(0) > (Y_1/Y_2)R_1(0), \\ R_2(0)Y_2 & \text{if } R_2(0) < (Y_1/Y_2)R_1(0), \end{cases}\\ &= \min\left(R_1(0)Y_1, R_2(0)Y_2\right).\end{aligned} \tag{S64}$$

That is, the yield equals the biomass produced by whichever resource is initially less, according to its yield.

Equation S63 also says that for initial conditions in $\Omega_1$, the biomass yield always depends on $R_1(0)$ (as expected since resource 1 runs out at the end), but it can also depend on the initial resource 2 $R_2(0)$ if the yield for resource 1 depends on the concentration of resource 2. The reverse is true for initial conditions in $\Omega_2$. Thus, for the biomass yield to depend on both resources simultaneously, the yield for one resource must depend on the the concentration of the other resource, which we interpret as a form of interaction between the resources.

#### B. Growth stops at some nonzero combinations of resource concentrations

It is also possible that growth could stop before any resources reach zero concentration, i.e., when they reach some nonzero concentrations $R_i^* > 0$ for each resource $i$ [17]. For example, this could occur because birth has to balance a nonzero death rate, as in a chemostat; it could also happen because of some cellular regulation that stops growth at nonzero concentrations, perhaps in preparation for starvation. If these concentrations are independent of each other, then the dynamics of resource depletion are the same as in the previous section, but with a shift in all resource concentrations by these thresholds $R_i^*$.

The more complicated situation occurs if the thresholds $R_i^*$ depend on each other, i.e., they form a curved line in the space of $R_1$ and $R_2$ (Fig. S4A). We can think of this as also representing a form of interaction between resources. We determine the final concentration of resources therefore by calculating the intersection of the resource trajectory given by Eq. S59 with the curve of thresholds $R_1^*$ and $R_2^*$. As an example, we focus on the case with constant yields $Y_1$ and $Y_2$. Let us assume the birth rate follows the additive model (Eq. S31) but the net growth rate contains a nonzero death rate $d$:

$$g(R_1, R_2) = \frac{g_{\max}}{1 + K_1/R_1 + K_2/R_2} - d. \tag{S65}$$

Therefore the thresholds $R_1^*$ and $R_2^*$ at which growth ceases are determined by the following equations:

$$\begin{aligned} 0 &= \frac{g_{\max}}{1 + K_1/R_1^* + K_2/R_2^*} - d, \\ R_2^* &= R_2(0) - \frac{Y_1}{Y_2}\left(R_1(0) - R_1^*\right). \end{aligned} \tag{S66}$$

We can solve these equations to obtain the final resource concentrations:

$$\begin{aligned} R_1^* &= \frac{y_1 + y_2 + (\gamma - 1)(y_1 r_1(0) - y_2 r_2(0)) + \sqrt{D}}{2y_1(\gamma - 1)} K_1, \\ R_2^* &= \frac{y_1 + y_2 + (\gamma - 1)(y_2 r_2(0) - y_1 r_1(0)) + \sqrt{D}}{2y_2(\gamma - 1)} K_2, \end{aligned} \tag{S67}$$

where

$$\begin{aligned} \gamma =& g_{\max}/d, \\ r_1(0) =& \frac{R_1(0)}{K_1}, \\ r_2(0) =& \frac{R_2(0)}{K_2}, \\ y_1 =& K_1 Y_1, \\ y_2 =& K_2 Y_2, \\ D =& y_1^2(1 + r_1(0)[r_1(0)(\gamma - 1) - 2][\gamma - 1]) \\ &+ y_2^2(1 + r_2(0)[r_2(0)(\gamma - 1) - 2][\gamma - 1]) \\ &+ 2y_1y_2(1 - r_1(0) - r_2(0) - r_1(0)r_2(0) + [r_1(0) + r_2(0) + 2r_1(0)r_2(0)]\gamma - r_1(0)r_2(0)\gamma^2). \end{aligned} \tag{S68}$$

The total biomass yield at the end of growth is therefore

$$\begin{aligned} \Delta N &= (R_1(0) - R_1^*)Y_1, \\ &= (R_2(0) - R_2^*)Y_2. \end{aligned} \tag{S69}$$

We see that even in this case where yields are constant, the biomass yield depends on both initial resource concentrations $R_1(0)$ and $R_2(0)$ (Figs. S4C and S5A,B).

### C. Resource depletion dynamics and biomass yield with resource consumption for maintenance

#### 1. *Case of one resource*

In this section we consider the possibility of resources being consumed by existing biomass for maintenance, as well as by new biomass for growth. We first consider the case of a single resource. Modifying Eq. S56, the dynamics of biomass growth and resource consumption are given by

$$\frac{d}{dt}N(t) = g(R(t))N(t),$$
$$\frac{d}{dt}R(t) = \left(-\frac{1}{Y_{\text{growth}}}\frac{d}{dt}N(t) - \frac{1}{Y_{\text{main}}}N(t)\right)\Theta\left(R(t)\right), \tag{S70}$$

where $Y_{\text{growth}}$ is the "growth yield" (amount of new biomass produced per unit resource consumed) and $Y_{\text{main}}$ is the "maintenance yield" (amount of existing biomass maintained per unit resource consumed per unit time). For simplicity, we assume these are constant parameters and do not also depend on the resource concentrations as in the previous section. The Heaviside step function $\Theta$ is necessary so that once resources reach zero, there is no additional consumption due to maintenance (otherwise resource concentrations will become negative). Presumably the existing biomass will start to decay at some point after that, but we do not model it here.

We can express the second line of Eq. S70 in terms of an effective growth yield

$$\frac{d}{dt}R(t) = -\frac{1}{\tilde{Y}_{\text{growth}}(R(t))}\frac{d}{dt}N(t) \tag{S71}$$

where

$$\tilde{Y}_{\text{growth}}(R) = \left(\frac{1}{Y_{\text{growth}}} + \frac{1}{Y_{\text{main}}g(R)}\right)^{-1}. \tag{S72}$$

This predicts a positive but nonlinear correlation between growth rate and yield, across varying resource concentrations $R$ [18, 19]. In particular, when the growth rate is zero, the effective growth yield is also zero, because all resource consumption goes to maintenance. When the growth rate is infinite, then the effective growth yield converges to the regular growth yield $Y_{\text{growth}}$, since maintenance consumption becomes negligible. If the growth rate is described by a Monod model $g(R) = g_{\text{max}}R/(R+K)$, then the effective growth yield becomes

$$\tilde{Y}_{\text{growth}}(R) = \tilde{Y}_{\text{growth}}(\infty)\frac{R}{R + \frac{K\tilde{Y}_{\text{growth}}(\infty)}{g_{\text{max}}Y_{\text{main}}}}, \tag{S73}$$

where the effective growth yield at infinite resource is

$$\tilde{Y}_{\text{growth}}(\infty) = \left(\frac{1}{Y_{\text{growth}}} + \frac{1}{Y_{\text{main}}g_{\text{max}}}\right)^{-1}. \tag{S74}$$

Thus the effective growth yield depends as a Monod function on the resource concentration $R$, with a saturating yield at $\tilde{Y}_{\text{growth}}(\infty)$ (the growth yield at maximum growth rate) and a half-saturation concentration of $K\tilde{Y}_{\text{growth}}(\infty)/(g_{\text{max}}Y_{\text{main}})$. This therefore constitutes a model of variable stoichiometry: the resource consumption changes dynamically over a growth cycle.

What is the biomass yield? We can integrate Eq. S71 to get the total change in biomass:

$$\begin{aligned}\Delta N &= \int_0^{R(0)} dR\ \tilde{Y}_{\text{growth}}(R)\\ &= \tilde{Y}_{\text{growth}}(\infty)\left(R(0) + \frac{K\tilde{Y}_{\text{growth}}(\infty)}{g_{\text{max}}Y_{\text{main}}}\log\left[\frac{\frac{K\tilde{Y}_{\text{growth}}(\infty)}{g_{\text{max}}Y_{\text{main}}}}{\frac{K\tilde{Y}_{\text{growth}}(\infty)}{g_{\text{max}}Y_{\text{main}}} + R(0)}\right]\right).\end{aligned} \tag{S75}$$

For very large initial resource concentrations $R(0)$ this is approximately the regular biomass yield $\tilde{Y}_{\text{growth}}(\infty)R(0)$, since growth consumption dominates over maintenance in that case. The logarithm term is a correction at low initial resource concentrations $R(0)$ due to maintenance.

#### 2. Two resources

If there are two resources, the effective growth yields are

$$\begin{aligned} \tilde{Y}_{1,\text{growth}}(R_1, R_2) &= \left( \frac{1}{Y_{1,\text{growth}}} + \frac{1}{Y_{1,\text{main}} g(R_1, R_2)} \right)^{-1}, \\ \tilde{Y}_{2,\text{growth}}(R_1, R_2) &= \left( \frac{1}{Y_{2,\text{growth}}} + \frac{1}{Y_{2,\text{main}} g(R_1, R_2)} \right)^{-1}. \end{aligned} \tag{S76}$$

If we take the ratio to get the stoichiometry, we can then express the depletion trajectory of both resources as

$$\begin{aligned} \frac{dR_2}{dR_1} &= \frac{\tilde{Y}_{1,\text{growth}}(R_1, R_2)}{\tilde{Y}_{2,\text{growth}}(R_1, R_2)} \\ &= \frac{\left( \frac{1}{Y_{2,\text{growth}}} + \frac{1}{Y_{2,\text{main}} g(R_1, R_2)} \right)}{\left( \frac{1}{Y_{1,\text{growth}}} + \frac{1}{Y_{1,\text{main}} g(R_1, R_2)} \right)}. \end{aligned} \tag{S77}$$

Since the stoichiometry varies with the resource concentrations, this will lead to curved trajectories of resource depletion (Fig. S4). The stoichiometry only depends on the resources through the growth rate, we can characterize its behavior as a function of growth rate. That is, the stoichiometry interpolates between two values at different extremes of growth rate:

$$\frac{Y_1}{Y_2} \approx \begin{cases} \frac{Y_{1,\text{main}}}{Y_{2,\text{main}}} & \text{for slow growth (low resource concentrations),} \\ \\ \frac{\frac{1}{Y_{1,\text{growth}}} + \frac{1}{Y_{1,\text{main}} g_{\text{max}}}}{\frac{1}{Y_{2,\text{growth}}} + \frac{1}{Y_{2,\text{main}} g_{\text{max}}}} & \text{for fast growth (high resource concentrations).} \end{cases} \tag{S78}$$

We plot an example phase portrait of this using the additive model of growth rate (Eq. S31) in Fig. S5C.

Moreover, since the yield for each resource depends on the other resource's concentration (e.g., $Y_1$ depends on $R_2$ via the growth rate in Eq. S76), this will also produce biomass gains $\Delta N$ that always depend on both resource concentrations regardless of the initial conditions (Fig. S4D). Thus maintenance is a mechanism for non-Liebig biomass dependence.

### D. Resource depletion dynamics and biomass yield with dynamic proteome allocation

Besides maintenance, another plausible mechanism of variable stoichiometry is that cells regulate their stoichiometry based on their growth rate. For example, we know that the proteome allocation between different sectors of proteins changes significantly at different growth rates [20]. Suppose that cells consist of two biomass components, A and B. Let the yields of biomass component A on resources 1 and 2 be $Y_1^{\text{A}}$ and $Y_2^{\text{A}}$, while the yields of component B are $Y_1^{\text{B}}$ and $Y_2^{\text{B}}$. Thus the stoichiometry of component A is $Y_1^{\text{A}}/Y_2^{\text{A}}$ and the stoichiometry of B is $Y_1^{\text{B}}/Y_2^{\text{B}}$. We assume these stoichiometries are unequal, since otherwise the two biomass components are equivalent in their resource usage.

Let the physiological state variable $\theta$ parameterize the fraction of biomass in component A (so the fraction in component B is $1 - \theta$). The yields of both resources depend on this biomass state according to

$$\begin{aligned} Y_1 &= \left( \frac{\theta}{Y_1^{\text{A}}} + \frac{1-\theta}{Y_1^{\text{B}}} \right)^{-1}, \\ Y_2 &= \left( \frac{\theta}{Y_2^{\text{A}}} + \frac{1-\theta}{Y_2^{\text{B}}} \right)^{-1}. \end{aligned} \tag{S79}$$

We assume the physiological state is regulated by growth rate, such that the two are linearly proportional:

$$\theta(g) = \left(\frac{\theta_{\max} - \theta_{\min}}{g_{\max}}\right) g + \theta_{\min}, \tag{S80}$$

so that at zero growth, $\theta_{\min}$ is the fraction of component A in the biomass, while at maximum growth rate $g_{\max}$, $\theta_{\max}$ is the fraction of A. This is motivated by the linear relationship between growth rate and proteome sectors. Without loss of generality, we assume the shift is in favor of component A at high growth rates.

Since the growth rate depends on the resource concentrations $g(R_1, R_2)$ according to a model from SI Appendix, section S3, we arrive at a model with variable stoichiometry:

$$\begin{aligned}\frac{Y_1}{Y_2} &= \frac{\frac{1}{Y_2^{\mathrm{A}}}\theta(g(R_1,R_2)) + \frac{1}{Y_2^{\mathrm{B}}}\left[1-\theta(g(R_1,R_2))\right]}{\frac{1}{Y_1^{\mathrm{A}}}\theta(g(R_1,R_2)) + \frac{1}{Y_1^{\mathrm{B}}}\left[1-\theta(g(R_1,R_2))\right]} \\ &= \frac{\frac{1}{Y_2^{\mathrm{A}}}\left[\left(\frac{\theta_{\max}-\theta_{\min}}{g_{\max}}\right)g(R_1,R_2)+\theta_{\min}\right] + \frac{1}{Y_2^{\mathrm{B}}}\left[1-\left(\frac{\theta_{\max}-\theta_{\min}}{g_{\max}}\right)g(R_1,R_2)-\theta_{\min}\right]}{\frac{1}{Y_1^{\mathrm{A}}}\left[\left(\frac{\theta_{\max}-\theta_{\min}}{g_{\max}}\right)g(R_1,R_2)+\theta_{\min}\right] + \frac{1}{Y_1^{\mathrm{B}}}\left[1-\left(\frac{\theta_{\max}-\theta_{\min}}{g_{\max}}\right)g(R_1,R_2)-\theta_{\min}\right]}.\end{aligned} \tag{S81}$$

In Fig. S4E we plot an example phase portrait for this model. As with variable stoichiometry arising from maintenance, stoichiometry in this model interpolates between two values depending on the growth rate: at slow growth rates, the biomass is made mostly of component B, and so the stoichiometry is approximately that of component B ($Y_1^{\mathrm{B}}/Y_2^{\mathrm{B}}$), while at fast growth rates, the biomass is made mostly of A and therefore reflects that component's stoichiometry ($Y_1^{\mathrm{A}}/Y_2^{\mathrm{A}}$). Figure S4F shows how the biomass yield depends in a non-Liebig manner over initial resource concentrations.

### S6. EFFECT OF UNMEASURED RESOURCES ON LIMITATION AND COLIMITATION ESTIMATES

In Fig. 4B,C we estimate limitation coefficients and the number of limiting resources for many species in natural environments. One caveat of these estimates is that we do not account for other resources for which we have no data on their growth rate response or environmental concentrations. However, we can deduce how such unmeasured resources would quantitatively affect our estimates.

For example, we can deduce how the limitation coefficient $L_1^{\mathrm{rate}}$ of a measured resource 1 is affected by the concentration $R_2$ of an unmeasured resource 2 using the contours in the second row of Fig. S2. If the unmeasured resource is at high concentration $R_2$ relative to its threshold $K_2$, then it has little effect on the limitation coefficient of the measured resource (i.e., blue contours of $L_1^{\mathrm{rate}}$ depend only weakly on $R_2$ for high $R_2$) under the PAT and additive models, and for the Liebig and multiplicative Monod models it has no effect at all since the contours are vertical. If the unmeasured resource is at low concentration, then the true limitation coefficient for the measured resource will always be lower than calculated without the unmeasured resource for the PAT and additive models. For the Liebig model, the limitation coefficient for the measured resource will be zero if the unmeasured resource is sufficient scarce. For the multiplicative Monod model, the limitation coefficient for the measured resource remains independent of all other resource concentrations. Therefore, the limitation coefficients we calculate here are at worst upper bounds on the true limitation coefficients.

---


[1] S. F. M. Hart, D. Skelding, A. J. Waite, J. C. Burton, and W. Shou. High-throughput quantification of microbial birth and death dynamics using fluorescence microscopy. *Quant Biol*, 7:69–81, 2019.

[2] H. Moser. *The Dynamics of Bacterial Populations Maintained in the Chemostat.* Carnegie Institution of Washington, 1958.

[3] C. S. Holling. Some Characteristics of Simple Types of Predation and Parasitism I. *The Canadian Entomologist*, 91:358–369, 1959.

[4] R. V. O'Neill, D. L. DeAngelis, J. J. Pastor, B. J. Jackson, and W. M. Post. Multiple nutrient limitations in ecological models. *Ecol Modelling*, 46:147–163, 1989.

[5] M. Zinn, B. Witholt, and T. Egli. Dual nutrient limited growth: Models, experimental observations, and applications. *J Biotechnol*, 113:263–279, 2004.

[6] M. A. Saito, T. J. Goepfert, and J. T. Ritt. Some thoughts on the concept of colimitation: Three definitions and the importance of bioavailability. *Limnol Oceanogr*, 53:276–290, 2008.

[7] M. E. Muscarella and J. P. O'Dwyer. Species dynamics and interactions via metabolically informed consumer-resource models. *Theor Ecol*, 13:503–518, 2020.
[8] J. Tang and W. J. Riley. Finding Liebig's law of the minimum. *Ecol Appl*, 31:e02458, 2021.
[9] J. von Liebig. *Organic chemistry in its application to agriculture and physiology.* Taylor and Walton, London, 1840.
[10] J. R. Casey and M. J. Follows. A steady-state model of microbial acclimation to substrate limitation. *PLoS Comput Biol*, 16:e1008140, 2020.
[11] E. Sperfeld, D. Martin-Creuzburg, and A. Wacker. Multiple resource limitation theory applied to herbivorous consumers: Liebig's minimum rule vs. interactive co-limitation. *Ecol Lett*, 15:142–150, 2012.
[12] S. A. L. M. Kooijman. The synthesizing unit as model for the stoichiometric fusion and branching of metabolic fluxes. *Biophys Chem*, 73:179–188, 1998.
[13] T. Mankad and H. R. Bungay. Model for microbial growth with more than one limiting nutrient. *J Biotechnol*, 7:161–166, 1988.
[14] L. Xie, A. E. Yuan, and W. Shou. Simulations reveal challenges to artificial community selection and possible strategies for success. *PLoS Biol*, 17:e3000295, 2019.
[15] D. W. Cantrell and E. W. Weisstein. Power mean. From MathWorld–A Wolfram Web Resource. URL: `https://mathworld.wolfram.com/PowerMean.html`, 2023. Accessed: 2023-05-21.
[16] D. Gupta, S. Garlaschi, S. Suweis, S. Azaele, and A. Maritan. Effective resource competition model for species coexistence. *Phys Rev Lett*, 127:208101, 2021.
[17] D. Tilman. *Resource competition and community structure.* Princeton University Press, Princeton, NJ, 1982.
[18] S. J. Pirt. The maintenance energy of bacteria in growing cultures. *Proc R Soc B*, 163:224–231, 1965.
[19] D. A. Lipson. The complex relationship between microbial growth rate and yield and its implications for ecosystem processes. *Front Microbiol*, 6:615, 2015.
[20] M. Mori, Z. Zhang, A. Banaei-Esfahani, J.-B. Lalanne, H. Okano, B. C. Collins, A. Schmidt, O. T. Schubert, D.-S. Lee, G.-W. Li, R. Aebersold, T. Hwa, and C. Ludwig. From coarse to fine: the absolute Escherichia coli proteome under diverse growth conditions. *Mol Syst Biol*, 17:e9536, 2021.
[21] R. Kishony and S. Leibler. Environmental Stresses Can Alleviate the Average Deleterious Effect of Mutations. *J Biol*, 2:14, 2003.

### S7. SI FIGURES

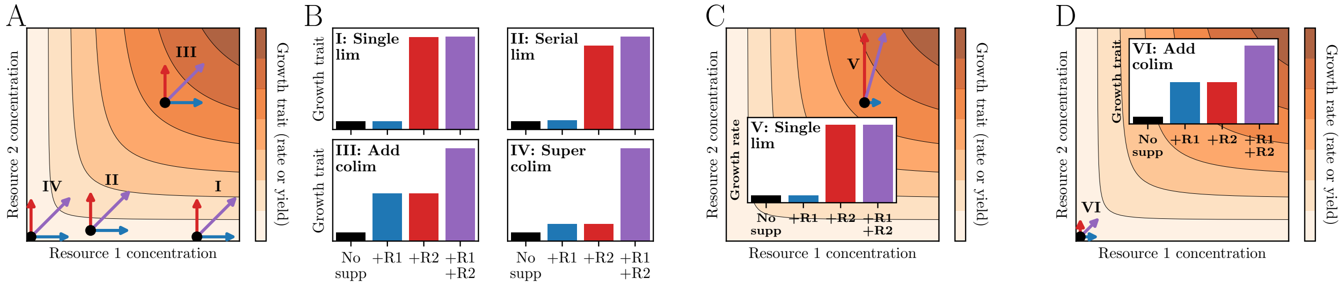


FIG. S1. **Factorial supplementation experiments are incomplete tests of colimitation.** (A) Schematic of four factorial supplementation experiments overlaid on the global dependence of a growth trait (e.g., growth rate or growth yield) on two resource concentrations. (B) Outcomes of the same four factorial supplementation experiments from (A), classified according to their apparent limitation outcome (single limitation, serial limitation, additive colimitation, super-additive colimitation). (C) Factorial supplementation experiment on the same background concentrations as experiment III from (A) and (B), but with different supplemented concentrations of both resources leading to a qualitatively different outcome (single limitation instead of additive colimitation). (D) Factorial supplementation experiment on the same background concentrations as experiment IV from (A) and (B), but with different supplemented concentrations of both resources leading to a qualitatively different outcome (additive colimitation instead of super-additive colimitation).

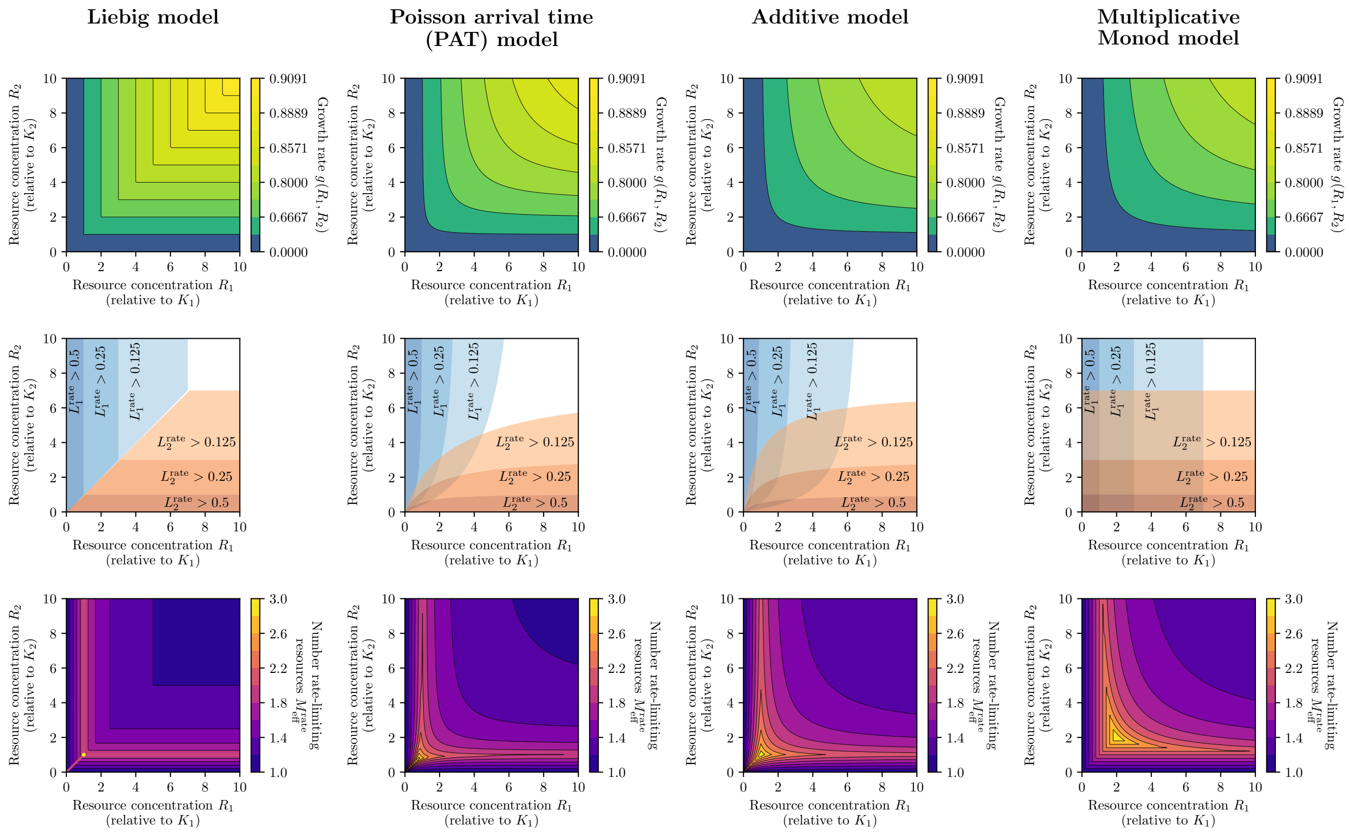


FIG. S2. **Summary of growth rate models and their limitation properties.** Each column corresponds to a different model of how growth rate depends on resource concentrations (SI Appendix, section S3). First row: For each growth rate model in a different column, we show the growth rates $g(R_1, R_2)$ as functions of resource concentrations $R_1$ and $R_2$. Second row: Rate limitation coefficients for resource 1 ($L_1^{\text{rate}}$, blue contours) and for resource 2 ($L_2^{\text{rate}}$, orange contours) as functions of resource concentrations (compare to schematic in Fig. 1B). Third row: Number of rate-limiting resources $M_{\text{eff}}^{\text{rate}}$ (Eq. 3) as functions of resource concentrations. All resource concentrations $R_i$ are scaled relative to their corresponding half-saturation concentrations $K_i = g_{\max}/a_i$, with $g_{\max} = 1$.

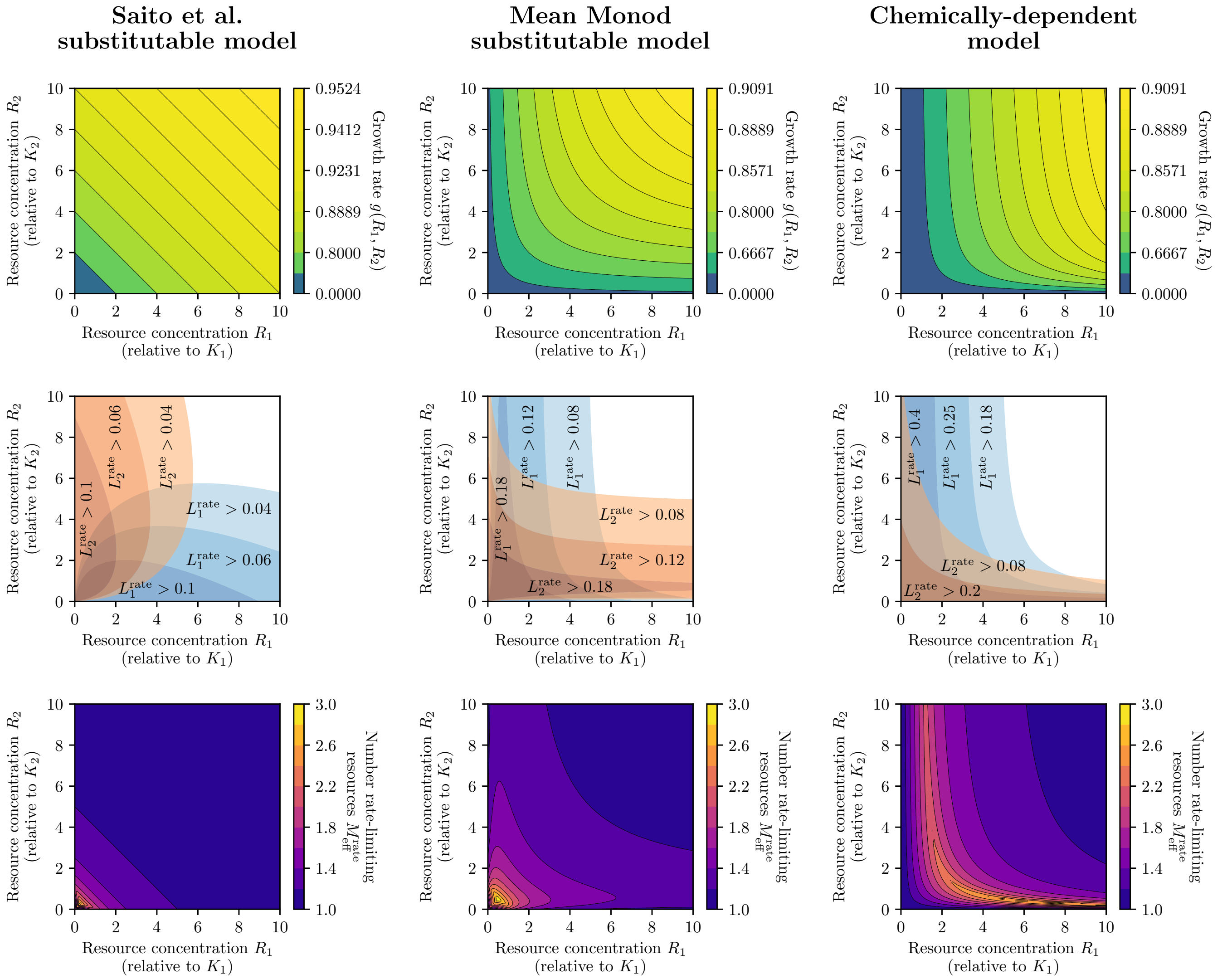


FIG. S3. **Summary of growth rate models and their limitation properties for substitutable and chemically-dependent resources.** Same as Fig. S2 but for the Saito et al. model [6] of substitutable resources (first column), the mean Monod model of substitutable resources (second column), and the model of chemically-dependent resources [6] (third column); see SI Appendix, sections S3 for definitions. All resource concentrations $R_i$ are scaled relative to their corresponding half-saturation concentrations $K_i = g_{\max}/a_i$, with $g_{\max} = 1$.

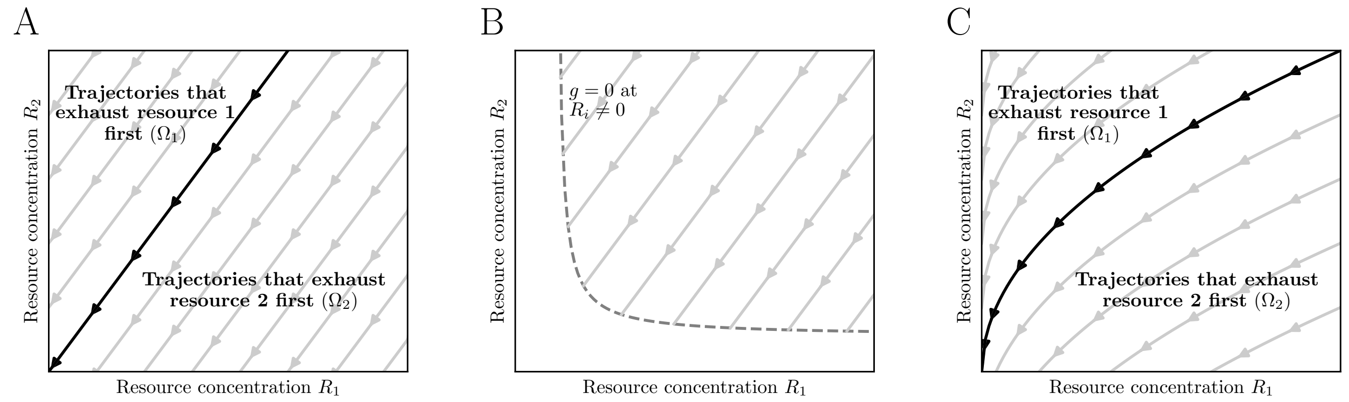


FIG. S4. **Different qualitative regimes of resource depletion trajectories.** (A) If resources are consumed with constant stoichiometry, then their concentrations deplete along straight lines (SI Appendix, section S5). The regions in which the first depleted resource is resource 1 or resource 2 are shown as $\Omega_1$ and $\Omega_2$. (B) Same as (A) for where resource depletion stops at some nonzero combination of concentrations (dashed gray line), e.g., when net growth rate reaches zero in a chemostat (SI Appendix, section S4). (C) Same as (A) but for resources consumed with variable stoichiometry, which causes their depletion trajectories to curve (SI Appendix, section S5).

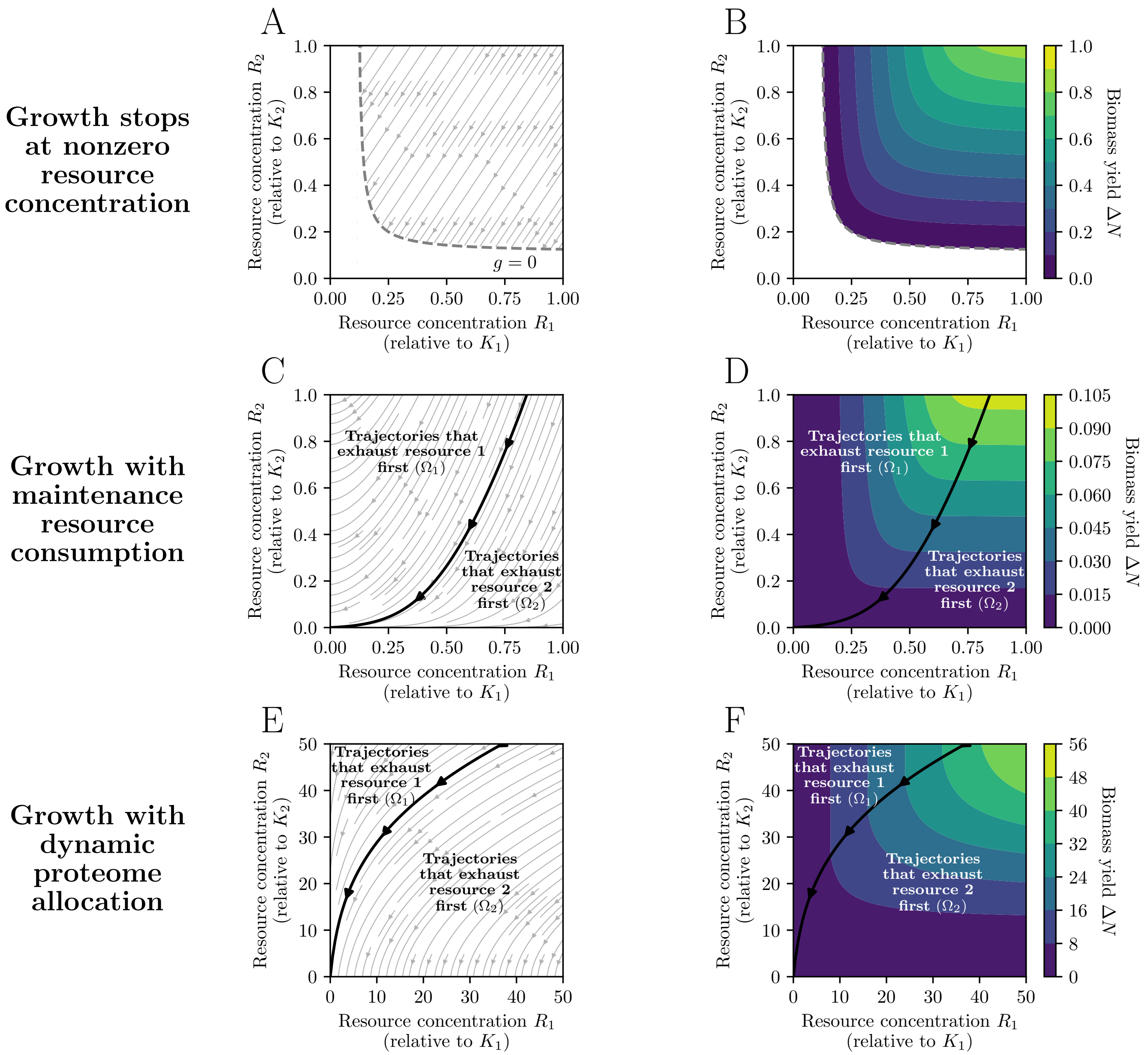


FIG. S5. **Yield limitation in different mechanistic models.** (A) Model of resource consumption in which growth stops at nonzero concentrations of resources (SI Appendix, section S5). These concentrations (zero net-growth isocline, ZNGI) are shown as the dashed gray line in the space of concentrations for resources 1 and 2. Gray stream lines show trajectories of resource depletion with constant stoichiometry. We use chemostat dynamics (SI Appendix, section S4) with the additive model of growth rate (SI Appendix, section S3) and parameter values $g_{\max} = 1$, $d = 0.1$, $a_1 = a_2 = 1$, $Y_1 = 1.5$, and $Y_2 = 1$. (B) For the model in (A), we calculate the growth yield $\Delta N$ after one batch growth cycle for all possible initial resource concentrations. (C) Model of resource consumption in which the population consumes resources for maintenance as well as for growth (SI Appendix, section S5). This causes stoichiometry to vary with resource concentration, and so depletion trajectories are curved. We use batch dynamics (SI Appendix, section S4) with the additive model of growth rate (SI Appendix, section S3) and parameter values $g_{\max} = 2$, $a_1 = a_2 = 1$, $Y_{1,\text{main}} = 0.5$, $Y_{2,\text{main}} = 10$, $Y_{1,\text{growth}} = 2$, and $Y_{2,\text{growth}} = 0.1$. (D) Same as (B) but for the maintenance model in (C). The regions in which the first depleted resource is resource 1 or resource 2 are shown as $\Omega_1$ and $\Omega_2$. (E) Model of resource consumption in which stoichiometry varies with growth rate according to changes in proteome allocation (SI Appendix, section S5). This causes stoichiometry to vary with resource concentration, and so depletion trajectories are curved. We use batch dynamics (SI Appendix, section S4) with the additive model of growth rate (SI Appendix, section S3) with parameter values $g_{\max} = 1$, $a_1 = a_2 = 1$, $\theta_{\min} = 0$, $\theta_{\max} = 1$, $Y_1^{\text{A}} = Y_1^{\text{B}} = 1$, $Y_2^{\text{A}} = 10$, and $Y_2^{\text{B}} = 0.1$. (F) Same as (D) but for the dynamic allocation model in (E).

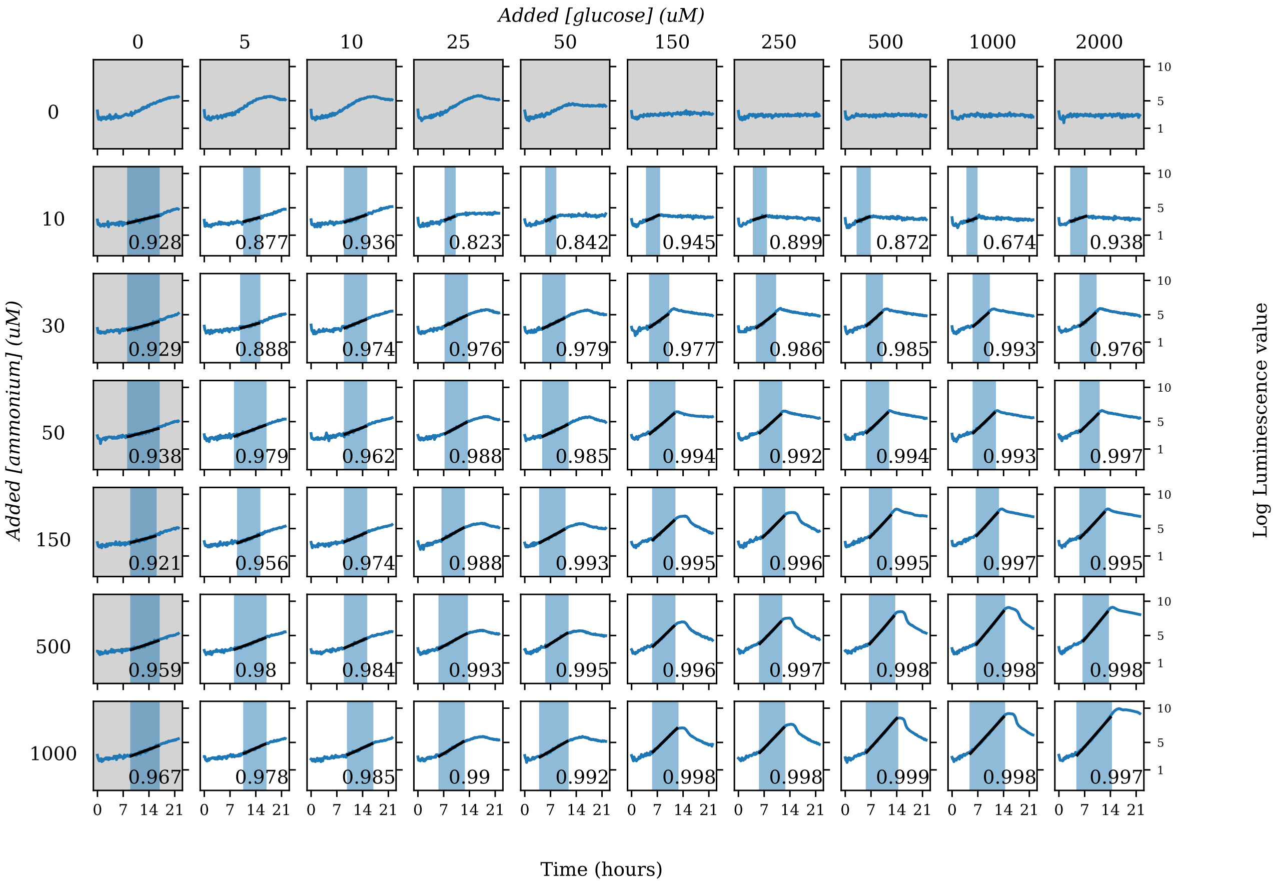


FIG. S6. **Luminescence growth curves across glucose and ammonium concentrations for biological replicate 1.** Growth curves of luminescent *E. coli* K-12 MG1655 pCS-$\lambda$ [21] with each well having a different starting concentration of glucose and ammonium. Data shown here is for replicate 1. The observed maximum growth rates, fitted from the data, are overlayed in black lines. The fitting interval (exponential growth phase) is shown in colored shading. Gray shading indicates wells where the added concentration of either glucose or ammonium was zero.

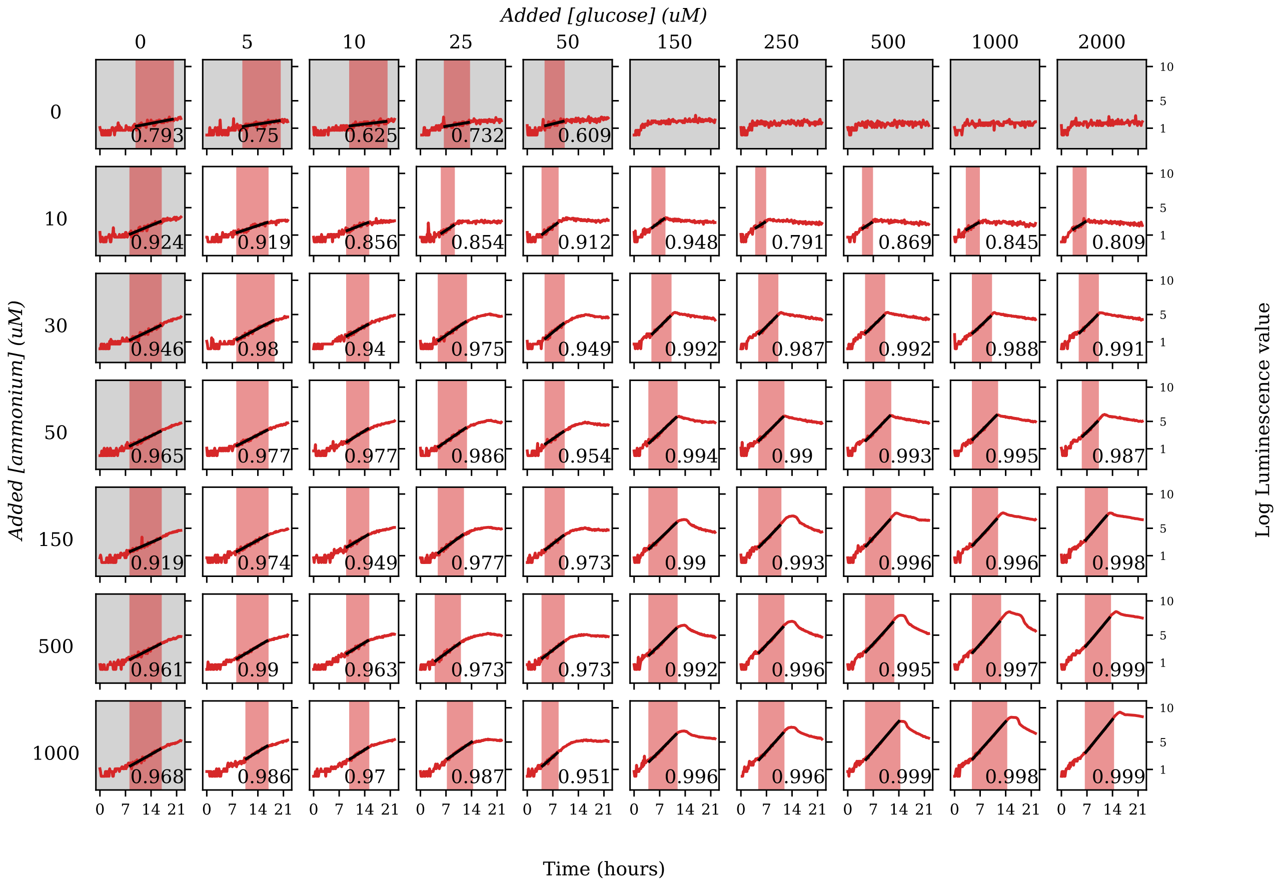


FIG. S7. **Luminescence growth curves across glucose and ammonium concentrations for biological replicate 2.** Same as Fig. S6 but for replicate 2.

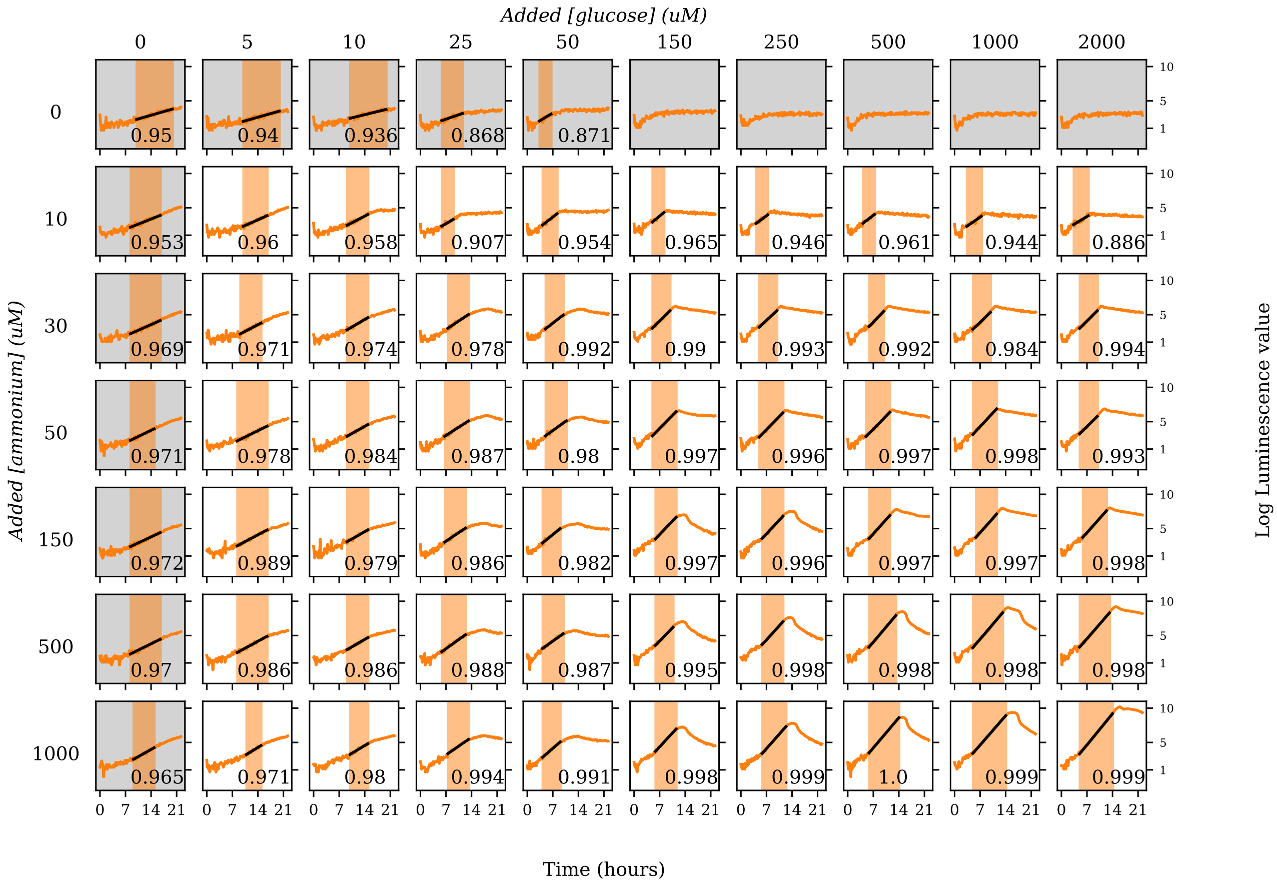


FIG. S8. **Luminescence growth curves across glucose and ammonium concentrations for biological replicate 3.** Same as Fig. S6 but for replicate 3.

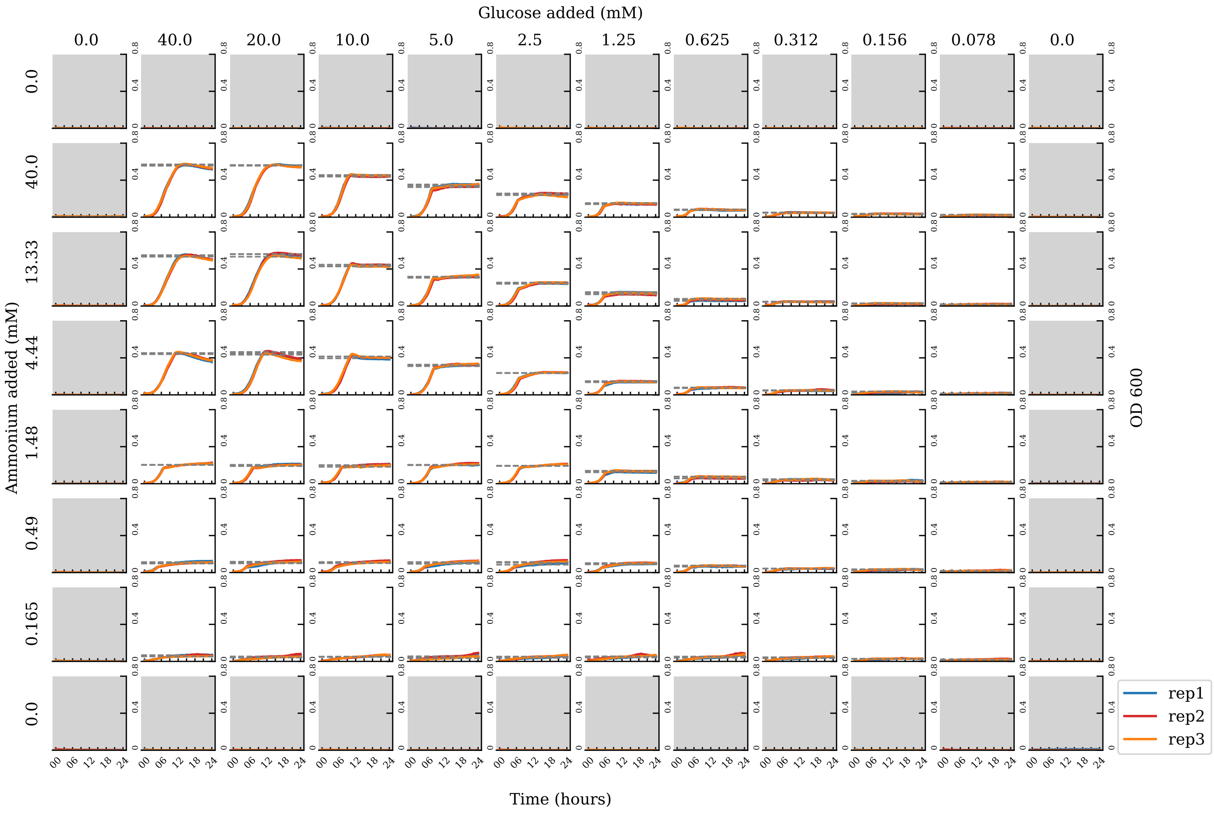


FIG. S9. **Optical density growth curves across glucose and ammonium concentrations.** Growth curves of OD at 600 nm for *E. coli* MG1655 with each well having a different starting concentration of glucose and ammonium. Triplicate growth curves are shown as colored lines; the observed growth yields, fitted from the data, are overlayed in gray dotted lines. Gray shading indicates wells were the concentration of either glucose or ammonium was zero. See Fig. S10 for the same data plotted on a log scale.

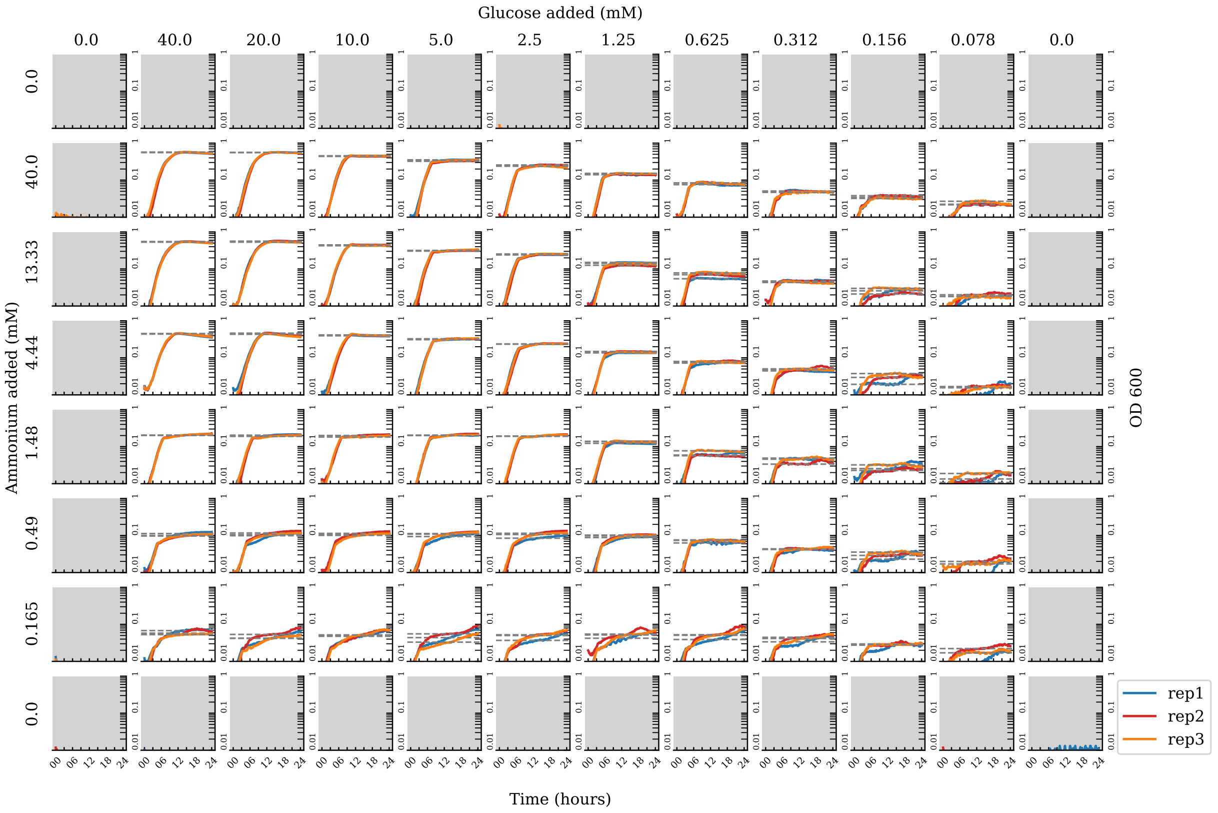


FIG. S10. **Log optical density growth curves across glucose and ammonium concentrations.** Same as Fig. S9 but with OD plotted on a logarithmic scale.

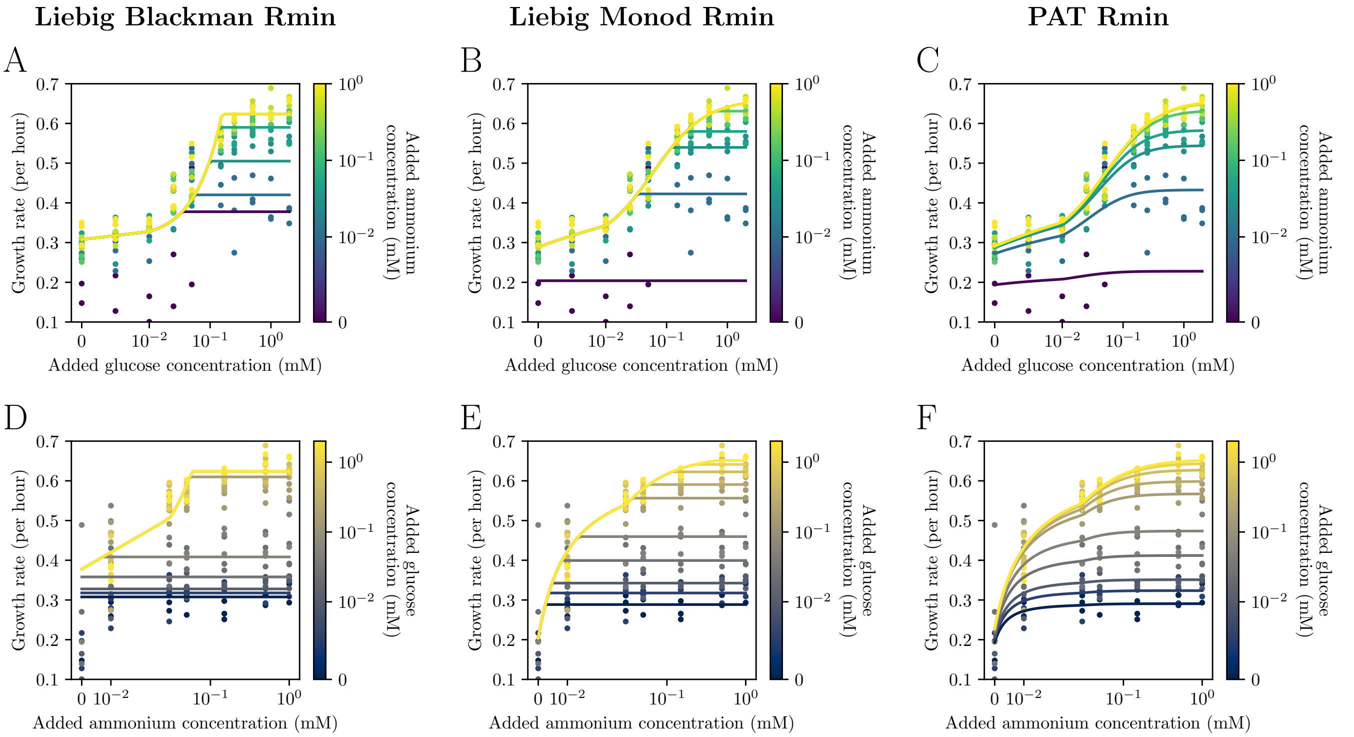


FIG. S11. **Example fits of growth rate scans to models.** Same as Fig. 2B but showing fits to the Liebig Blackman (A), Liebig Monod (B), and Poisson arrival time/synthesizing-unit (C) models, all with nonzero minimum resource concentration parameters (SI Appendix, section S3). Panels (D)–(F) are the same but projected onto the axis of ammonium concentrations, with colors showing glucose concentrations.

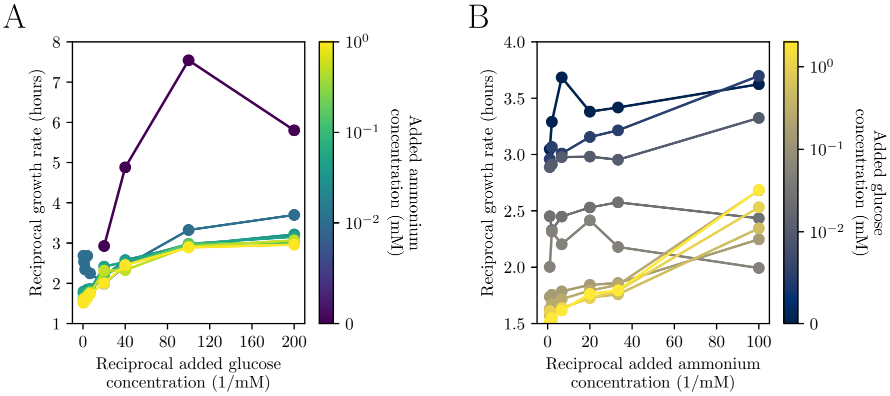


FIG. S12. **Lineweaver-Burke plots of growth rate vs. resource concentration.** Plot of reciprocal growth rate vs. reciprocal resource concentration, projected onto glucose concentrations (A) and ammonium concentrations (B), with colors in each representing the other resource concentration. Monod dependence appears as a straight line plotted this way. Points are averages over three experimental replicates.

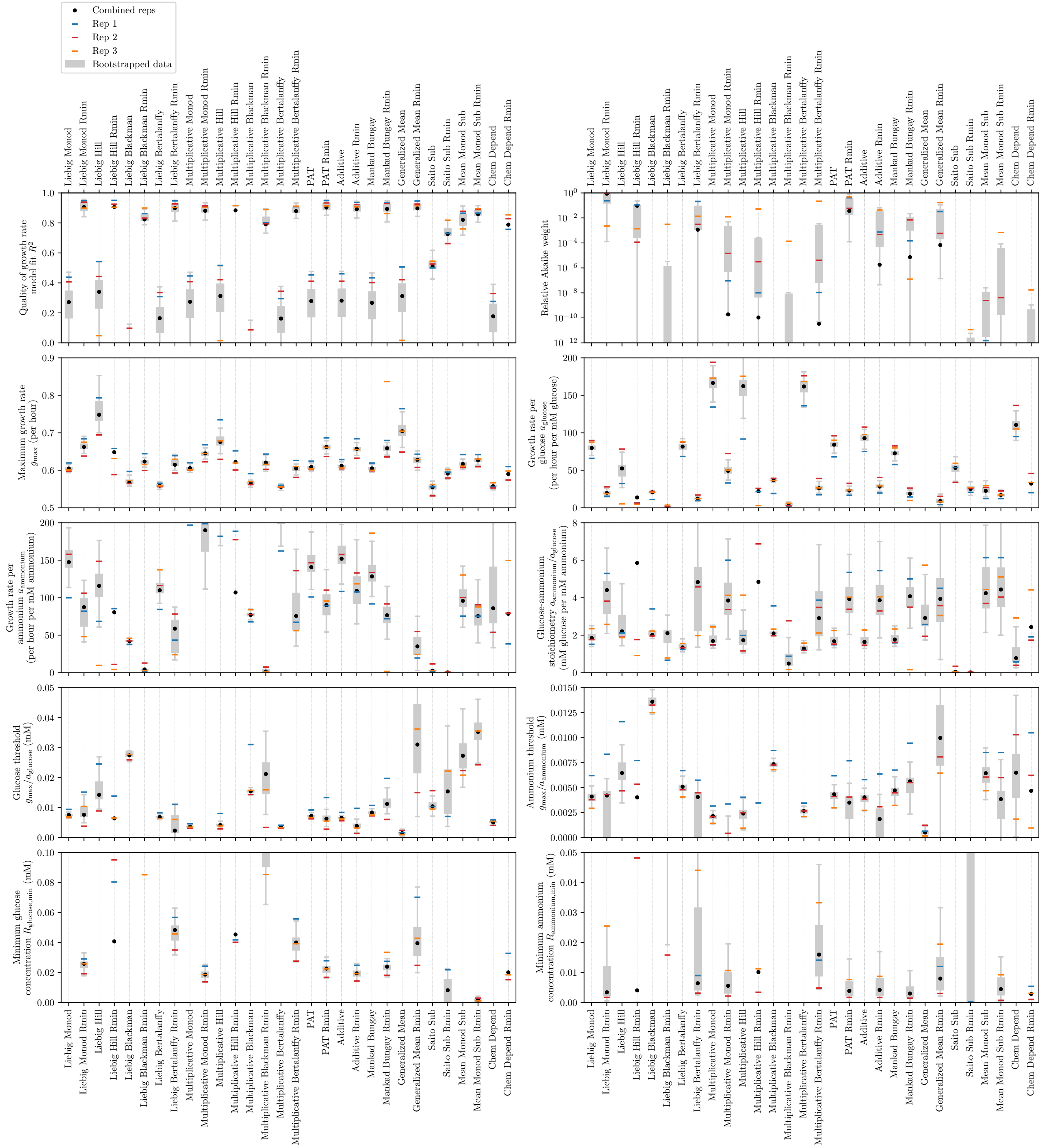

FIG. S13. **Fits of growth rate scans to models.** Each panel shows a different statistic from fitting the growth rate scan data to all models (across horizontal axes). The black dots mark statistics for fits to all three replicates together; the three colored bars show the statistics from the three replicates fitted separately. The gray boxes (first to third quartiles, with the whiskers extending to 1.5 times the interquartile range above and below) show the distributions of these statistics across 100 data sets bootstrapped from the three replicates (Materials and Methods). The thin vertical lines are to guide the eye to the model names on the bottom and top horizontal axes.

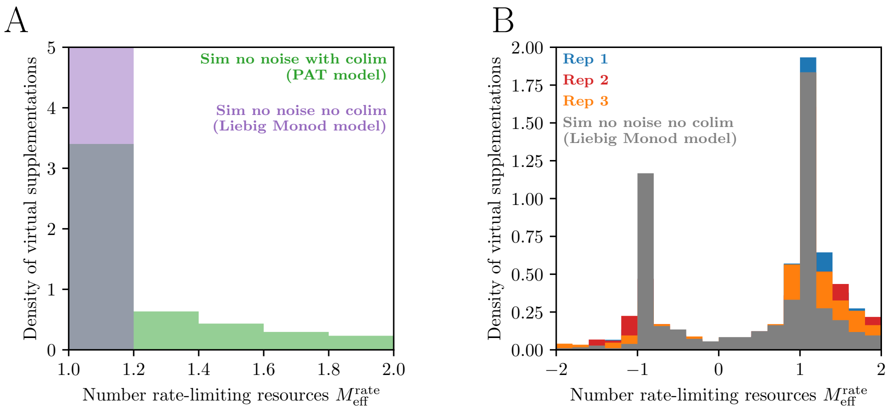


FIG. S14. **Growth rate colimitation across virtual supplementation experiments.** (A) Distribution of $M_{\mathrm{eff}}^{\mathrm{rate}}$ across virtual supplementation experiments for simulated growth rate models without colimitation (purple, Liebig Monod model; SI Appendix, section S3) and with colimitation (green, Poisson arrival time/synthesizing unit model; SI Appendix, section S3), in the absence of noise. (B) Distribution of $M_{\mathrm{eff}}^{\mathrm{rate}}$ across virtual supplementation experiments for experimental replicates (blue, red, and green) of growth rate measurements and for one simulated data set with noise (gray; Liebig Monod model, SI Appendix, section S3).

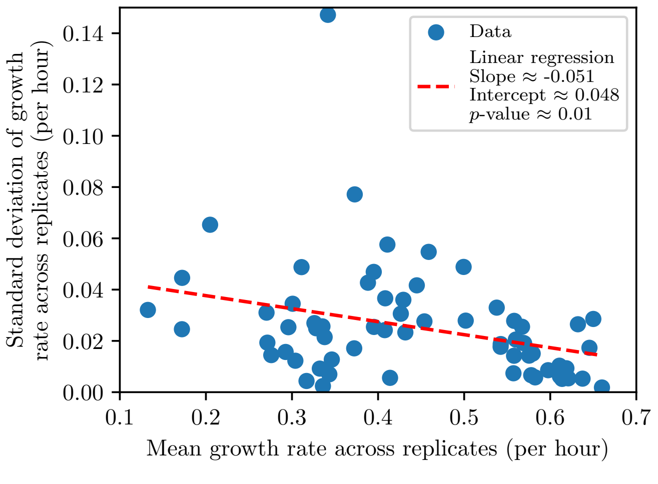


FIG. S15. **Variation of growth rate measurements across replicates.** Scaling of mean rate vs. standard deviation of rate across three experimental replicates (blue points). The red dashed line is the linear regression with parameters shown in the legend.

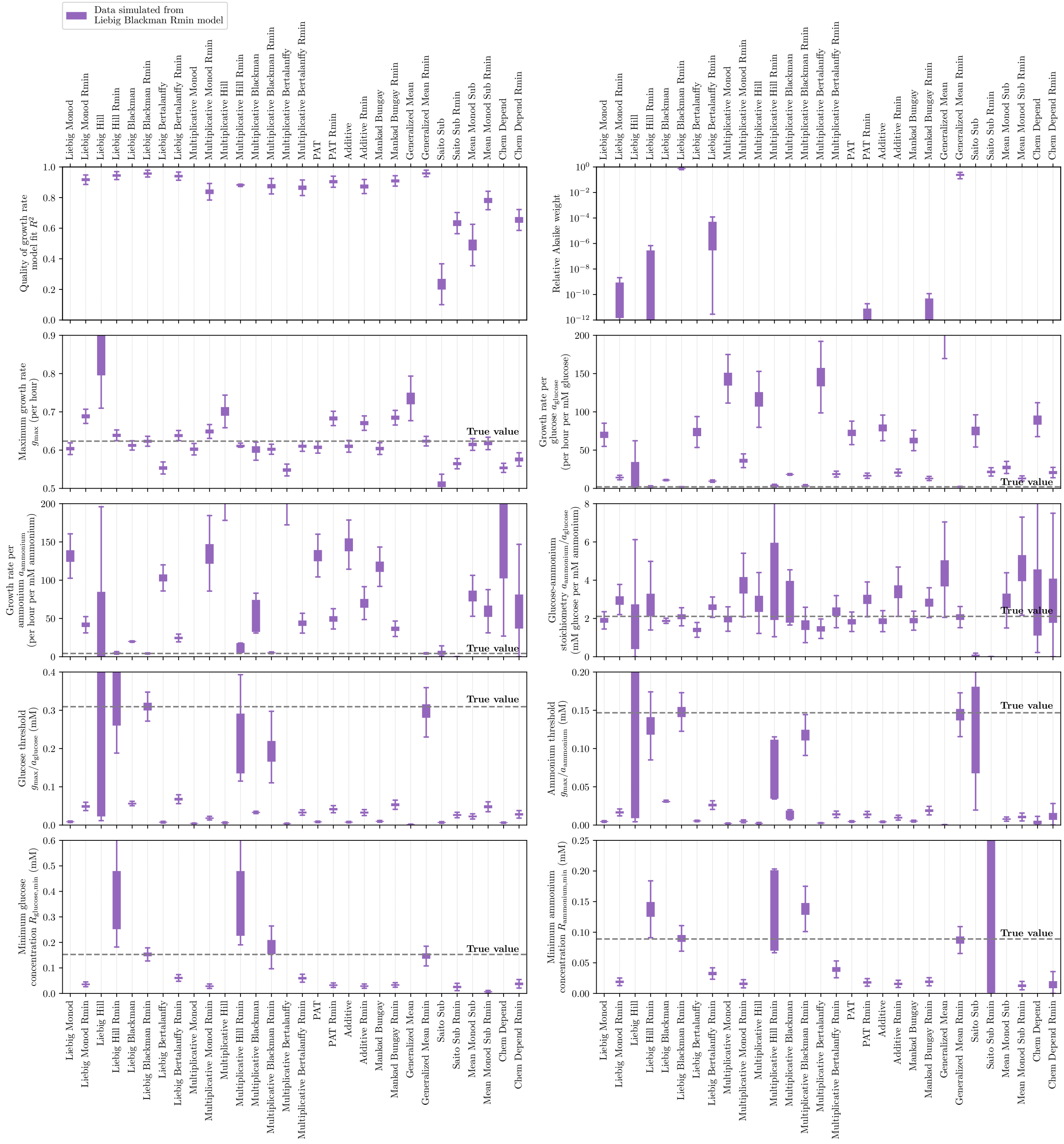


FIG. S16. **Fits of simulated (Liebig Blackman model) growth rate scans to models.** Similar to Fig. S13 but for fits to data simulated from the Liebig Blackman model (Materials and Methods; SI Appendix, section S3) with parameters from that model's fit to the experimental growth rate data ($g_{\max} \approx 0.62$ per hour, $a_{\text{glu}} \approx 2.01$ per hour per mM glucose, $a_{\text{amm}} \approx 4.2$ per hour per mM ammonium; $R_{\text{glu,min}} \approx 0.15$ mM, $R_{\text{amm,min}} \approx 0.089$ mM). We incorporated experimental noise to the simulations by adding a Gaussian-distributed random number to each measurement with mean zero and standard deviation that is a linear function of the mean according to the linear regression in Fig. S15. The purple boxes (first to third quartiles, with the whiskers extending to 1.5 times the interquartile range above and below) show the distributions of these statistics across $10^4$ simulated data sets. The true values of the parameters used in the simulations are marked by horizontal dashed lines.

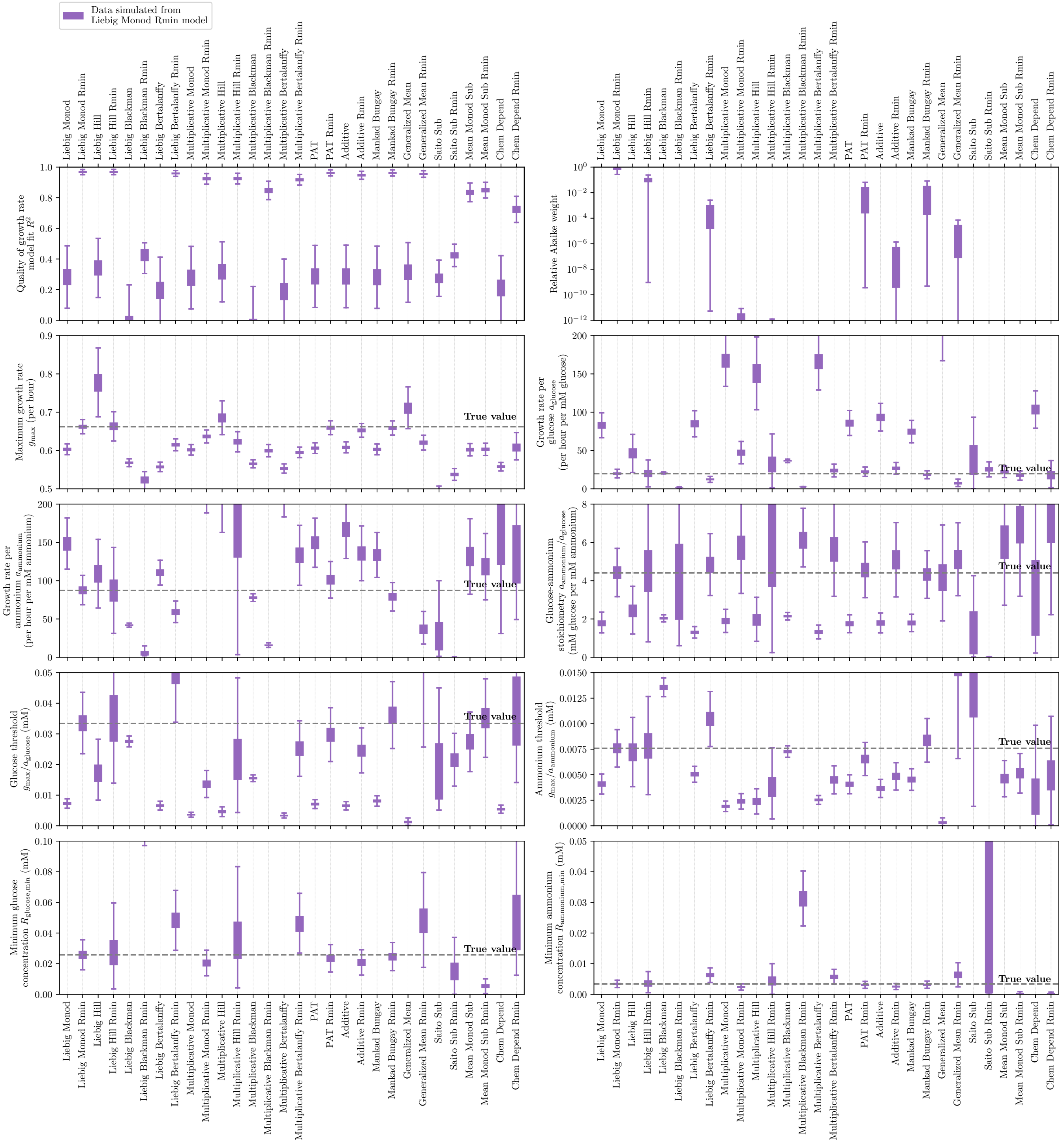


FIG. S17. **Fits of simulated (Liebig Monod model) growth rate scans to models.** Same as Fig. S16 but for $10^4$ simulations of the Liebig Monod model (Materials and Methods; SI Appendix, section S3) with parameters from that model's fit to the experimental growth rate data ($g_{\max} \approx 0.66$ per hour, $a_{\text{glu}} \approx 20$ per hour per mM glucose, $a_{\text{amm}} \approx 87$ per hour per mM ammonium; $R_{\text{glu,min}} \approx 0.026$ mM, $R_{\text{amm,min}} \approx 0.0034$ mM).

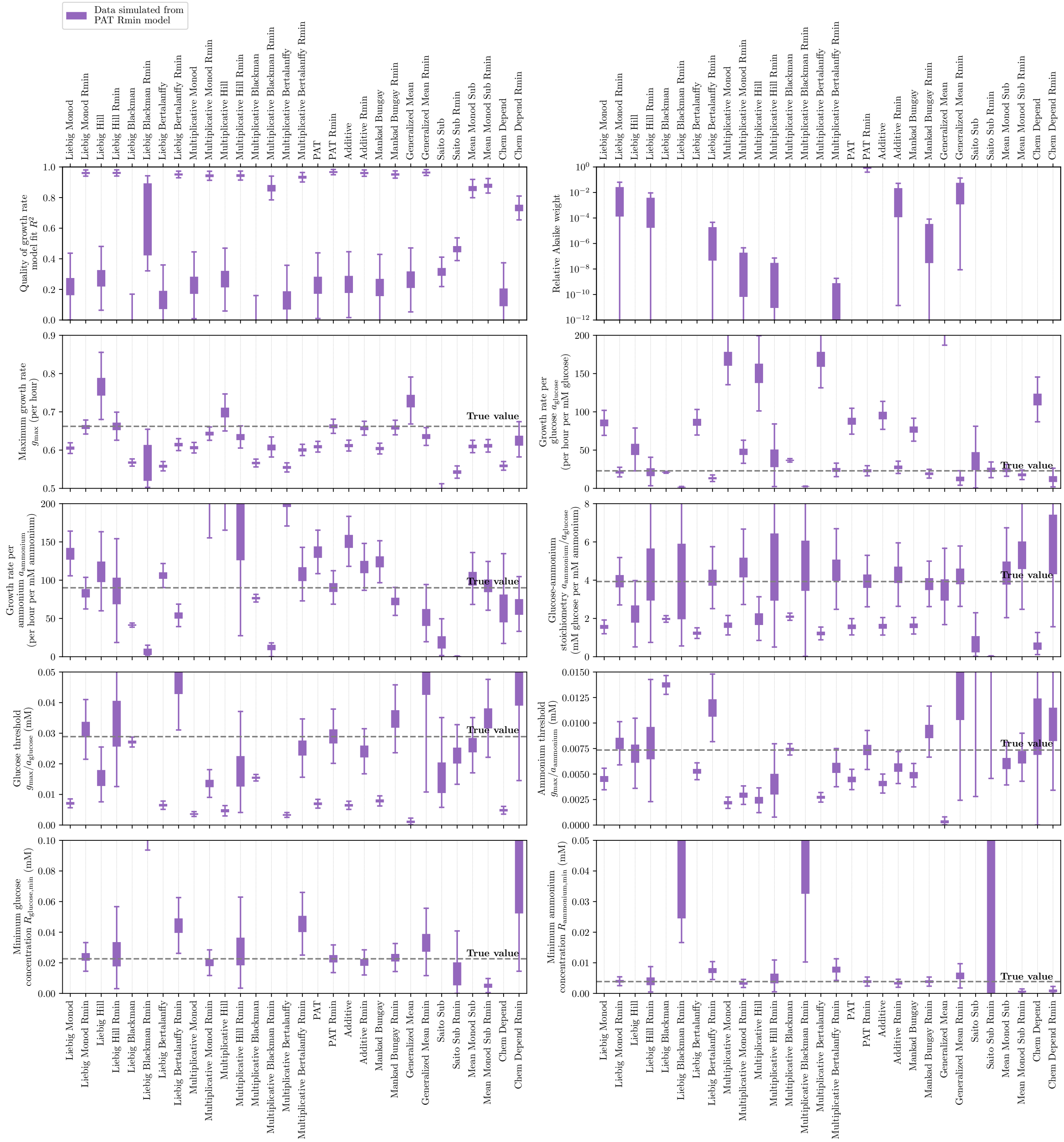


FIG. S18. **Fits of simulated (PAT model) growth rate scans to models.** Same as Fig. S16 but for $10^4$ simulations of the Poisson arrival time/synthesizing-unit model (Materials and Methods; SI Appendix, section S3) with parameters from that model's fit to the experimental growth rate data ($g_{\max} \approx 0.66$ per hour, $a_{\text{glu}} \approx 23$ per hour per mM glucose, $a_{\text{amm}} \approx 90$ per hour per mM ammonium; $R_{\text{glu,min}} \approx 0.023$ mM, $R_{\text{amm,min}} \approx 0.0039$ mM).

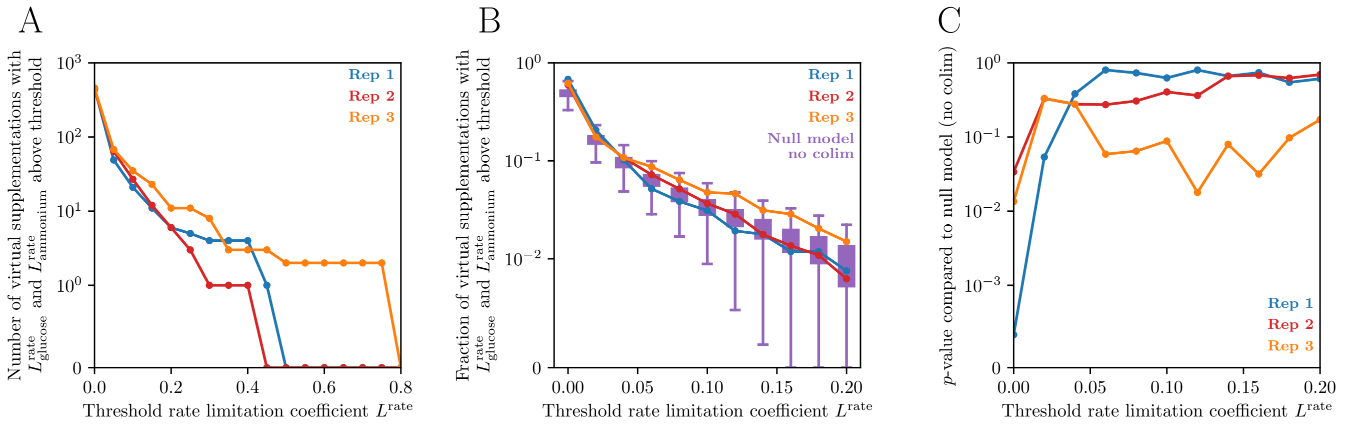


FIG. S19. **Testing significance of growth rate colimitation at different thresholds.** (A) Number of virtual supplementation experiments (Materials and Methods) with growth rate limitation coefficients for both glucose and ammonium above a given threshold, as a function of that threshold. (B) Fraction of virtual supplementation experiments with growth rate limitation coefficients for both glucose and ammonium above a given threshold, as a function of that threshold. Colored dots represent of each of the three experimental replicates, and the purple boxes show distributions across $10^4$ data sets simulated from a model with no colimitation (Liebig Monod model; Materials and Methods; SI Appendix, section S3). (C) $p$-values of the observed fractions of virtual supplementations with growth rate colimitation as functions of the limitation threshold.

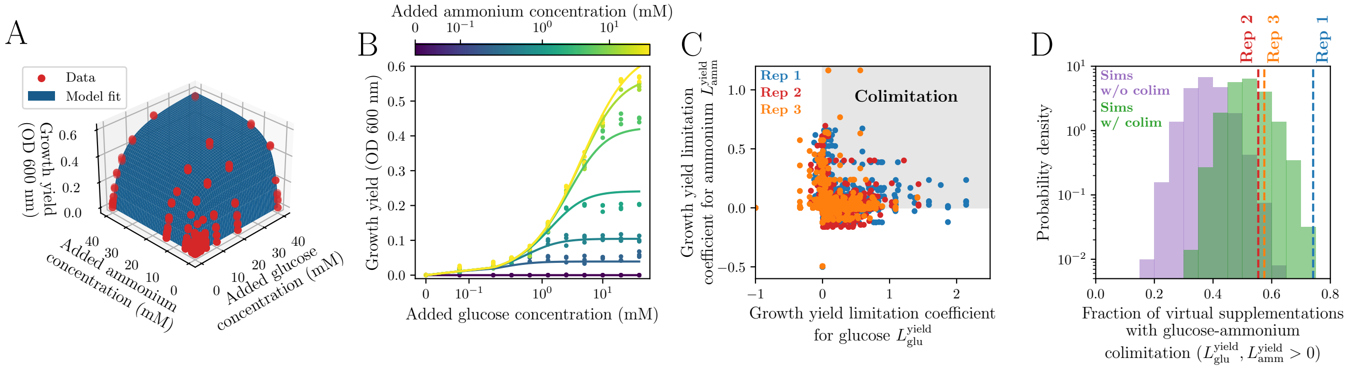


FIG. S20. **Measuring growth yield colimitation in laboratory conditions.** Same as Fig. 2 but for growth yield data (Dataset S2; see Figs. S9 and S10 for growth curves). In (A) the blue surface is a model fit to all replicate measurements (Poisson arrival time/synthesizing-unit model with $N_{\mathrm{max}} \approx 0.68$ OD, $a_{\mathrm{glu}} \approx 0.15$ OD/mM glucose, and $a_{\mathrm{amm}} \approx 0.25$ OD/mM ammonium; Materials and Methods; Dataset S2). The $p$-values in (D) are $p = 0$ for replicate 1, $p = 0.0032$ for replicate 2, and $p = 0.0012$ for replicate 3.

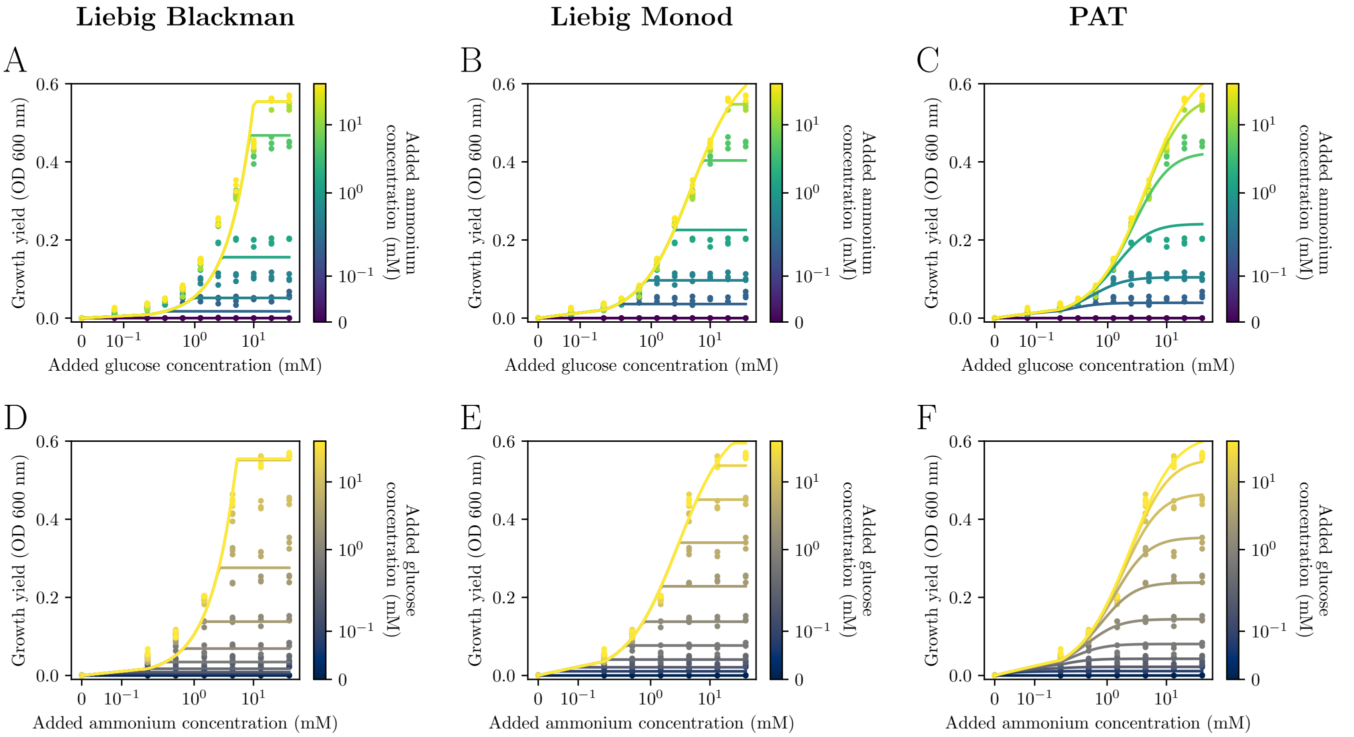


FIG. S21. **Example fits of growth yield scans to models.** Same as Fig. S11 but for growth yield data.

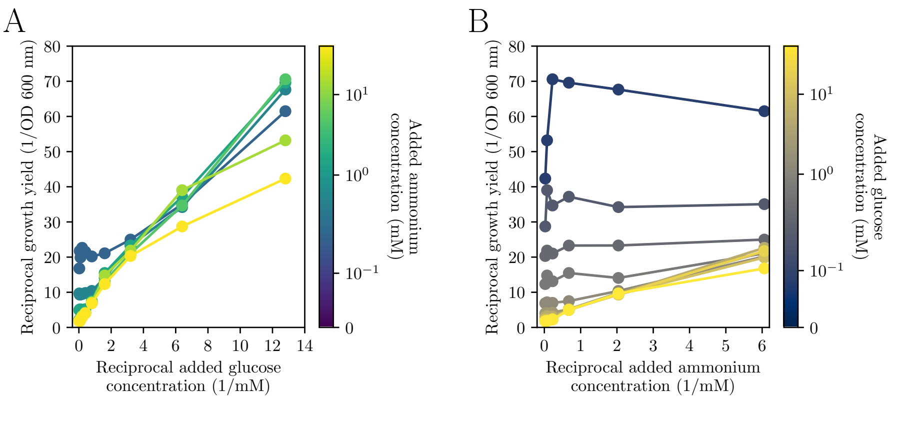


FIG. S22. **Lineweaver-Burke plots of growth yield vs. resource concentration.** Same as Fig. S12 but for growth yield.

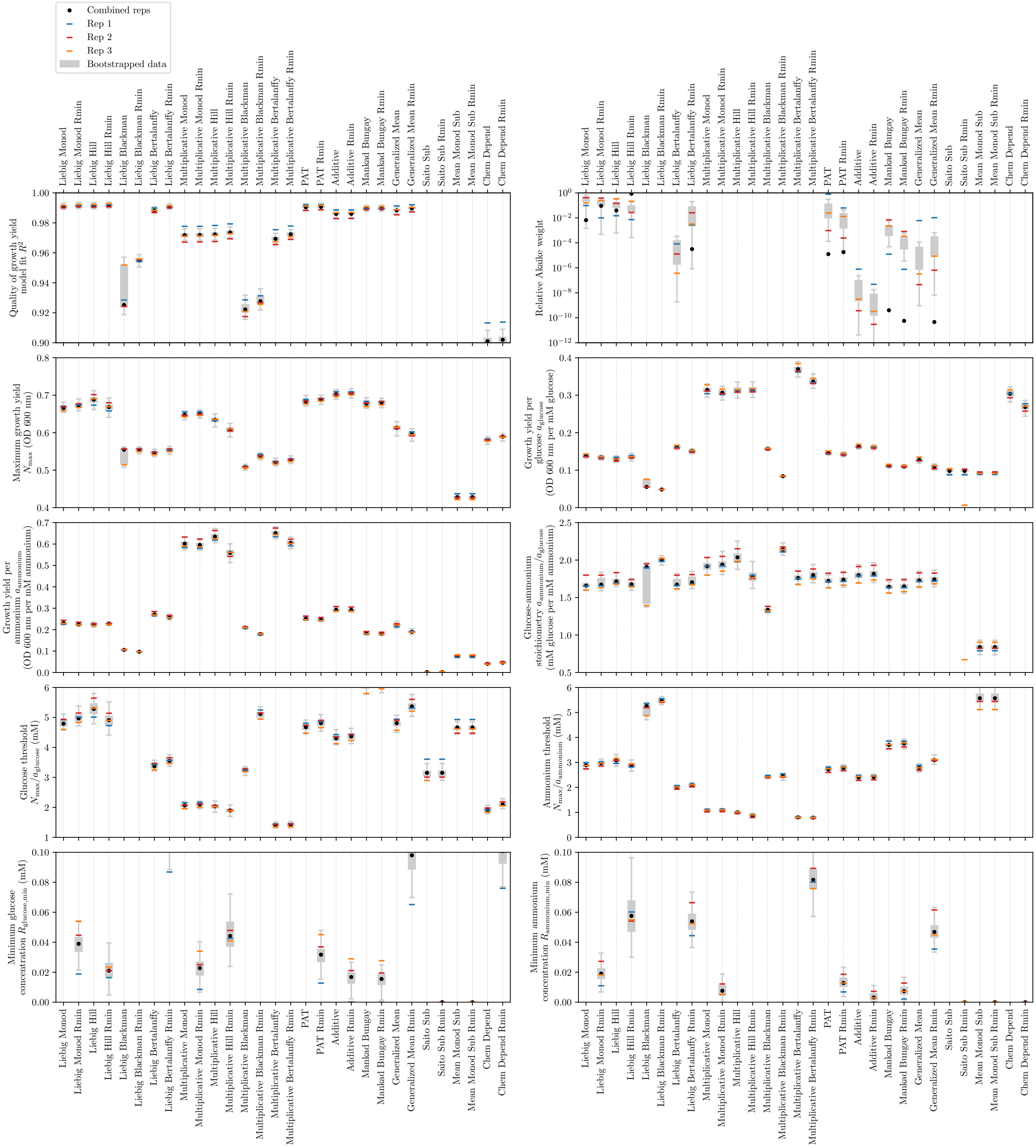


FIG. S23. **Fits of growth yield scans to models.** Same as Fig. S13 but for fits to growth yield data.

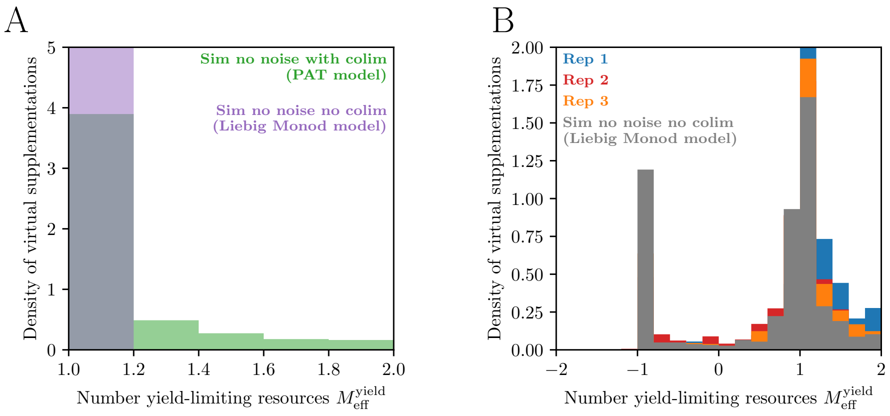


FIG. S24. **Growth yield colimitation across virtual supplementation experiments.** Same as Fig. S14 but for growth yield.

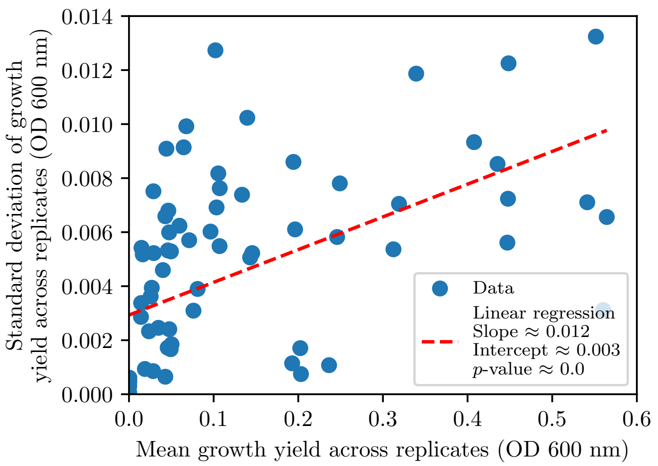


FIG. S25. **Variation of growth yield measurements across replicates.** Same as Fig. S15 but for growth yield data.

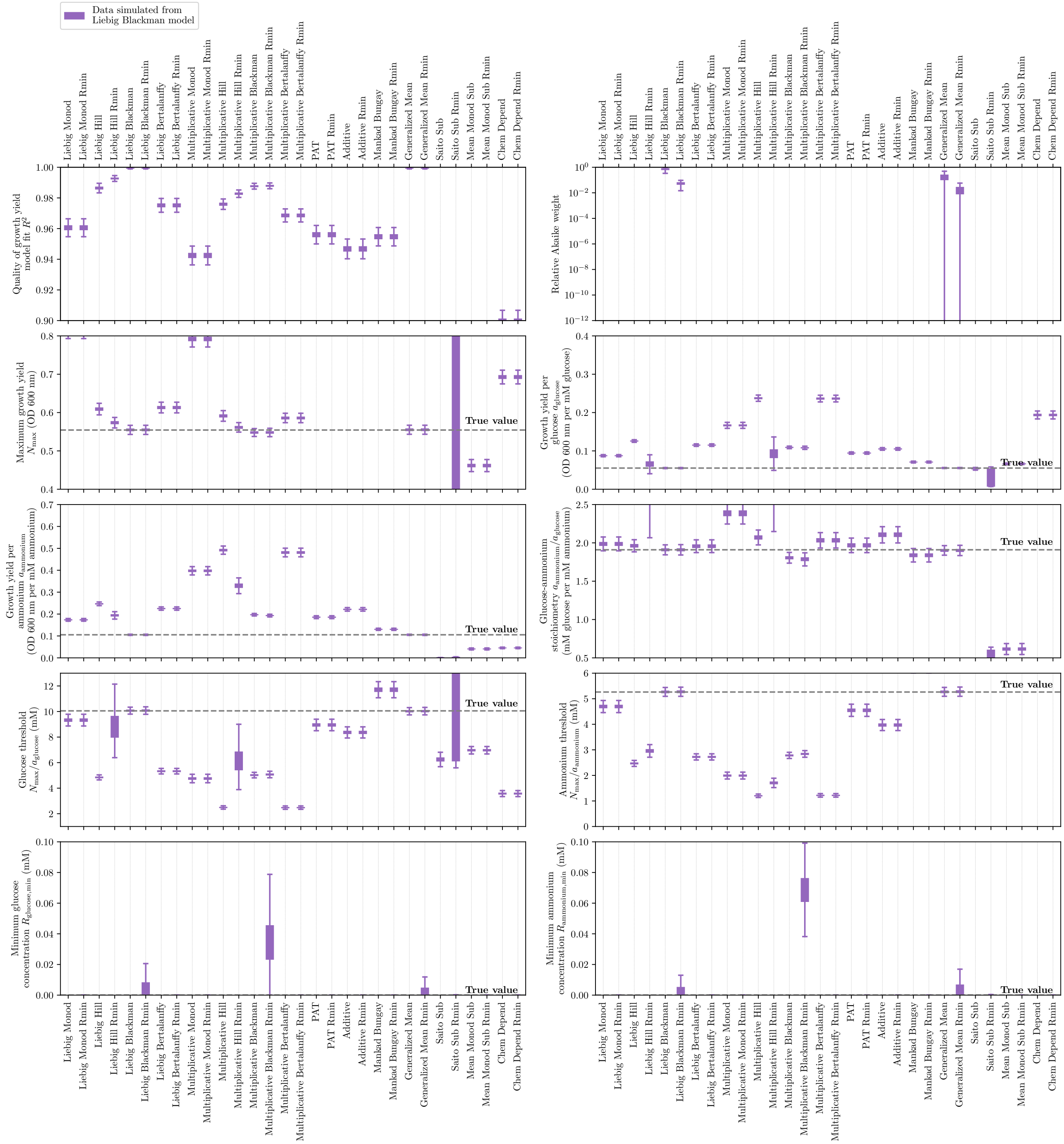

FIG. S26. **Fits of simulated (Liebig Blackman model) growth yield scans to models.** Same as Fig. S16 but for $10^4$ simulations of the Liebig Blackman model (Materials and Methods; SI Appendix, section S3) with parameters from that model's fit to the experimental growth yield data ($N_{\max} \approx 0.55$ OD, $a_{\text{glu}} \approx 0.055$ OD/mM glucose, $a_{\text{amm}} \approx 0.11$ OD/mM ammonium, $R_{\text{glu,min}} = 0$, and $R_{\text{amm,min}} = 0$).

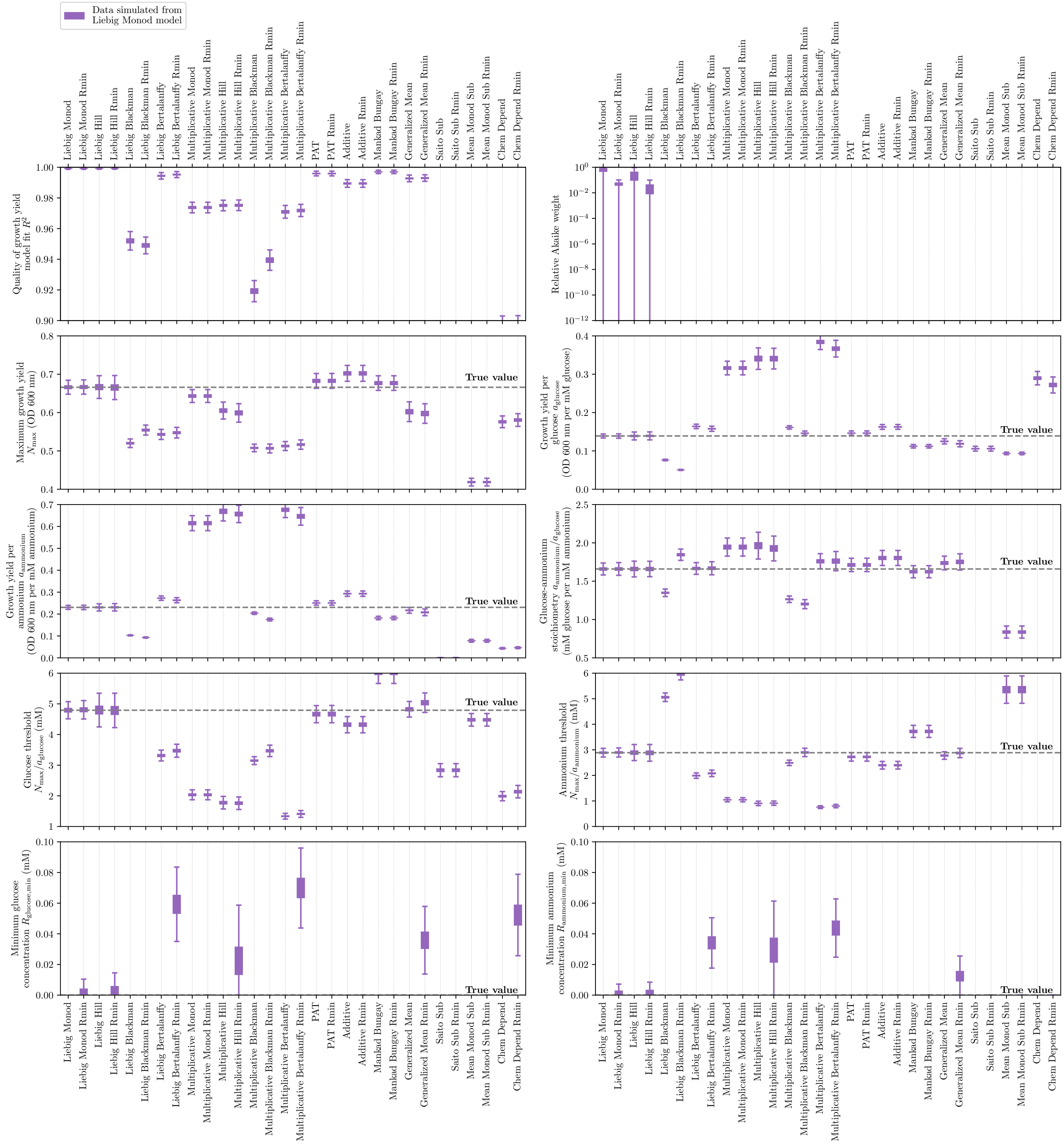


FIG. S27. **Fits of simulated (Liebig Monod model) growth yield scans to models.** Same as Fig. S16 but for $10^4$ simulations of the Liebig Monod model (Materials and Methods; SI Appendix, section S3) with parameters from that model's fit to the experimental growth yield data ($N_{\mathrm{max}} \approx 0.67$ OD, $a_{\mathrm{glu}} \approx 0.14$ OD/mM glucose, $a_{\mathrm{amm}} \approx 0.23$ OD/mM ammonium, $R_{\mathrm{glu,min}} = 0$, and $R_{\mathrm{amm,min}} = 0$).

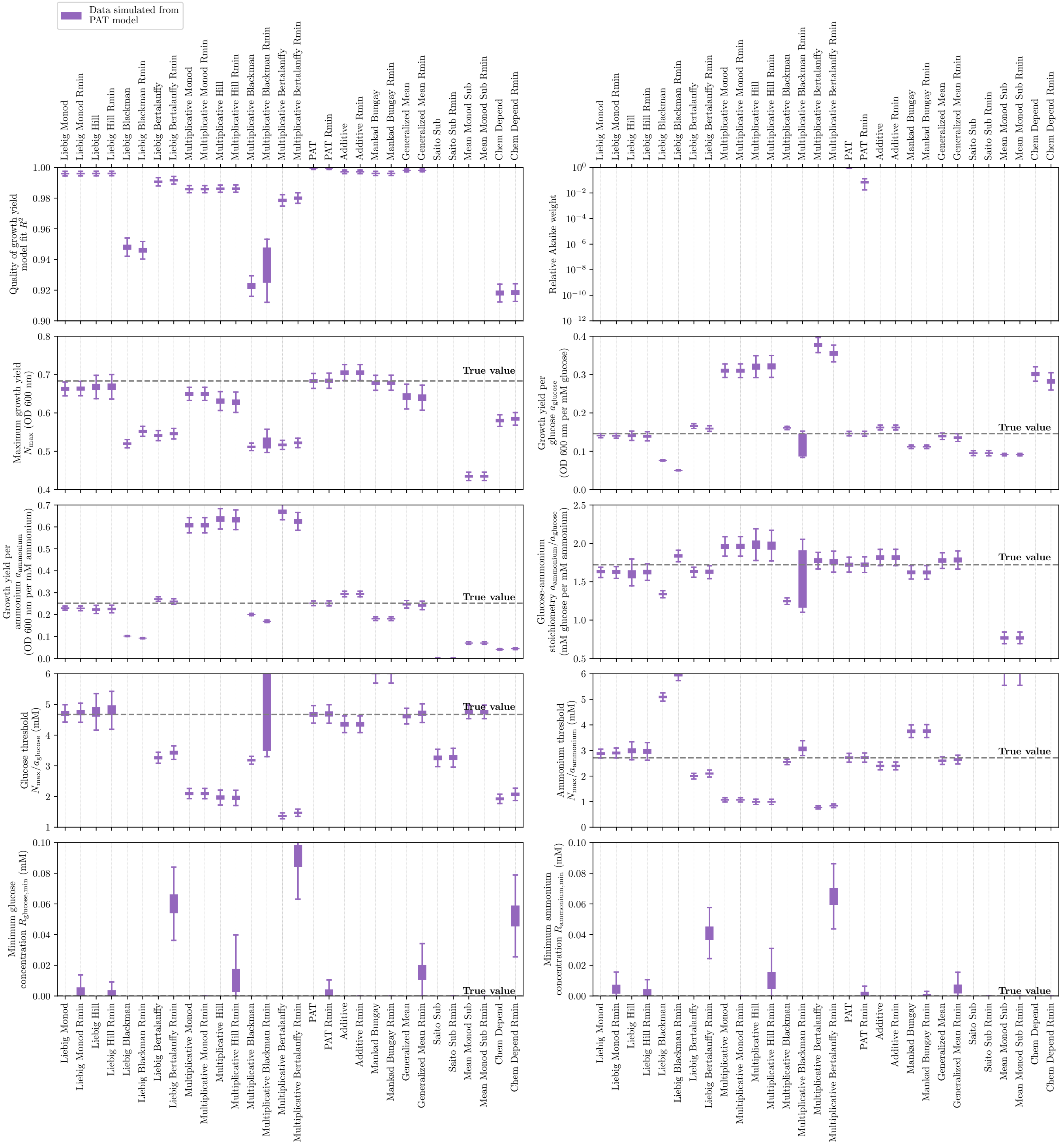

FIG. S28. **Fits of simulated (PAT model) growth yield scans to models.** Same as Fig. S16 but for $10^4$ simulations of the Poisson arrival time/synthesizing-unit model (Materials and Methods; SI Appendix, section S3) with parameters from that model's fit to the experimental growth yield data ($N_{\max} \approx 0.68$ OD, $a_{\mathrm{glu}} \approx 0.15$ OD/mM glucose, $a_{\mathrm{amm}} \approx 0.25$ OD/mM ammonium, $R_{\mathrm{glu,min}} = 0$, and $R_{\mathrm{amm,min}} = 0$).

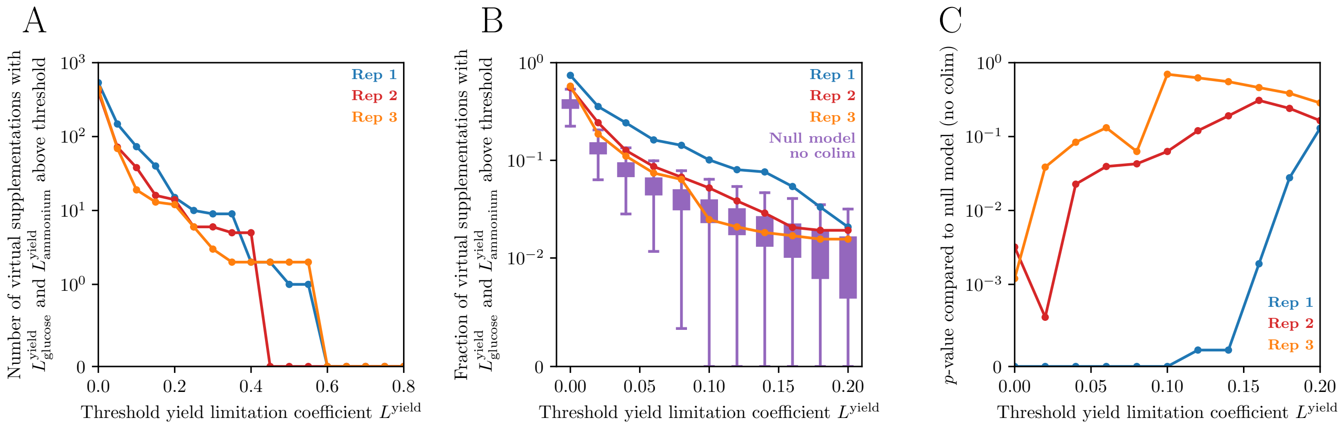


FIG. S29. **Testing significance of growth yield colimitation at different thresholds.** Same as Fig. S19 but for growth yield.

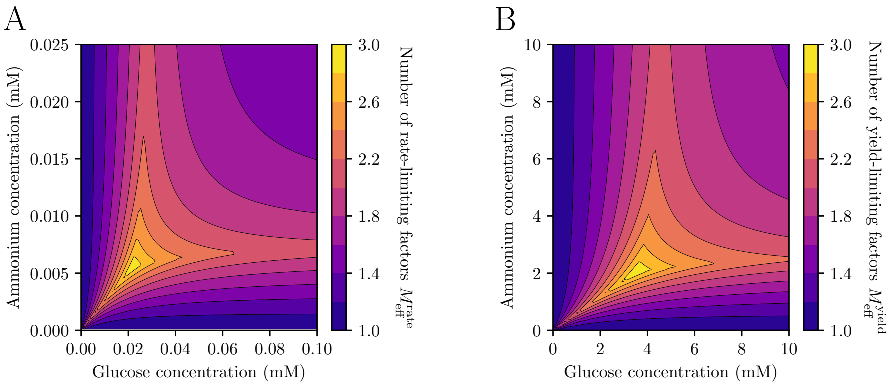


FIG. S30. **Number of limiting resources for growth rate and growth yield.** (A) The number of rate-limiting resources as a function of glucose and ammonium concentrations, implied by the model fit to the growth rate scan data in Fig. 2A (Poisson arrival time/synthesizing-unit model with $g_{\text{max}} \approx 0.66$ per hour, $s_{\text{glu}} \approx 23$ per hour per mM glucose, $s_{\text{amm}} \approx 90$ per hour per mM ammonium; $R_{\text{glu,min}} \approx 0.023$ mM, $R_{\text{amm,min}} \approx 0.0039$ mM; Materials and Methods; Dataset S1). (B) The number of yield-limiting resources as a function of glucose and ammonium concentrations, implied by the model fit to the growth yield scan data in Fig. S20A (Poisson arrival time/synthesizing-unit model with $N_{\text{max}} \approx 0.68$ OD, $a_{\text{glu}} \approx 0.15$ OD/mM glucose, and $a_{\text{amm}} \approx 0.25$ OD/mM ammonium; Materials and Methods; Dataset S2).

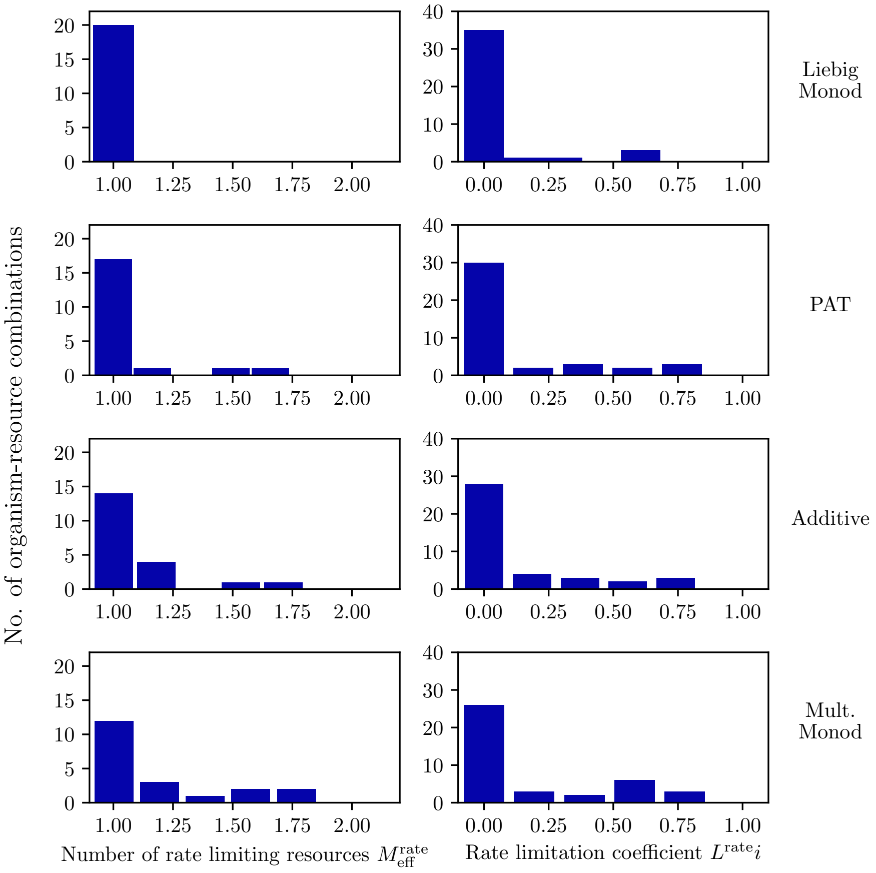


FIG. S31. **Histograms of number of rate-limiting resources and rate limitation coefficients for collected organism-resource combinations.** Same data as in Fig. 4B,C but plotted as histograms across each example growth rate model.
